# Atlas of Fshr Expression from Novel Reporter Mice

**DOI:** 10.1101/2023.11.05.565707

**Authors:** Hong-Qian Chen, Hui-Qing Fang, Jin-Tao Liu, Shi-Yu Chang, Wen-huan Chai, Li-Ben Cheng, Ming-Xin Sun, Zhi-wei Yang, Jian-Rui Feng, Ze-Min Liu, Xiao-Li Li, Yong-Hong Zhang, Clifford Rosen, Peng Liu

**Affiliations:** Laboratory of Bone and Adipose Biology Shanxi Medical University Taiyuan, Shanxi, China; Department of Dentistry The 980; Hospital of the PLA Joint Logistic Support Force Shijiazhuang, Hebei, China; Department of Orthopedic Surgery The Second Hospital University Shanxi Medical University Taiyuan, Shanxi, China; Maine Medical Center Research Institute Scarborough ME 04074, USA

**Author notes:** The corresponding author: Peng Liu, M.D., Ph.D., Laboratory of Bone and Adipose Biology Shanxi Medical University, Taiyuan, Shanxi China. Equal contributions to this work (Hong-Qian Chen, Hui-Qing Fang).

**Keywords:** Fsh, Fshr, Gene Expression, Leydig cells, and GFP Reporter

## Abstract

The FSH-FSHR signaling pathway has traditionally been considered an essential regulator in reproductive development and fertility. But there has been emerging evidence of FSHR expression in extragonadal tissues/organs. This poses new questions and long-term debates regarding the physiological role of the FSH-FSHR pathway, and underscores the need for reliable, in vivo analysis of FSHR expression in animal models. However, conventional methods have proven insufficient for examining FSHR expression due to limitations, such as the scarcity of ‘reliable’ antibodies, rapid turnover/degradation of transcripts, and a lack of robust in vivo tools. To address this challenge, we developed Fshr-ZsGreen ‘knockin’ reporter mice under the control of Fshr endogenous promoter using CRISPR/Cas9 genome-editing technology to append a P2A-ZsGreen targeting vector into a locus between the last exon and the stop codon of Fshr. With this novel genetic tool, we provide a reliable readout of Fshr expression at single-cell resolution level in vivo and in real time. Reporter animals were also subjected to additional analyses, including immunohistochemical staining, ddRT-PCR, and *in situ* hybridization, to define the accurate expression profile of FSHR in gonadal and extragonadal organs/tissues. Our compelling results not only demonstrated Fshr expression in intragonadal tissues but also, strikingly, unveiled notably increased expression in Leydig cells, osteoblast lineage cells, endothelial cells in vascular structures, and epithelial cells in bronchi of the lung and renal tubes. The genetic decoding of the widespread pattern of Fshr expression highlights its physiological relevance beyond reproduction and fertility, and opens new avenues for therapeutic options for age-related disorders of the bones, lungs, kidneys, and hearts, among other tissues/organs. Exploiting the power of the Fshr knockin reporter animals, this report provides the first comprehensive genetic record of the spatial distribution of FSHR expression, correcting a long-term misconception about Fshr expression and offering prospects for extensive exploration of FSH-FSHR biology.

## Introduction

Follicle-stimulating hormone (FSH), secreted by anterior pituitary gonadotrophs, is recognized as a crucial regulator of male and female gonadal function; indeed, reproductive biology textbooks strongly emphasize its role in normal physiology ^1^. FSH exerts its effects via a specific receptor, follicle-stimulating hormone receptor (FSHR), which belongs to the highly conserved family of G-protein-coupled receptors ^2^. Traditional views hold that in females, FSHR is expressed in granulosa cells and controls the maturation of Graafian follicles, granulosa cell proliferation, and estrogen production ^1^, while in males, FSHR is expressed in testicular Sertoli cells and regulates their metabolic functions, which are essential for proper spermatogenesis and germ cell survival ^3^. The large FSHR gene is located on chromosome 2p21 and comprises 10 exons ^4^.

However, accumulating evidence demonstrates that FSHR is also expressed in extragonadal tissues, such as endothelium ^5^, monocytes ^6^, developing placenta ^7^, endometrium ^8^, malignant tissues ^9^, bone, adipose and neural cells ^10^. Intriguingly, blocking the interaction of Fsh with Fshr mitigates some degenerative disorders in mice, such as low bone mass ^11^, obesity ^12^, and neurocognitive decline ^13^. Fshr was reported to be expressed in osteoclasts *in vitro*, and global Fsh or Fshr knockout resulted in an increase in bone mass by inhibiting bone resorption ^11^. Blocking the Fsh-Fshr interaction with either Fsh polyclonal or monoclonal specific antibodies triggered thermogenesis in adipose tissues, significantly reduced body weights ^12^, and reduced Alzheimer’s symptoms in mice ^13^. Fshr expression is also found in the vasculature of tissues ^7^ and is particularly high in solid tumors ^14^. Nevertheless, the relevance of its extragonadal expression and functions have been debated ^2,15^ Therefore, precisely defining the localization of Fshr expression remains an imperative challenge in FSH-FSHR biology due to the concern of the ‘non-specificity’ of available antibodies used to localize FSHR expression and the ‘quick turnover’ and ‘rapid degradation’ of Fshr transcripts ^2,10,16^. Thus, a suitable genetic approach to resolve this issue is warranted ^17^.

To address this challenge, we can utilize a GFP reporter driven by the endogenous promoter of Fshr, which enables the visualization of the expression of low-abundance transcripts in more accurate and context-specific ways ^18,19^, when other approaches, e.g., antibodies and RT-PCR, are limited in their ability to detect Fshr expression. Therefore, in this study, we employed CRISPR/Cas9-mediated technology to create a novel Fshr-ZsGreen reporter mouse model for precisely clarifying Fsh expression. We believe that this powerful approach, as a reliable readout of Fshr expression at the single-cell level, should allow us to accurately capture Fshr expression in real time in a tissue-specific manner. We also used other techniques to ensure the accuracy and reliability of the results. Our results challenge the current understanding of Fshr expression and refine that Fshr has a wider expression profile than previously thought. The findings of Fshr in reproductive and nonreproductive tissues/organs will provide significant insights into new roles of Fsh and Fshr in physiology and pathology and have implications for the development of new therapies for reproductive and nonreproductive disorders, particularly metabolic diseases and degenerative diseases.

## Methods

### 1. Generation of the CRISPR/Cas9-mediated Fshr-ZsGreen knockin reporter line

Fshr-P2A-ZsGreen knockin reporter mice were generated by a CRISPR/Cas9-based approach. Briefly, one sgRNA was designed by the CRISPR design tool (http://www.sanger.ac.uk/) to target the region of the stop codon in the transcript NM_013523.3 exon 10 of mouse Fshr ^20^, and then was screened for on-target activity using a Universal CRISPR Activity Assay [UCATM, Biocytogen Pharmaceuticals (Beijing) Co., Ltd]. The targeting vector containing P2A-ZsGreen and 2 homology arms of left (1378 bp) and right (1493 bp) each was used as a template to repair the DSBs generated by Cas9/sgRNA. P2A-ZsGreen was precisely inserted before the stop codon of the Fshr locus. The T7 promoter sequence was added to the Cas9 or sgRNA template by PCR amplification in vitro. Cas9 mRNA, sgRNA and the targeting vector were co-injected into the cytoplasm of one-cell-stage fertilized C57BL/6J eggs. The injected zygotes were transferred into oviducts of Kunming pseudopregnant females to generate F0 mice. F0 mice with the expected genotype as confirmed by tail genomic DNA PCR, DNA sequencing and Southern blotting were mated with C57BL/6J mice to establish germline transmitted F1 heterozygous mice. F1 heterozygous mice were further genotyped by tail genomic PCR, DNA sequencing and Southern blotting. Primer sequences for genotyping F0 and F1 are described in Supplementary Material 1.

The produced Fshr-ZsGreen (Fshr-ZsG) knockin mice were maintained as heterozygotes, and homozygotes were used for experiments. For genotyping, genomic DNA was extracted from tail tips and assayed using polymerase chain reaction (PCR) primer sets for the Fshr-ZsGreen allele. The primer sequences for genotyping are described in Supplementary Materials 1. All mice were maintained on a 12 h light/dark cycle with food and water *ad libitum*. The care and treatment of animals in all procedures strictly followed the NIH Guide for the Care and Use of Laboratory Animals. The animal protocols used in this study were approved by the Shanxi Medical University IACUC committee.

### 2. Tissue harvest and preparation

Mice were terminated by CO_2_, and organs were harvested and fixed with 4% paraformaldehyde (PFA) for 12 to 24 hours at 4°C. For bone samples, decalcification was performed with daily changes of EDTA solution (0.5 M, pH 8) for 7 days. After fixation and decalcification processes, samples were transferred to 15% and 30% sucrose overnight, respectively. The tissues were then embedded with optimal cutting temperature (OCT) compound and stored at -80°C. The embedded samples were sectioned into 5- to 25-μm-thick sections using a Leica cryostat CM1950.

### 3. Immunofluorescence staining

Immunofluorescence staining was performed as described previously ^21^. Briefly, air-dried 7- to 25-μm-thick frozen sections were washed with 1×PBS for 10 minutes three to five times, followed by blocking with 10% BSA for 20 minutes at room temperature, and the samples were permeabilized with 0.5% Triton X-100 in 1×PBS for 10 minutes. Then, diluted primary antibodies were added to the slides. After overnight incubation with antibodies at 4°C, the slides were washed with 1×PBS for 15 minutes three to five times and then incubated with secondary antibody conjugated with fluorescence for 30 minutes at room temperature. The slides were washed again thoroughly with 1×PBS for 10 minutes three times. The slides were then stained with DAPI, rinsed with 1×PBS three times for 5 minutes each, and mounted with 50% glycerol for confocal imaging. Rabbit IgG was used as the negative control. Primary and secondary antibodies were purchased from Servicebio (Wuhan Servicebio Technology Co., Wuhan, China), except when indicated otherwise. The primary antibodies are listed in Supplementary Materials 2. In addition, sections of different tissues/organs derived from C57BL/6J (B6) mice were used as negative controls for Fshr-ZsGreen expression. A donkey anti-rabbit IgG conjugated with Cy3 was employed at a 1:400 dilution as the secondary antibody (Servicebio, Wuhan, China).

### 4. Single RNA-fluorescence in situ hybridization (RNA-smFISH)

RNA in situ hybridization, the gold standard method for visualizing RNA expression and localization in cells, tissue sections, and whole organs, was performed on tissue sections as described previously ^22^. Briefly, frozen tissue sections of the testis from Fshr-ZsGreen mice were washed with 1XPBS three times. The sections were then treated with proteinase K (20 μg/ml) for 5 minutes at 37°C to permeabilize the cells and allow probe penetration, followed by three washes with 1XPBS. Specific oligonucleotide sense and antisense DNA probes for Fshr were designed and synthesized by the manufacturer (GeneCreate Biological Engineering Co., Ltd., Wuhan, China). The probes were labeled with a fluorescent dye (e.g. Cy3) for visualization. RNA-smFISH was performed on tissue sections using a commercially available kit (e.g., SureFISH, Agilent Technologies) at 37 ° C for 2 hours. After hybridization, the sections were washed with SSC solution (2xSSC, 37°C for 10 mins, 1xSSC, 37°C for 5 mins twice, 0.5xSSC for 10 mins) to remove unbound probes, and counterstained with a nuclear stain (e.g., DAPI) to visualize the tissue architecture. The sections were imaged using a fluorescence microscope equipped with appropriate filter sets. A sense probe was used as a negative control to ensure specificity and sensitivity. The sequences of sense and antisense probes are described in Supplementary Materials 2.

### 6. Imaging

Imaging of the slides was carried out as described before ^21^. Briefly, the fluorescence images of the frozen sections were obtained using a Nikon A1 HD25 confocal microscope with a DUVB detector and plan Apo λ 4×, plan Apo VC 20× DIC N2, plan Apo λ 40×, and plan Apo λ 100×C oil objectives, illuminated with a wavelength of 405, 488, or 561 nm to excite DAPI, GFP, or Cy3, respectively; detection was performed with a 425–475, 500–550, or 570–620 nm bandpass filter. To assess the number of cells in each field of view, tissue-cleared images were converted from 3D to MAIP (maximum projection of the Z-stack across the whole section). Data were acquired with NIS- Elements AR 5.20.00 64-bit software. Structured illumination microscope (SIM) images were obtained and processed with N-SIM-S super-resolution model (Nikon) and Image J-Colocalization Finder.

### 7. Droplet digital RT‒PCR (ddRT-PCR)

Tissues were harvested from 10-week-old B6 mice. Samples were dissected free of connective tissue and homogenized with a fast multi-sample tissue cryogenic grinder (LC-FG-96, Lichen Instrument Technology Co., Ltd., Shanghai, China), and total RNA was extracted using NucleoZOL (NucleoZOL; Macherey-Nagel GmbH & Co., KG, Dylan, Germany). mRNA was reverse transcribed using an M5 Super qPCR RT kit with gDNA remover (MF012-T, Mei5bio, Beijing). Droplet digital PCR was performed as described previously ^21^. The primers used for ddRT-PCR were as follows: mFshr Fwd-5’ ccgcagggacttcttcgtcc-3’; mFshr Rev-5’-ttggtgactctgggagccga-3’.

### 8. Cell cultures of Leydig cell line TM3

TM3 cells were purchased from Pricella (Wuhan Pricella Biotechnology Co., Ltd., Wuhan, China) and subcultured with DMEM/F12 +5% HS +2.5% FBS +1% P/S with the company’s instruction in T25 flasks for extraction of total RNA and on coverslips for immunofluorescence staining of Fshr expression.

## Results

To precisely define Fshr expression in mice, we utilized several complementary strategies, including Fshr endogenous promoter driving ZsGreen knockin reporter mice, immunofluorescence staining with antibodies against tissue/cell type-specific markers and Fshr, ddRT-PCR and RNA-smFISH, to comprehensively examine Fshr expression.

### 1. Generation of Fshr-ZsGreen knockin reporter mice

CRISPR/Cas9-mediated homologous recombination was used to generate embryonic stem cell (ESC) clones, in which a P2A-ZsGreen cassette was precisely inserted before the stop codon of the Fshr gene followed by the 3′ UTR of the Fshr allele, as described in the Methods (Fig. 1A). This P2A-ZsGreen expression cassette under the control of endogenous Fshr regulatory elements ultimately generates ZsGreen proteins without disrupting Fshr expression. The 19 amino acids P2A sequences functions to cause ribosomal “ skipping” during translation, resulting in a missing peptide bond and effectively separates the two proteins, e.g. Fshr and ZsGreen ^23,24,25^. The injected zygotes were transferred into oviducts of Kunming pseudopregnant females to generate F0 mice. F0 mice with the expected genotype were confirmed by tail genomic DNA PCR, DNA sequencing and Southern blotting (Fig. 1B and C) and then mated with C57BL/6J mice to establish F1 heterozygous mice with the germline-transmitted transgene. F1 heterozygous mice were further genotyped by tail genomic PCR, DNA sequencing and Southern blotting (Fig. 1B and C). The results from these tail genomic PCR, DNA sequencing and Southern blots in both F0 and F1 pups demonstrated that the targeted P2A-ZsGreen cassette was accurately inserted into the designed site between exon 10 and the stop codon of the Fshr gene before the 3’ UTR. We maintained heterogeneous Fshr-ZsGreen mice and used homogeneous mice for experiments. These mice were genotyped using primer sets specific to P2A-ZsGreen, as shown in Figure 1D. Fshr and GFP are transcribed in a bicistronic mRNA but subsequently translated into two independent proteins rather than as a fusion protein. This design ensures unaffected Gαi3 transcription and function and simultaneous GFP expression as a reporter protein. All animals were fertile and showed normal behavior and no obvious abnormal phenotypes.

**Figure 1.**
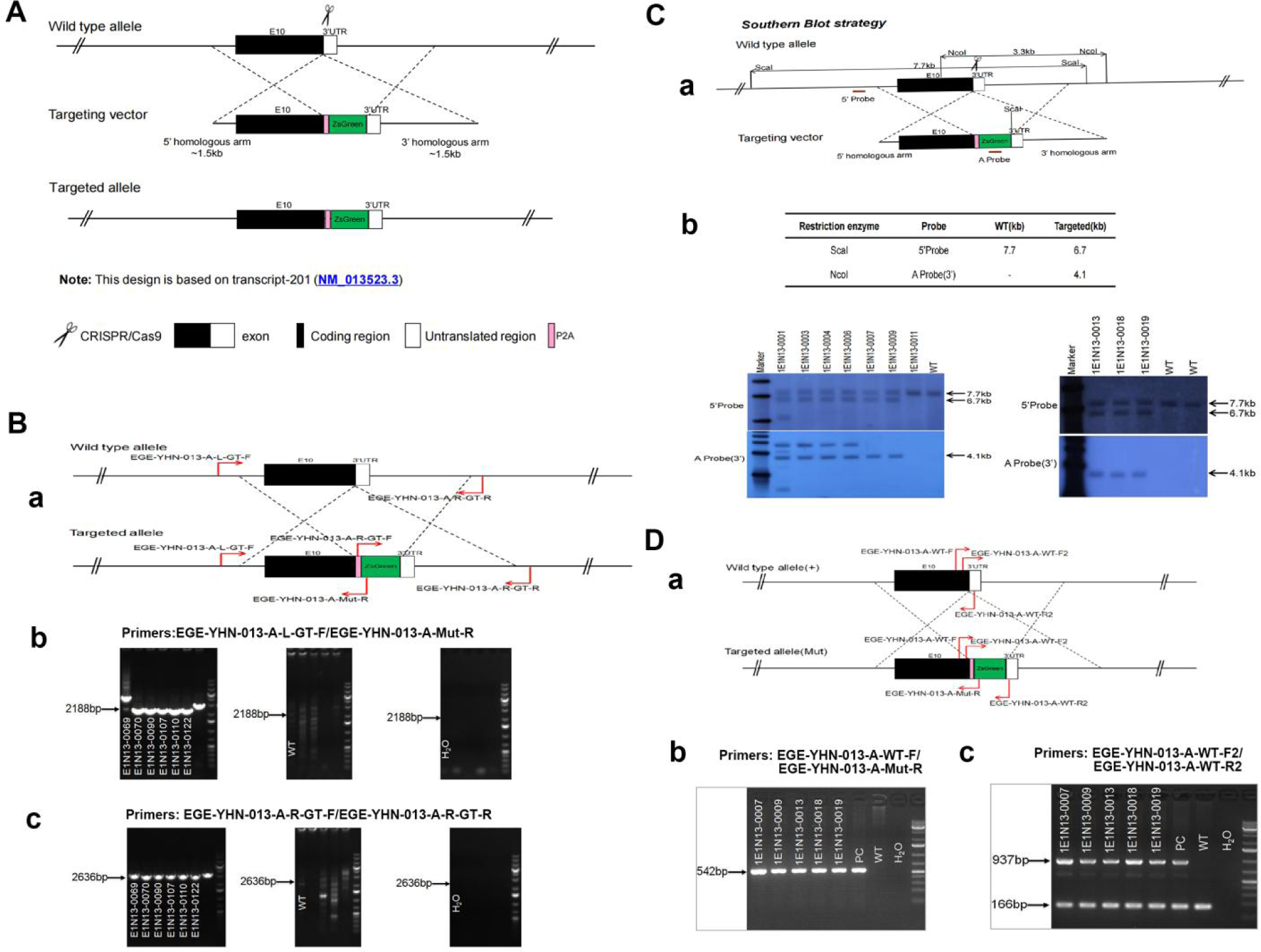
Generation of CRISPR/Cas9-mediated Fshr-ZsGreen knockin reporter mice. A. CRISPR/Cas9-mediated targeting strategy to generate Fshr-P2A-ZsGreen knockin mice. **B**. Detection of integration by PCR in F0 and F1 mice: (a) Schematic of PCR primer design specific to Fshr-P2A-ZsGreen and the wild-type allele. The results of genomic DNA PCR genotypes using the primer pair EGE-YHN-013-A-L-GT-F/EGE-YHN- 013-A-Mut-R (b) and another primer pair EGE-YHN-013-A-R-GT-F/EGE-YHN-013-A-R- GT-R (c). **C**. Southern blot confirmation of the correct integration of the P2A-ZsGreen allele in F0 and F1 mice. The Southern blot results demonstrated the successful generation of the targeted P2A-Fshr-ZsGreen allele: (a) Restriction sites in the wild-type sequence and targeted vector; (b) Southern blot analysis probes, expected restriction fragment lengths as indicated and blotted images. **D**. The 2^nd^ PCR genotyping strategy using primers: (a) schematic for the design of PCR primer sets for the Fshr-ZsGreen allele; (b and c) genotyping readout of heterozygous mice;

### 2. Examination of Fshr expression in Fshr-ZsGreen reporter mice

With confirmation that the Fshr-P2A-ZsGreen targeting vector was successfully inserted into the Fshr locus, we investigated Fshr expression by confocal microscopy to detect ZsGreen and immunostaining for tissue/cell markers in frozen sections of tissues/organs of Fshr-ZsGreen reporter mice. To ensure that there was no nonspecific expression of Fshr-ZsGreen in the examined tissues/organs, we took frozen sections derived from wild-type mice (B6) as negative controls. The negative controls were imaged under the conditions used for examining Fshr-ZsGreen expression. The representative results are shown in Supplementary Data 1, showing no nonspecific expression of Fshr-ZsGreen in the negative controls. On this basis, we performed the following imaging to examine Fshr-ZsGreen expression in the major organs and tissues.

### 1) Reproductive organs

As the reproductive system is well known to express Fshr, we first tested Fshr- ZsGreen expression in the ovary and testis to ensure ZsGreen expression driven by the endogenous Fshr promoter. In the ovary, we observed Fshr-ZsGreen expression in the different stages of follicles from primordial follicles to primary follicles, secondary follicles, and the corpus luteum (Fig. 2A-b and e). In the ovarian/Graafian follicles, expression was observed in the oocytes, granulosa cells/follicle cells and theca (interna and externa) (Fig. 2A-b, c, e and f). We also found Fshr-ZsGreen expression in the ciliated epithelial cells in the oviduct (Fig. 2A-d and h). Furthermore, we employed an antibody against Stra8 ^26^ to perform immunofluorescence staining to identify reproductive cells and observed the colocalization of Fshr-ZsGreen with Stra8 staining as a marker for reproductive cells (Fig. 2A).

**Figure 2.**
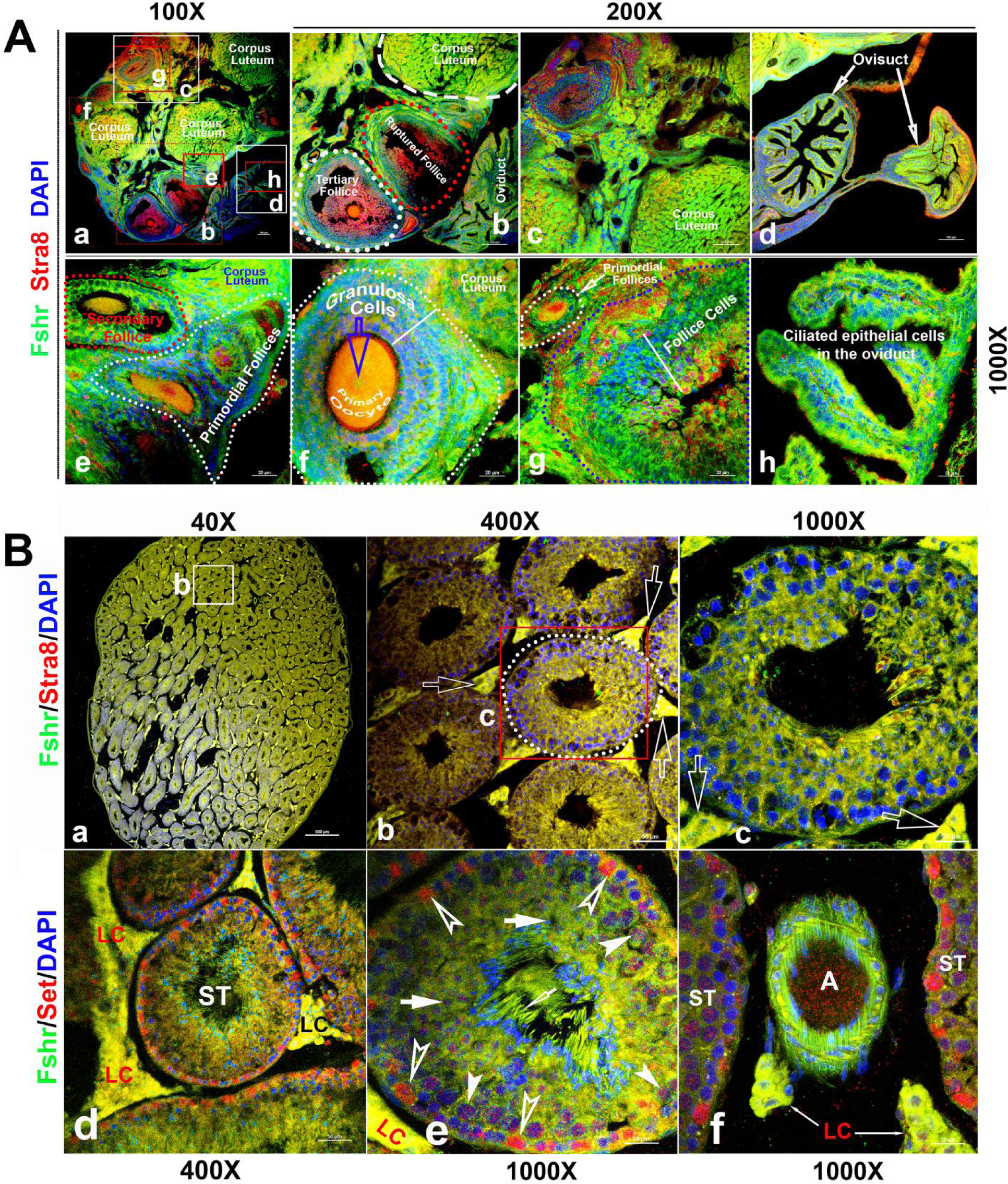
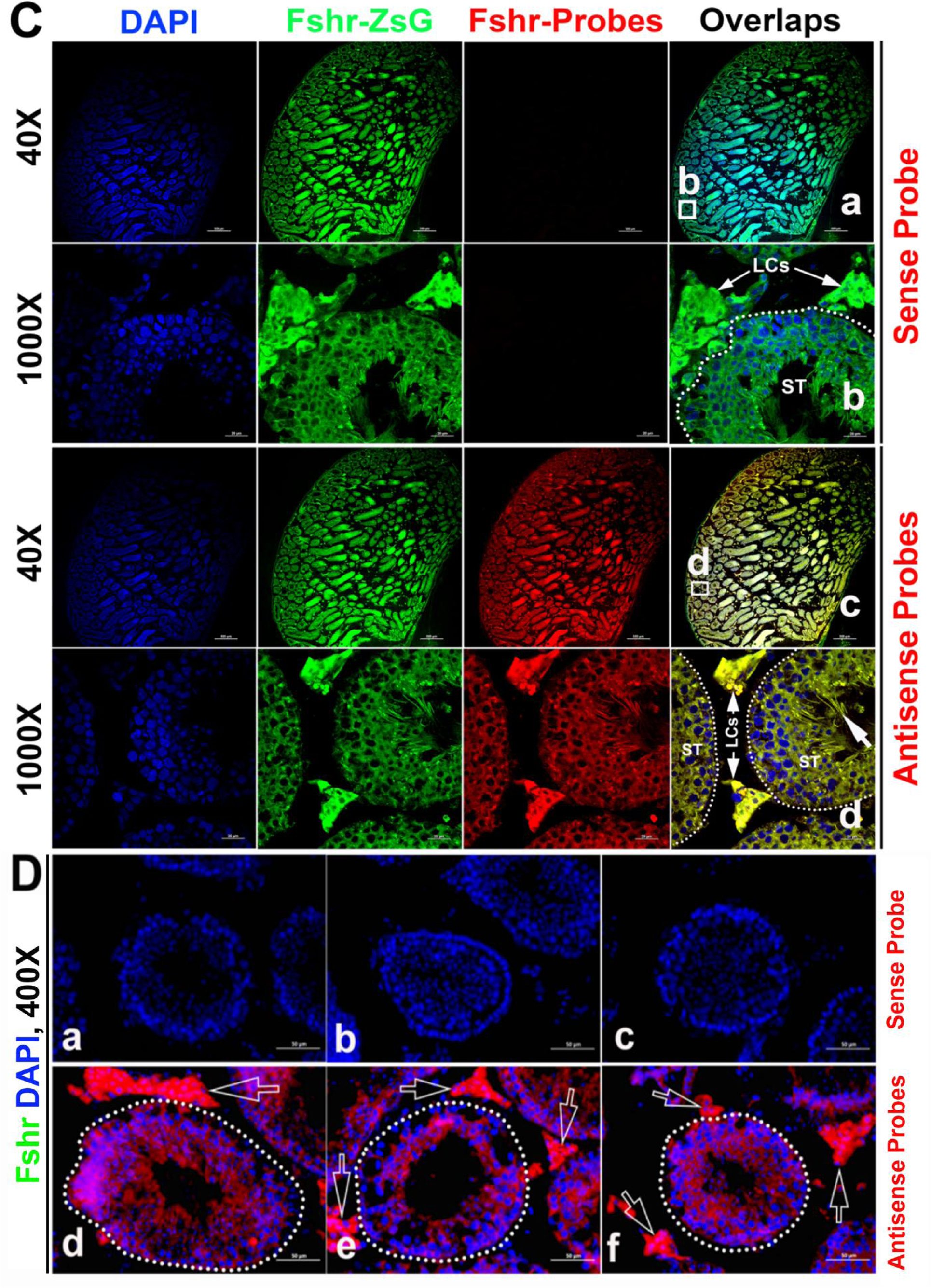
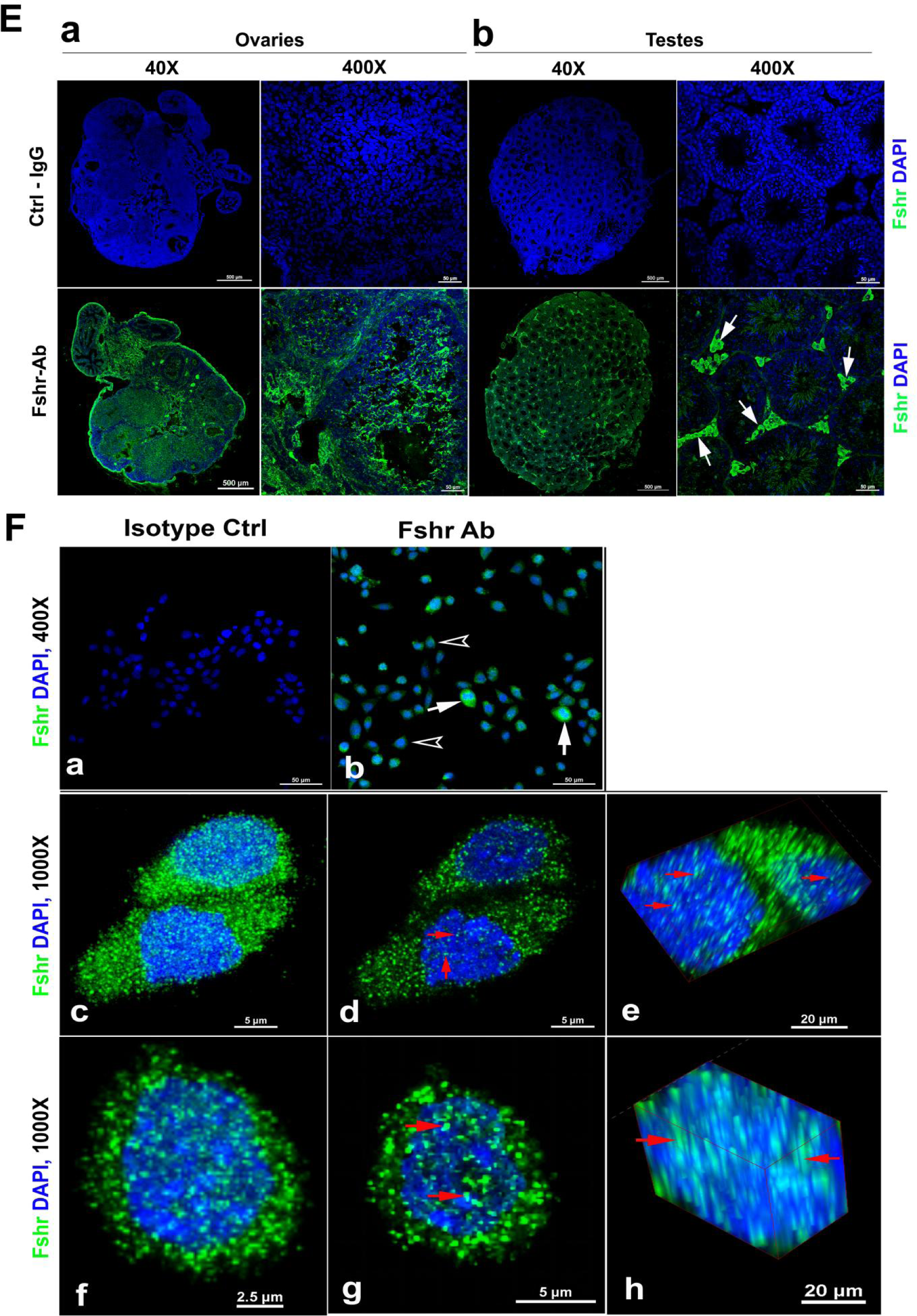
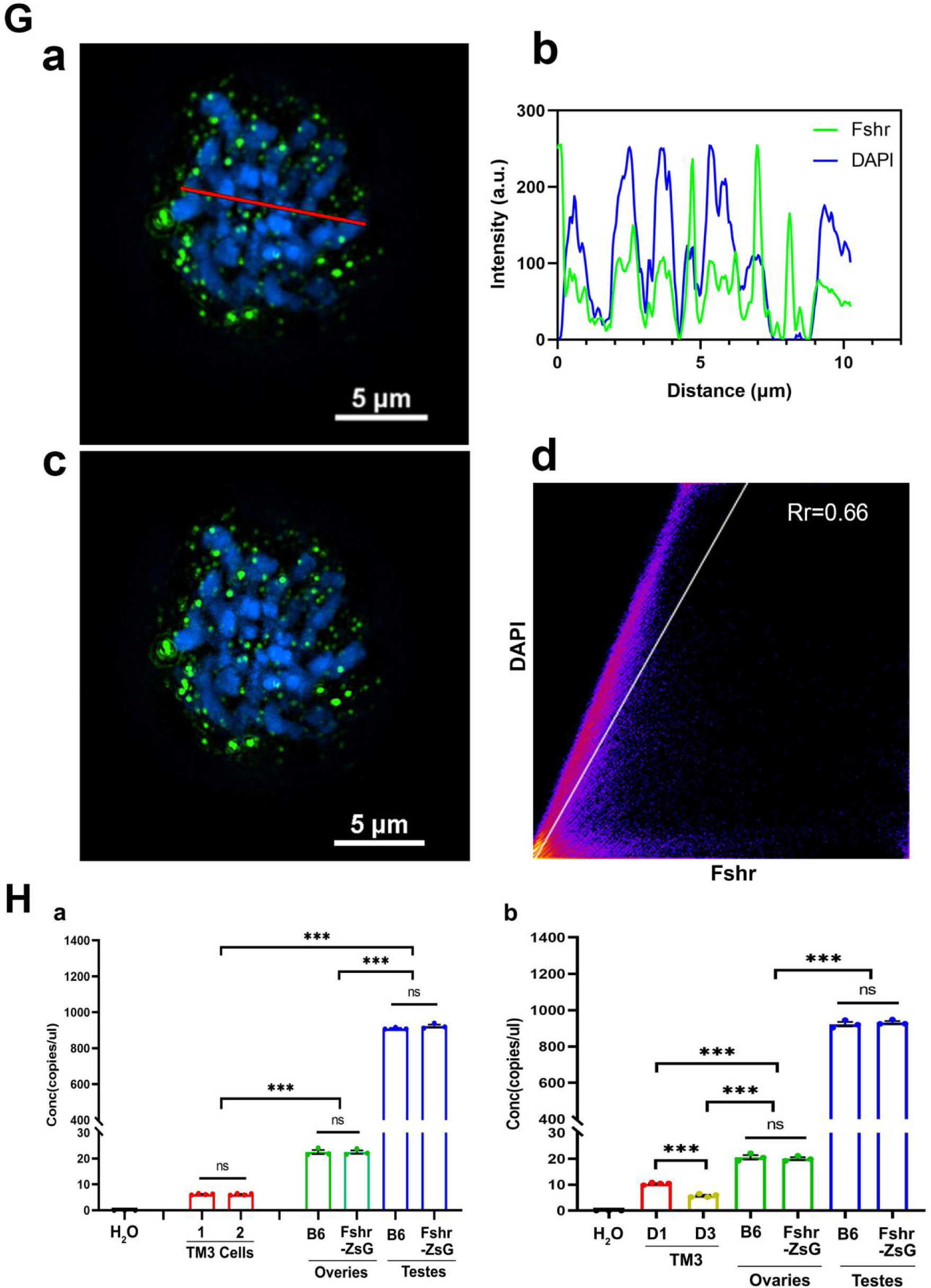
Imaging of Fshr-ZsGreen expression in the reproductive system. A. Fshr expression in the ovary. Frozen sections of the ovary were immunostained with an antibody against mouse Stra 8, followed by imaging of Fshr-ZsGreen expression and its colocalization with Stra8 staining. The entire picture of the sectioned ovary is shown in A-a. The representative images of the section were taken at 400X magnification. A-b. an area containing a corpus luteum, tertiary and ruptured follicles and a partial oviduct; A-c. an area with more corpora lutea; A-d. oviducts. Furthermore, representative images at 1000X magnification; A-e. primordial and secondary follicles; A-f. a mature follicle; A-g. follicle cells in a corpus luteum; and A-h. ciliated epithelial cells in the oviduct. Scale bars: 100 μm for a, b, c and d, 20 μm for e to h. B. Fshr-ZsGreen expression and its colocalization with Stra8 (a to c) and Set (d to f) in the testis. B-a, the whole image of the sectioned testis; B-b, the representative area of seminiferous tubes with interstitial cells (Leydig cells); and B-c, an seminiferous tube. White empty arrowheads indicate Leydig cells; B-d, the representative area of seminiferous tubes with interstitial cells (Leydig cells) stained for Set; B-e, an seminiferous tube stained for Set and an artery with partial areas of seminiferous tubes stained for Set. Magnifications: 40X for a, 400X for b and d, and 1000X for c, e and f . Scale bars: 500 μm for b and d, 50 μm for c and d, and 20 μm for c, e and f. C. RNA-smFISH confirmation of Fshr expression in Fshr-ZsG mice. Mixed anti- sense probes were applied for detection of Fshr while a sense probe was taken as a negative control. The entire image of the sectioned testis is taken at 40X magnification (a and c), and two representative areas of seminiferous tubes with Leydig cells are demonstrated at 1000X magnification (b and d). White arrows indicate spermatozoa. Abbrev.: LCs-Leydig cells, and ST-seminiferous tubule. Scale bars: 50 μm. D. RNA-smFISH confirmation of Fshr expression in B6 mice. Mixed anti-sense probes were applied for detection of Fshr while a sense probe was taken as a negative control. Three representative areas of seminiferous tubes with Leydig cells were imaged at 400X magnification (a to c for the sense probe; d to f for antisense RNA probes). White empty arrows indicate Leydig cells. Scale bars: 50 μm. E. Fshr expression in the ovary (a) and testis (b) of B6 mice. Frozen sections of the ovary and testis were immunofluorescence stained for Fshr. Representative areas of whole the ovary or the testis image are present at 400X magnification. White arrows indicate Leydig cells. F. Immunofluorescence staining for Fshr in TM3 cells. a. isotype control (IgG); b. positive staining for Fshr with mouse Fshr antibody, empty arrowheads indicate lower Fshr expression cells and white arrows indicate higher Fshr expression cells; c and f, higher magnification of Fshr positively stained cells (maximum intensity projected images); d and g, single layer images across the centers of the nuclei in c and f; e and h. 3D images of the nuclei showing Fshr located in the nuclei; G. SIM images of Fshr localized in the nuclei of TM3 cells. a. a single layer of the nuclei for analysis of Fshr colocalization of DAPI, a red line indicates a single layer for analysis of colocalization; b. fluorescence analysis of colocalization of Fshr with DAPI from a. c. the whole layers of nuclei for colocalizationof Fshr with DAPI; d. Pearson coefficient analysis of the colocation by Image J-Colocalization Finder, Rr=0.66 [PCC (Rr): 0.5 to 1 indicating colocalization]. H. Comparison of Fshr expression in TM3 cells, the testes and ovaries of Fshr- ZsGreen and B6 mice assessed by ddRT-PCR. a, the first comparison of Fshr expression among TM3 cells (cultured for 1 day); the testes and the ovaries from two types of mice, two groups of TM3 cells (1 and 2) for measuring Fshr expression (n=4 per group). b. the second comparison, TM3 cells were cultured for 1 day (D1) and 3 day (D3). Three samples for each organ (n=3 mice/each organ, aged 3 months). ***p<0.001; ns-no significant difference in the comparisons.

In the testis, we found Fshr-ZsGreen expression and its colocalization with Stra8 ^27^ staining in the cells of seminiferous tubules (STs), including primary spermatocytes, Sertoli cells and spermatids, and particularly in interstitial Leydig cells (LCs) between STs (Fig. 2B). Figure 2B-a shows an image of the whole sectioned testis. A representative ST is displayed at two magnifications (Fig. 2B-b and c), demonstrating strong expression of Fshr-ZsGreen in Leydig cells, as indicated by empty white arrows. In addition, we also applied an antibody against Set to identify testis cells, whose expression was reported in multiple cell types of the mouse testis at different developmental stages ^28^. In Figure 2C, a representative image of ST with Leydig cells is shown at a lower magnification (400X) and a higher magnification (1000X) (Fig. 2C-a and b), showing colocalization of Fshr-ZsGreen and Set staining in testis cells. We found strong Set signal in Fshr-ZsGreen-positive spermatogonia, as indicated by empty white arrowheads (Fig. 2C-b). Fshr-ZsGreen was also observed in the arterioles of the testis (Fig. 2C-c).

Because of RNA in situ hybridization as the gold standard method for visualizing RNA expression and localization in cells, tissue sections, and whole organs ^22,29^, we further carried out single RNA molecule-fluorescence in situ hybridization to confirm Fshr expression in the LCs, using antisense probe for identifying Fshr expression in the sections of Fshr-ZsGreen and B6 mice, whereas its sense probe was used as a negative control (Fig. 2C and D). Additionally, we also examined Fshr expression by immunofluorescence in Leydig cell line TM3 ^30^ and found that the majority of TM3 cells stained for lower levels of Fshr, only a few cells showed relatively higher levels of Fshr expression (Fig. 2E-a and b). Interestingly, we also noticed Fshr expression located in the nuclei, demonstrated by confocal 3D images (Fig. 2E-c-h) and SIM images (Fig. 2G- a-di) of the nuclei.

We then performed ddRT-PCR to compare Fshr expression in TM3 cells, testis, and ovaries of Fshr-ZsG and B6 mice at 3 months old (Fig 2H-a and b, two representative of four experiments). The results showed that Fshr expression was similar in the testes and ovaries of both types of mice, indicating that the insertion of P2A- ZsGreen did not disrupt Fshr expression in Fshr-ZsGreen mice. However, Fshr expression was significantly higher in the testes compared to the ovaries, by almost 40- fold. Interestingly, TM3 cells exhibited much lower Fshr expression levels than the testes and ovaries. In fact, the levels were approximately 145-fold lower than in the testes and 3.6-fold lower than in the ovaries.

Last, we also searched for FSHR expression in scRNA-seq databses (DISCO (immunesinglecell.org/genepage/FSHR), BioGPS (http://biogps.org/#goto=genereport&id=2492) and CZ CELL×GENE Discover (https://cellxgene.cziscience.com/e/535e9336-2d8d-43c3-944d-bcbebe20df8a.cxg/), showing FSHR expression in human Leydig cells (Supplementary Data 3).

Overall, we observed Fshr-ZsGreen expression in the reproductive system, demonstrating that ZsGreen is a readout for Fshr expression. In addition to its sole expression in granulosa cells and Sertoli cells, as reported previously, our findings clearly reveal that Fshr is also expressed in other cell types in the reproductive system, particularly in Leydig cells.

### 2) Skeletal tissues

Fsh has been thought to have a direct role in bone ^11^, therefore we next examined the expression pattern of Fshr in femoral sections, as shown in Figure 3A, under confocal fluorescence microscopy. The representative areas are presented at two magnifications of 400X and 1000X. In the epiphyseal growth plate, we observed lower expression of Fshr in chondrocytes, as indicated by two dotted lines, compared to its expression in cells located in the sponge area above or under the dotted lines (Fig. 3A-a). At higher magnification, Fshr was expressed in the columns of chondrocytes from the resting zone to the transformation zone (Fig. 3A-e).

**Figure 3.**
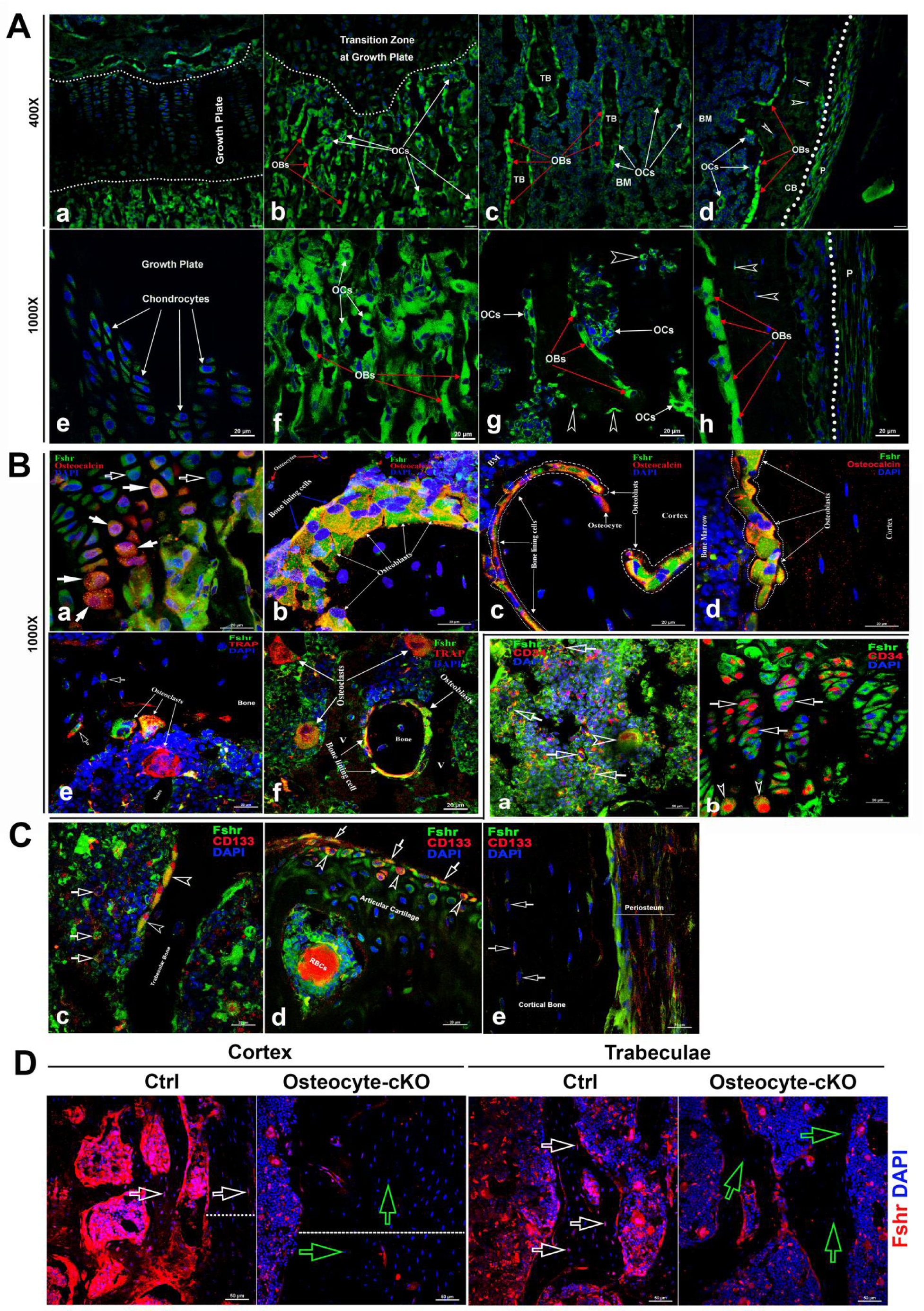
Fshr-ZsGreen expression in skeletal tissues. **A.** Detection of Fshr-ZsGreen expression in frozen sectioned skeletal tissues by fluorescence confocal microscopy. The upper panels show Fshr-ZsGreen-positive cells in the representative areas of skeletal tissues at low magnification (400X), while the lower panels show their corresponding cells at high magnification (1000X). These representative areas demonstrate the Fshr-ZsGreen-positive cells in chondrocytes of the growth plate (a and e), on the surfaces of sponge bone under the growth plate (b and f), in trabecular bone in the bone marrow (c and g), and in the cortex (d and h). Abbreviations: OBs-osteoblasts; OCs-osteoclasts; BM-bone marrow; CB-cortical bone; and P-periosteum. Arrows: white empty arrowhead indicates GFP-positive osteocytes. Magnifications: 400X for a to d and 1000X for e to h; scale bars: 50 μm for a to d and 20 μm for e to h. **B.** Confirmation of GFP-positive cell identities with antibodies against either osteocalcin or Trap as markers for osteoblasts or osteoclasts. Colocations of GFP expression with osteocalcin in mature chondrocytes (a), osteoblasts, bone lining cells and osteocytes on the surface of or within sponge bone (b), trabecular bone (c), and cortical bone (d). Colocations of Trap with multinucleated GFP-positive cells are shown in e and f, which represent osteoclasts on the resorptive areas and in bone marrow adjacent to bone surfaces and bone lining cells over trabecular bone. Arrows: white empty arrows indicate early chondrocytes with GFP expression but undetectable osteocalcin expression, while white arrows point to mature chondrocytes with both strong GFP and osteocalcin expression. Magnifications: 1000X for a to f. Scale bars: 20 μm for a to f. **C.** Identification of stem/progenitor cells. a. Colocalization of Fshr-ZsGreen with CD34-positive staining in bone marrow, indicated by empty white arrows; an empty white arrowhead indicates multinucleated Fshr-ZsGreen cells with CD34-positive staining. b. CD34-positive stained cells in growth plates. Empty white arrows indicate spindle- shaped ZsGreen-positive chondrocytes stained positively for CD34, where empty white arrowheads point to round cells with CD34-positive staining located in the bottom of the growth plate. c. Colocalization of Fshr-ZsGreen with CD133 staining in the cells of bone marrow and on the bone surface. Empty white arrows indicate bone marrow cells positive for both Fshr-ZsGreen and CD133 staining, and empty white arrowheads indicate Fshr-ZsGreen-positive cells on the trabecular bone surface with positive staining for CD133. d. Positive staining for CD133 on Fshr-ZsGreen-positive chondrocytes on the surface of articular cartilage. e. CD133 staining in the weak Fshr-ZsGreen-positive fibroblast-like cells in the periosteum. **D.** Reduced Fshr expression by Fshr cKO in osteocytes. Immunofluorescence staining with Fshr antibody was performed in decalcified frozen sections of femurs from the control and inducible osteocytes Fshr cKO mice (DMP1-CreERT^2+^:Fshr^fl/fl^ treated tamoxifen and DMP1-CreERT^2-^:Fshr^fl/fl^ as the control treated with corn oil). White empty arrows-osteocytes in the control; Green empty arrows-osteocytes with reduced Fshr expression in the cKO. Dotted lines indicate the thickness of the cortex. Magnifications: 400X. Scale bars: 50μm.

In contrast, we found that Fshr was brightly expressed in osteoblasts and osteoclasts in the metaphyseal trabeculae, which were recognized based on nuclear DAPI staining- osteoblasts were stained with a single DAPI nucleus, while osteoclasts contained more than two DAPI-stained nuclei (Fig. 3A-c and d). Similarly, we further observed that these cells on the surfaces of trabeculae in bone marrow and cortical bone also clearly expressed Fshr-ZsGreen, as shown in Fig. 3A-e and f (trabeculae) and-g and h (cortex). Importantly, we noted Fshr-ZsGreen expression in osteocytes (indicated by empty white arrowheads, Fig. 3A-d, g and h),the most abundant cell in the skeleton. These were embedded in trabecular and cortical bone, as well as in the periosteum (P) (Fig. 3A-h).

To confirm the identification of osteoblasts/osteocytes and osteoclasts that expressed Fshr, we performed immunofluorescence (IF) staining using an antibody against osteocalcin, a marker of osteoblasts ^31^. As shown in Fig. 1B, we observed colocalization of Fshr expression with osteocalcin staining in chondrocytes in the transforming zone, as indicated by white arrows (Fig. 3B-a), in cuboid and nucleated osteoblasts and bone lining cells on the trabecular and endosteal surfaces (Fig. 3B-b and c), cortical bone, and osteocytes within the mineralized matrix (Fig. 3B-b, c and e).

To identify osteoclasts, we performed fluorescence immunostaining with an antibody against Trap, an osteoclast marker ^32^. Osteoclasts were recognized by positive staining for TRAP with more than two DAPI-stained nuclei. Multinucleated Fshr-ZsGreen-positive cells were stained positively for Trap located on the surface of the resorptive bays or areas adjacent to the trabecular bone (Fig. 3B-e and f).

To examine whether skeletal stem/progenitor cells express Fshr, we performed IF staining for stem markers with antibodies against CD34 or CD133 ^33–37^. Using these well- known stem markers, we identified Fshr-ZsGreen-positive cells as stem/progenitor cells located in the bone marrow, growth plate/articular cartilage, and periosteum, as shown in Figure 2C-a, b, c, e and d, respectively. These cells also featured an increased nuclear- cytoplasmic ratio, except for these cells on the trabecular surface and in the periosteum.

We also examined Fshr expression in osteocytes from DMP1-CreERT^2+^:Fshr^fl/fl^ mice (osteocyte-specific Fshr cKO) where the control mice (DMP1-CreERT^2-^:Fshr^fl/fl^) as a positive control. We noticed a significant drop of Fshr expression in osteocytes in Fshr cKO mice when tamoxifen was intraperitoneally administrated to the mice (Fig. 3D). We also found an increased thickness of cortex triggered by Fshr cKO (Fig. 3D-the left panels, the thickness indicated by dotted lines). The finding further demonstrated Fshr present in osteocytes, the largest cell population in bones, and the specificity of the Fshr antibody.

### 3) Adipose tissues

Because our previous works and others have provided evidence on the role of Fsh in adipose tissues, we then examined Fshr-ZsGreen expression in adipose tissues. As expected, we found Fshr-ZsGreen expression in adipocytes of the frozen sectioned inguinal WAT, as shown in the left panel of Figure 4A at a low magnification (40X), which was further confirmed under a higher magnification (400X) in the three representative areas, demonstrating that Fshr- ZsGreen was expressed in the cellular membranes of individual adipocytes (Fig. 4A-a, b and c).

**Figure 4.**
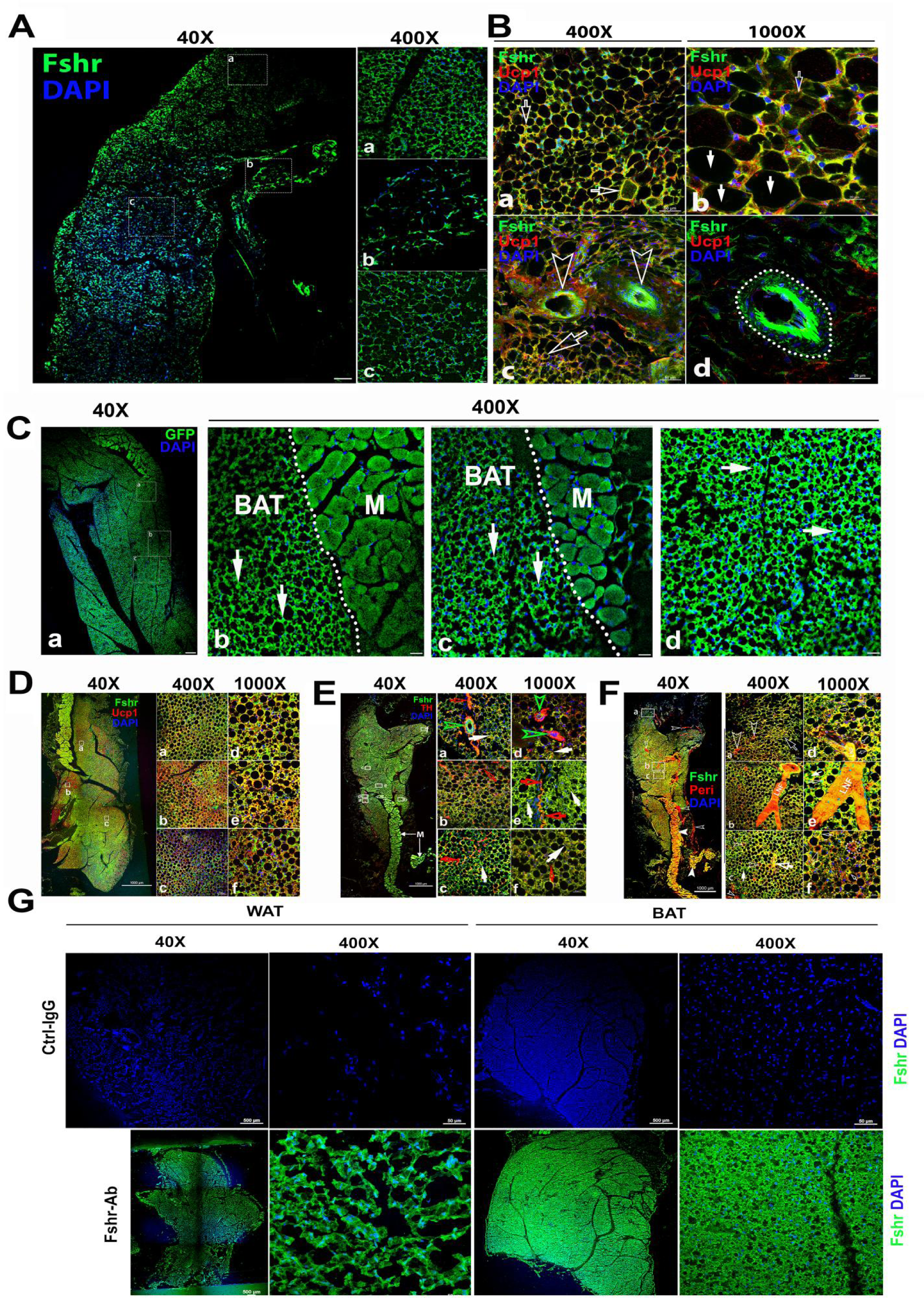
Fshr-ZsGreen expression in adipose tissues. **A.** Confocal imaging of Fshr-ZsGreen expression in frozen sections of inguinal WAT. The whole image of a piece of the inguinal WAT was imaged under confocal microscopy with a low magnification (40X), and three representative areas were also presented with a higher magnification (400X) showing low and high cellular densities (low-b and high-a and c). These images demonstrate Fshr-ZsGreen expression in the cellular membranes of adipocytes. **B.** IF staining of Fshr-ZsGreen-positive adipocytes with an antibody against mouse Ucp1. To identify the beige cells in WAT, we performed IF staining with an antibody against mouse Ucp1. Positive staining for Ucp1 was colocalized within Fshr-ZsGreen- positive cells in the areas with a higher cell density (a and b) imaged at low magnification, as shown in the left panels, while higher magnification images are shown in the right panels (a and d). In addition, strong Fshr-ZsGreen expression was observed in the arterioles of WAT (c and d). Arrows: white empty arrows indicate beige cells; white arrows point to white adipocytes, and empty arrowheads indicate GFP-positive arterioles. Magnifications: 400X for a and c; 1000X for b and d. Scale bars: 50 μm for a and c and 20 μm for b and d. **C.** Fluorescence images of Fshr-ZsGreen expression in frozen sectioned BAT. A whole image of BAT was imaged under confocal fluorescence microscopy at low magnification (a, 40X). Three representative areas are also presented at a higher magnification (b, c and d, 400X), which clearly show the Fshr-ZsGreen expressed in the cells of BAT and skeletal muscles in the right parts of the images (b and c). Abbreviation: M-muscle. Arrows: white arrows indicate strong Fshr-ZsGreen expression in BAT cells. Magnifications: 40X for a and 40°C for b, c and d. Scale bars: 100 μm for a and 50 μm for b, c and d. **D.** Identification of brown adipocytes in BAT by IF staining. Brown fat cells were identified using an antibody against mouse Ucp1. Ucp1 staining is shown at lower magnification (40X) in the left panel covering a whole piece of BAT. Three representative areas are imaged at higher magnifications (a, b and c, 400X and 1000X), showing Fshr- ZsGreen colocalization with Ucp1-positive staining. Magnifications and scale bars are indicated in the figure. **E.** Examination of peripheral neural fibers within BAT. To determine whether Fshr- ZsGreen is expressed in the peripheral nerves in BAT, we performed IF staining with an antibody against tyrosine hydroxylase (TH), a marker for peripheral sympathetic neurons. A whole piece of BAT was imaged at a lower magnification (40X) after being stained for TH, as shown in the left panel. Representative areas are presented in the right panels with higher magnifications (400X -a, b and c and 1000X-d, e and f, respectively). Arrows: red outlined arrowheads indicate Fshr-ZsGreen- and TH-positive large peripheral nerves; green outlined arrowheads point to Th-stained nerve fibrils accompanying Fshr-ZsGreen- positive nerve fibrils around a Fshr-ZsGreen-positive arteriole; red arrows indicate Fshr- ZsGreen- and TH-positive nerve fibrils; and white arrows indicate brown adipocytes with both Fshr-ZsGreen and TH expression. Magnifications: 40X for the whole image of BAT in the left panel; 400X for a, b and c; and 1000X for a, e, and f. Scale bars: indicated in the images. **F.** Further confirmation of Fshr-ZsGreen expression in the peripheral nerves. To confirm Fshr-ZsGreen expression in peripheral neurons, we also employed another antibody against Peripherin (Peri), a 57-kD type III intermediate filament that is a specific marker for peripheral neurons, to further identify peripheral neurons in BAT. The left panel shows an entire image of the BAT stained for Peri at a lower magnification (40X). The three representatives are shown in the right panel: a. the first area located in the edge of the BAT with more white adipocytes; b. the second area with more brown adipocytes and a large peripheral nerve; c. the last area enriched with brown adipocytes. Their corresponding higher magnifications (1000X) are shown in a, e and f, respectively. Abbrev.: LNF-large never fibril. Arrows: Empty white arrows-small peripheral fibrils; white arrow-Fshr-ZsGreen-positive fibrils and empty white arrows-both Fshr-ZsGreen- and Peri-positive brown adipocytes. Magnifications: 40X for the t panel; 400X for a, b, and c and 1000X for d, e and f. Scale bars: 1000 μm for the whole images in the left panel; 50 μm for a, b and c, and 20 μm for d, e and f. **G.** Detection of Fshr expression in adipose tissues of B6 mice. Immunofluorescence staining for Fshr expression was carried out in frozen sections of white adipose tissue (the left panels) and brown fat (the right panels) from B6 mice at age of 3 months. Magnifications: 40X and 400X. Scale bars: 500 μm and 50 μm.

To confirm adipocyte identification, we performed IF with an antibody against mouse Ucp1 ^12^ to recognize adipocytes, showing that the majority of Fshr-ZsGreen-positive cells were stained positively for Ucp1, and found two types of Fshr^+^ adipocyte populations: one with colocalization of the two markers only in the membrane (indicated by empty white arrowheads) and another with two markers in both the membrane and cytoplasm (indicated by white arrows) (Fig. 4B-a and b). In addition, we observed that Fshr-ZsGreen was expressed in arterioles in adipose tissue, denoted by white empty arrowheads in Figure 4B-c and a dotted circle in Figure 4B-d.

We further examined Fshr-ZsGreen expression in BAT. Fshr-ZsGreen was observed across the whole section of examined BAT at a lower magnification (Fig. 3A-a). At a higher magnification (400X), three representative areas are presented, as shown in Figure 3A-b, c and d, in which Fshr-ZsGreen was expressed not only in the cellular membranes but also in areas of cytoplasm close to the membranes, as indicated by white arrows (Fig. 4C-b, c, and d).

Furthermore, we also performed IF with three antibodies against Ucp1, Th and Peri to identify brown cells and peripheral fibers in the Fshr-positive section. We found that several areas in the section were strongly stained for Ucp1 at low magnification (40X), as shown in Figure 4D-a. Under higher magnifications of 400X and 1000X, we colocalized Fshr-ZsGreen with Ucp1 in the three representative areas, in which some locations had higher Fshr-ZsGreen expression and others had higher Ucp1 expression (Fig. 4D-a, b and d).

To examine whether peripheral neural fibers express Fshr, we used antibodies against Th and Peri that can recognize peripheral neural fibers. We found that neural fibers stained positively and surrounded Fshr-ZsGreen-positive arterioles in BAT (Fig. 4E-a and b). In addition, Fshr-ZsGreen was expressed in the nodes of Ranvier of TH-stained small neural fibers (indicated by empty red arrows, Fig. 4E-e). We further confirmed Fshr- ZsGreen expression in peripheral neural fibers by IF with an antibody against peripherin (Peri), which is a type III intermediate filament protein found predominantly in peripheral nerves, specifically in sensory and autonomic neurons (Fig. 4F). We noted colocalization of Fshr-ZsGreen and Peri staining in large neural fibers (Fig. 4F-b, d and e). Interestingly, we observed that both markers for peripheral neural fibers were also expressed in the cytoplasm of BAT cells, in which Fshr-ZsGreen was strongly expressed (Fig. 4E-b, c and f; Fig. 4Fe, c and f).

To further ensure Fshr expression in adipose tissues, we also performed immunofluorescence staining with Fshr antibody in white and BAT adipose sections of B6 mice. The results showed the similar expression pattern as seen in Fshr-ZsGreen mice (Fig. 4G).

Taken together, the above-described findings on Fshr-ZsGreen expression in reproductive, skeletal, and adipose tissues convincingly demonstrate that Fshr-ZsGreen is a reliable readout of Fshr expression. Furthermore, we identified Fshr expression in Leydig cells and follicles at different developmental stages, cells of osteoblast lineage and peripheral nerve fibers. With confidence that Fshr-ZsGreen is a reliable readout, we used this powerful tool to further examine Fshr expression in other tissues and organs.

### 4) Heart and aorta

To examine Fshr expression in the cardiovascular system, we used the heart and aorta as key organs/tissues to detect Fshr-ZsGreen expression. As expected, we observed strong Fshr-ZsGreen expression in the myocardium (Fig. 5A-a) and large muscular arteries (a representative is shown in Fig. 5A-b). Then, we further confirmed Fshr-ZsGreen expression by IFs with two antibodies against α-SMA and EMCN ^21^ that recognize alpha smooth muscle actin of smooth muscle and endomucin of endothelial cells at higher magnifications. With IF staining for α-SMA, we imaged several areas of both heart and blood vessels, as shown in the left image of the upper panel (40X). In the heart, we observed that Fshr-ZsGreen was highly expressed in cardiomyocytes in longitudinal and transverse orientations of the myocardium, which were positively stained for α-SMA (Fig. 6B-a, b and c). At a magnification of 1000X, it was also expressed in the endothelial layer of arterioles between muscle fibers (Fig. 5B-i).

**Figure 5.**
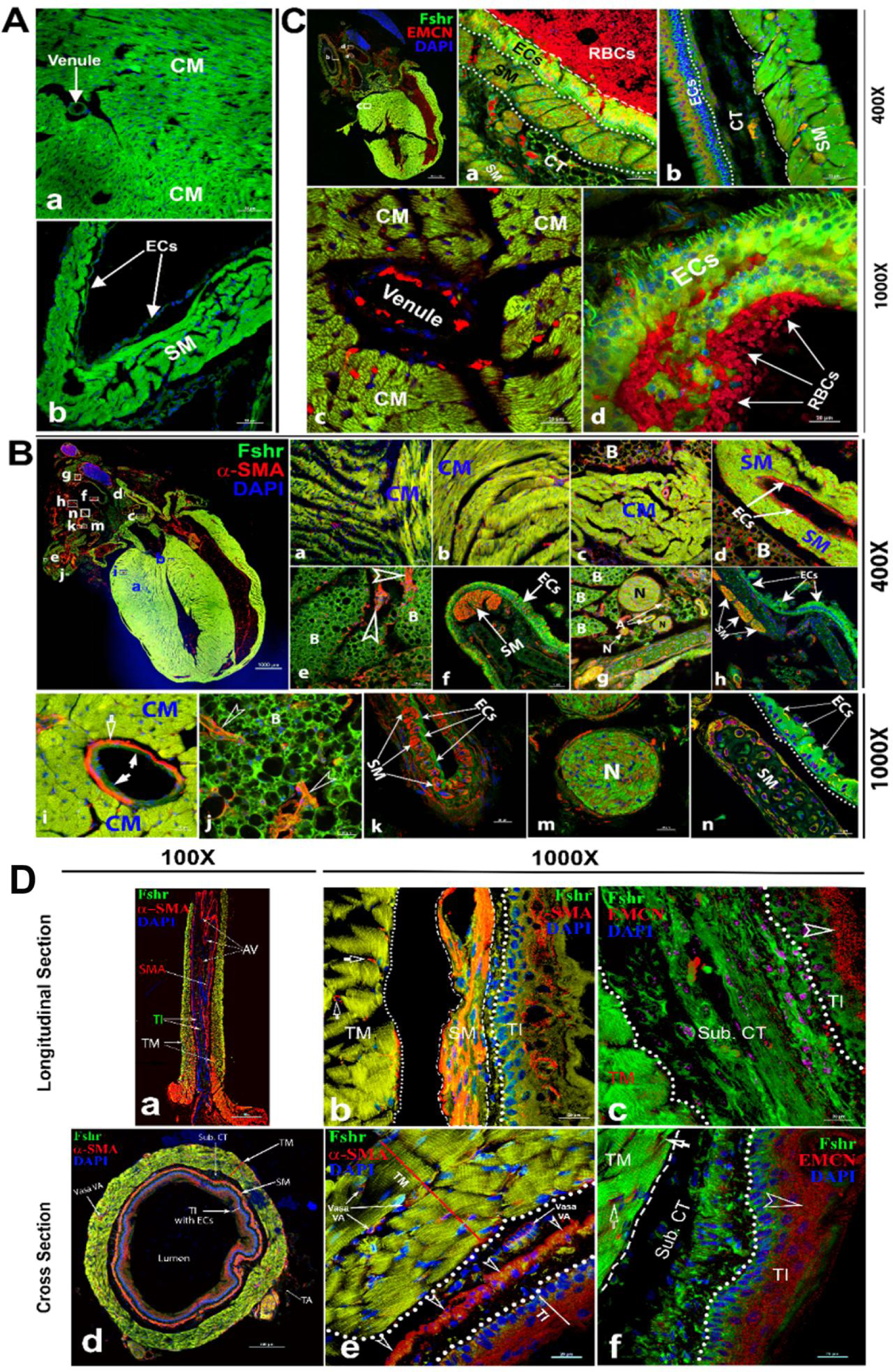
Imaging of Fshr-ZsGreen expression in the heart and aorta. Fshr-ZsGreen expression was imaged in frozen sections of the heart (A): cardiomyocytes (a) and smooth muscles (b). Then, IF staining with antibodies against α- SMA (B) and EMCN (C) was carried out to identify cardiomyocytes and smooth muscles. The whole image of the heart with Fshr-ZsGreen expression and staining for α-SMA is shown in the left upper panel of B. Its representative areas are presented at two magnifications of 400X (B-a to h) and 1000X (B-i to n), respectively, in the following: (1) 400X magnification: a-cross-oriented cardiomyocytes; b-longitudinally oriented cardiomyocytes; c-cross-oriented smooth muscle of the ascending aorta with brown adipose tissue; d-smooth muscle of the left pulmonary artery with brown adipose tissue; e-brown adipose tissue attached to a large artery; e-layer of endothall cells of the superior vena cava (SVC); g. connective tissues between arteries containing brown adipose, different sized nerves and arteries; h. another part of the SVC with layers of smooth muscles and connective tissue. (2) 1000X magnification: i, cardiomyocytes with an arteriole; j. brown adipose tissue attached to the circulation system above the heart; k. the layers of endothelial cells and smooth muscles of a bronchial artery; m. transverse section of nerve fibers; n. the layers of ECs and SM of the SVC. In addition, imaging of the IF staining for EMCN is shown in C. The whole image of Fshr-ZsGreen and EMCN staining in the heart at a magnification of 40X in the left upper panel, from which a representative of pulmonary artery at a magnification of 400 is shown in a at a magnification of 400X (a); a representative of aorta is presented in b. Representative images at higher magnification of 1000X: c-cardiomyocytes in transverse orientation with a venule and d-a layer of endothelial cells of pulmonary artery. Abbrev.: CM- cardiomyocytes; SM-smooth muscle; ECs-endothelial cells; N-never; B-brown adipose. Magnifications and scale bars are indicated as in the figure. Two orientations of the ascending aorta were examined for Fshr-ZsGreen after staining for α-SMA or EMCN: the longitudinal section (D-a to c) and the cross section (D-d to f)). D-a shows the entire image of the ascending aorta stained for α-SMA at a magnification of 100X, while representative images at a higher magnification of 1000X are presented in D-b for α-SMA staining and D-c for EMCN staining. The entire image of the transversely sectioned aorta (D-d) and representative images at a higher magnification of 1000X for α-SMA (D-e) and EMCN (D-f). Abbrev.: TI- tunica intima, TM- tunica media, AV- aortic valve, SM- smooth muscles, Sub. CT-subendothelial connective tissue. Arrows: empty white arrows indicate arterioles positively stained for α-SMA in b; empty arrowheads point to endothelial cells positively stained for either α-SMA or EMCN. Scale bars: 100 μm for a and d and 20 μm for b, c, e, and f.

**Figure 6.**
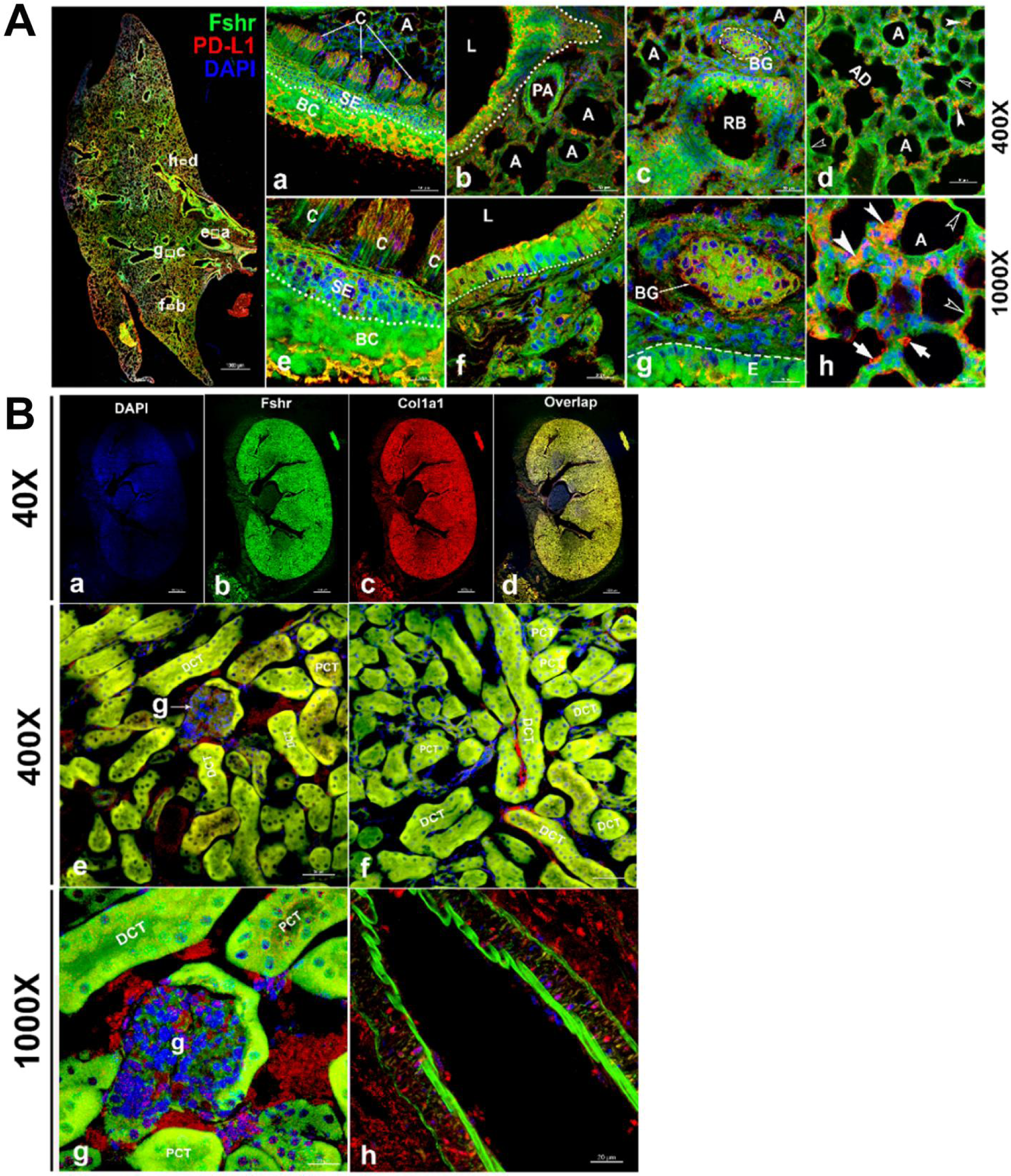
Fshr-ZsGreen expression in the lung and kidney. Detection of Fshr-ZsGreen expression and IF staining with PD-L1 was performed in lung sections at different magnifications. The whole image of the frozen sectioned lung is shown in the left panel at 40X magnification (A-left panel). Representative areas are shown in the right panels at magnifications of 400X (Figure 6A-a to d) and 1000X (A-e to h). They are the ciliated columnar cells of primary bronchi (a and c), the bronchioles with alveoli (b and f), respiratory bronchiole with bronchial gland (c and g) and alveoli (d and h). Arrows: empty white arrowheads-type I pneumocytes, white arrowheads-type II pneumocytes and white arrows-macrophages. Abbrev.: C-ciliated epithelium; PA- pulmonary arteriole; L-lumen; RB-respiratory bronchiole; BG-bronchial gland; A-alveoli; AD-alveolar duct. Magnifications: 40X for the left mage of the whole section, 400X for a to d, and 1000X for e to h. Scale bars: 1000 μm for the left panel, 50 μm for a to d and 20 μm for e to h. Fshr-ZsGreen expression and staining with Col1a1 were examined in the sectioned kidney under confocal fluorescence microscopy at three magnifications (B). The images in the top panel are images of the whole section at 40X magnification (a-d). The images in the middle panel show the colocalization of Fshr expression with positive staining for Col1a1 in the glomerulus and renal tubes (proximal and distal convoluted tubes) at 400X magnification (e and f), while the bottom panel shows images of the glomerulus (g) and arteriole (h) at 1000X magnification. Magnification: 1000X. Scale bars: 100 μm for a-d, 50 μm for e and f and 20 μm for g and h. Abbrev. A-arteriole; PCT- proximal convoluted tubule; DCT-distal convoluted tubule; g-glomerulus.

In addition to the cardiomyocytes, we also observed Fshr-ZsGreen in α-SMA-stained smooth muscles and endothelial cells of large blood vessels above the heart (Fig. 7B-d). Interestingly, we found Fshr-ZsGreen in adipose tissue around the blood vessels, and the adipocytes morphologically appeared to be brown adipose cells, as the majority of these brown-like cells were full of ZsGreen-positive cytoplasm, instead of single large fat droplets with Fshr-ZsGreen expression in cellular membranes (Fig. 5B-c, d, e and j). The adipose tissue was stained positively for α-SMA, suggesting that the ZsGreen^+^ structures costained for α-SMA are blood vessels within the beige tissues, which are indicated by white empty arrowheads (Fig. 5B-e and j).

**Figure 7.**
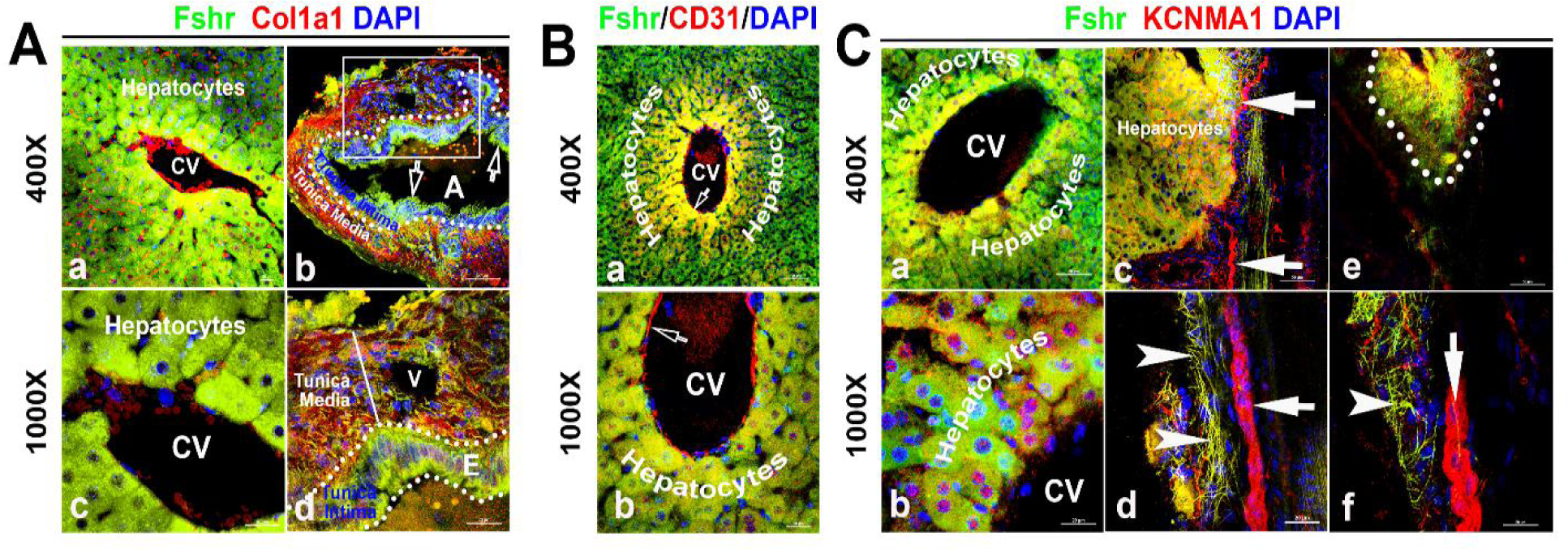
Identification of Fshr-ZsGreen expression in the liver. Frozen sectioned liver tissue was stained for Col1a1 (A), CD31 (B) or KCNMA1 (C), followed by florescent imaging for Fshr-ZsGreen and each of these stained molecules simultaneously. In the section stained for Col1a1 (A), two representative areas are shown at magnifications of 400X and 1000X as indicated: (1) hepatic cells with a central vein (CV) (a and c) and (2) hepatic artery (b and d). In the section stained for CD31 (B), a representative area of hepatic cells with a CV is shown at lower (a) and higher (b) magnifications. In the section stained for KCNMA1, three representative areas are presented at two magnifications as follows: (1) hepatic cells with a CV (a and b); (2) hepatic cells with peripheral nerve fibers (c and d); and (3) small nerve fibers located around a vein (e and f).

Furthermore, we observed bright Fshr-ZsGreen expression with slightly weak staining for α-SMA in the layer of endothelial cells of the large vein in Figure 5B-f and h. Under the endothelial cells, a cluster of smooth muscles showed colocalization of Fshr-ZsGreen with positive staining for α-SMA (Fig. 5B-f and h). In contrast to the vein, we observed a thin layer of endothelial cells (tunica intima) and stronger Fshr-ZsGreen expression with positive staining for α-SMA in the large artery (tunica media); Fshr-ZsGreen was also present in the cells in the tunica adventitia (Fig. 5B-k). In frozen sections of hearts, we observed strong Fshr-ZsGreen expression in neural fiber clusters that were in adipose tissue, as shown in Figure 5B-g and m.

Using an antibody against EMCN, a marker for endothelial cells, we further confirmed Fshr-ZsGreen expression in the layer of endothelium (tunica intima), which showed visible positive staining for EMCN when imaged at 400X and 1000X (Fig. 5C-a, b and d). In addition, Fshr-ZsGreen^+^ cardiomyocytes and smooth muscles under the endothelial layer were positive for EMCN staining (Fig. 5C-a, b and c).

In addition to large blood vessels on the heart, we also took a close look at the ascending aorta. We obtained two types of sections with longitudinal and transverse orientations for IFs with the two antibodies as above. In both sections, we found that Fshr-ZsGreen^+^ smooth muscle fibers were strongly stained for α-SMA in the first layer of SM close to the endothelium, where the second layer of smooth muscle was relatively weak for staining with α-SMA (Figs. 5D-a and 6B-e). Fshr-ZsGreen was also present at the endothelium, which was costained positively for α-SMA (Fig. 6A-a and 6B-e). Using an anti-EMCN antibody, we noticed that positive staining was in the upper part of Fshr- ZsGreen^+^ endothelial cells facing the lumen of the examined blood vessels (Fig. 5A-c and 5B-f). In addition to Fshr-ZsGreen expression in the endothelium of the tunica intima, it was also observed in the areas of subendothelial connective tissue and tunica media (smooth muscle), but these areas were not stained for EMCN (Fig. 5A-c and 5B-f).

### 5) Lung and kidney

To identify whether Fshr is expressed in epithelial cells, we first detected Fshr- ZsGreen in the lung. Unexpectedly, we observed high Fshr-ZsGreen expression in the lung (Fig. 6A). Fshr-ZsGreen was brightly expressed in the columnar epithelium of the respiratory conducting zone/tract, including the trachea, bronchus, bronchi and bronchiole, at low magnification (Fig. 6A-left panel). At higher magnifications, it was clearly shown that Fshr-ZsGreen was expressed not only in the columnar epithelium but also in the bronchial gland and alveoli (Fig. 6A-a, b, d, e, f and g). In the alveoli, Fshr-ZsGreen was observed in the respiratory portion of both type I and II cells, as indicated by empty arrowheads and white arrows, respectively (Fig. 6A-d and h). We confirmed the identification of respiratory cells by IF with an anti-PD-L1 antibody, which showed the colocalization of Fshr-GFP with PD-L1 ^38^ staining (Fig. 6A).

Then, we aimed to examine Fshr expression in epithelial cells of the kidney. In the frozen section of the kidney, it was astonishing to observe high expression of Fshr- ZsGreen in the proximal and distal convoluted tubules (Fig. 6B-e and f), whereas relatively weak expression was observed in the glomerular capillaries at different magnifications (Fig. 6B-e and f at 400X and g at 1000X). Again, we observed Fshr-ZsGreen expression in the endothelial layer of the arteriole in the kidney tissue (Fig. 6B-h). We also observed colocalization of Fshr-ZsGreen with positive staining for Col1a1 ^34^ in the kidney (Fig. 6B).

### 6). Other key tissues and organs (liver, pancreas, thyroid, skin and skeletal muscle, spleen, bone marrow, and brain)

With the power of Fshr-ZsGreen, we characterized Fshr-ZsGreen expression in the liver. We observed Fshr-ZsGreen expression in the hepatocytes and arteries inside the hepatocytes, which were positively stained for Col1a1 or CD31 ^39,40^, respectively (Fig. 7-A and B). Although it is weakly expressed inside large nerve fibers, Fshr-ZsGreen is strongly expressed in small neural fibers and shows a costaining pattern with KCBMA1 ^41^, a marker used for the detection of peripheral nerve fibers (Fig. 7-C, indicated by white arrowhead, whereas large nerve fibers are indicated by white arrows).

We then examined Fshr-ZsGreen expression in the pancreas and found that it was expressed not only in acinar cells but also in islets of Langerhans at low and high magnifications (Fig. 8-A-a and b). We then confirmed the identification of α and β-cells by IFs with antibodies against NG3, insulin or glucagon. The imaging results clearly demonstrated that Fshr-ZsGreen was expressed in both α and β-cells as well as acinar cells at 400X and 1000X magnifications, respectively (Fig. 8-A, -B and -C).

**Figure 8.**
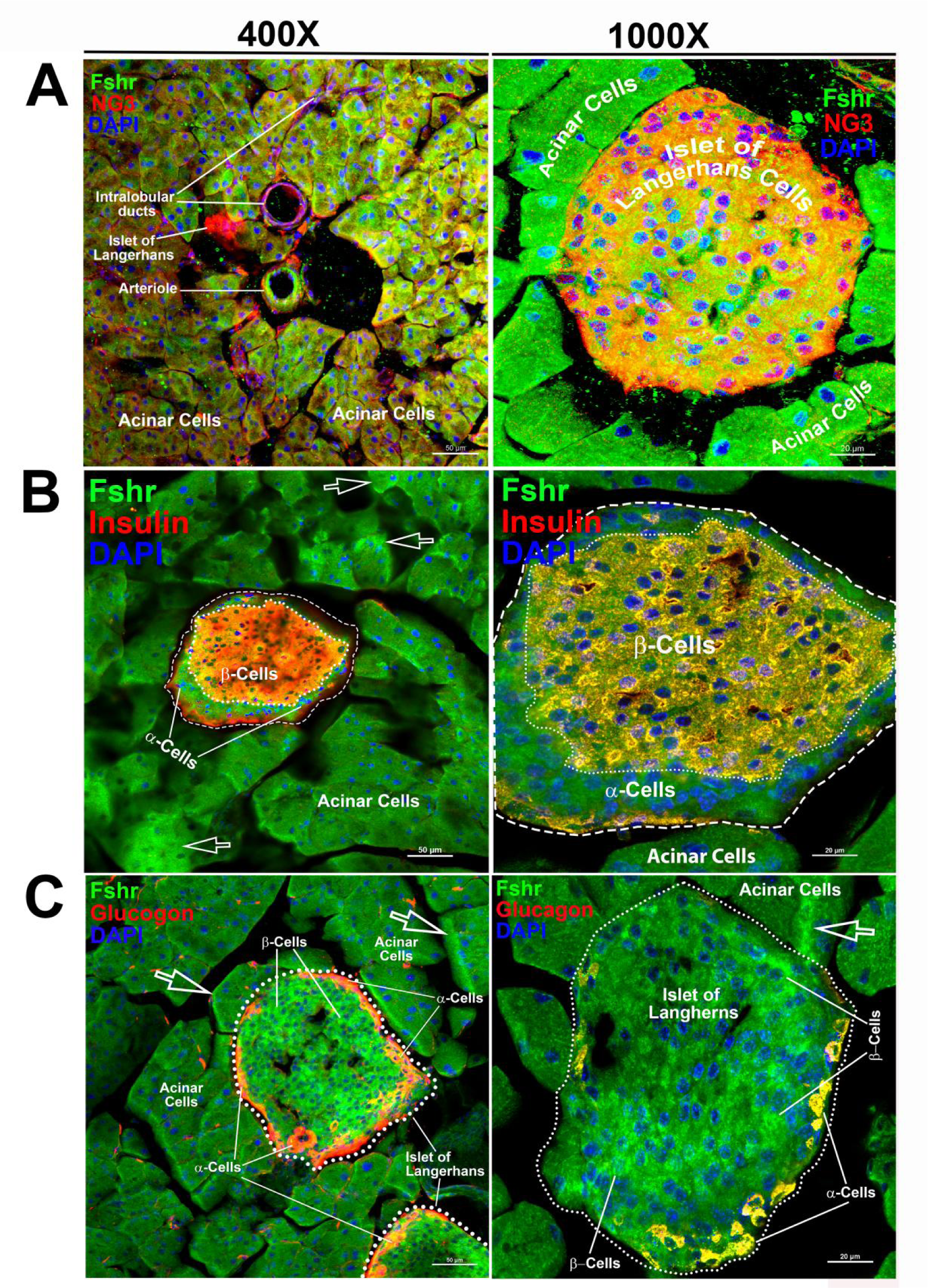
Visualization of Fshr-ZsGreen expression in the pancreas. Visualized Fshr-ZsGreen expression was obtained in frozen sections of the pancreas under fluorescence microscopy after immunostaining with antibodies against NG3, insulin or glucagon at two magnifications (400X and 1000X). Images of Fshr- ZsGreen expression with NG3, insulin and glucagon staining are shown in A, B and C, respectively.

Regarding Fshr-ZsGreen expression in the thyroid, we found Fshr-ZsGreen in both follicular cells and parafollicular cells (C-cells) at low and high magnifications (400X and 1000X) (Fig. 9-A-a, b, c, d and e). We further used an anti-Tsh receptor antibody to confirm Fshr-ZsGreen-positive cells, which showed that follicular cells were stained positively for Tshr in the nuclei, as indicated by dotted circles and white arrowheads, and C-cells were indicated by white arrows (Fig. 9-b to e).

**Figure 9.**
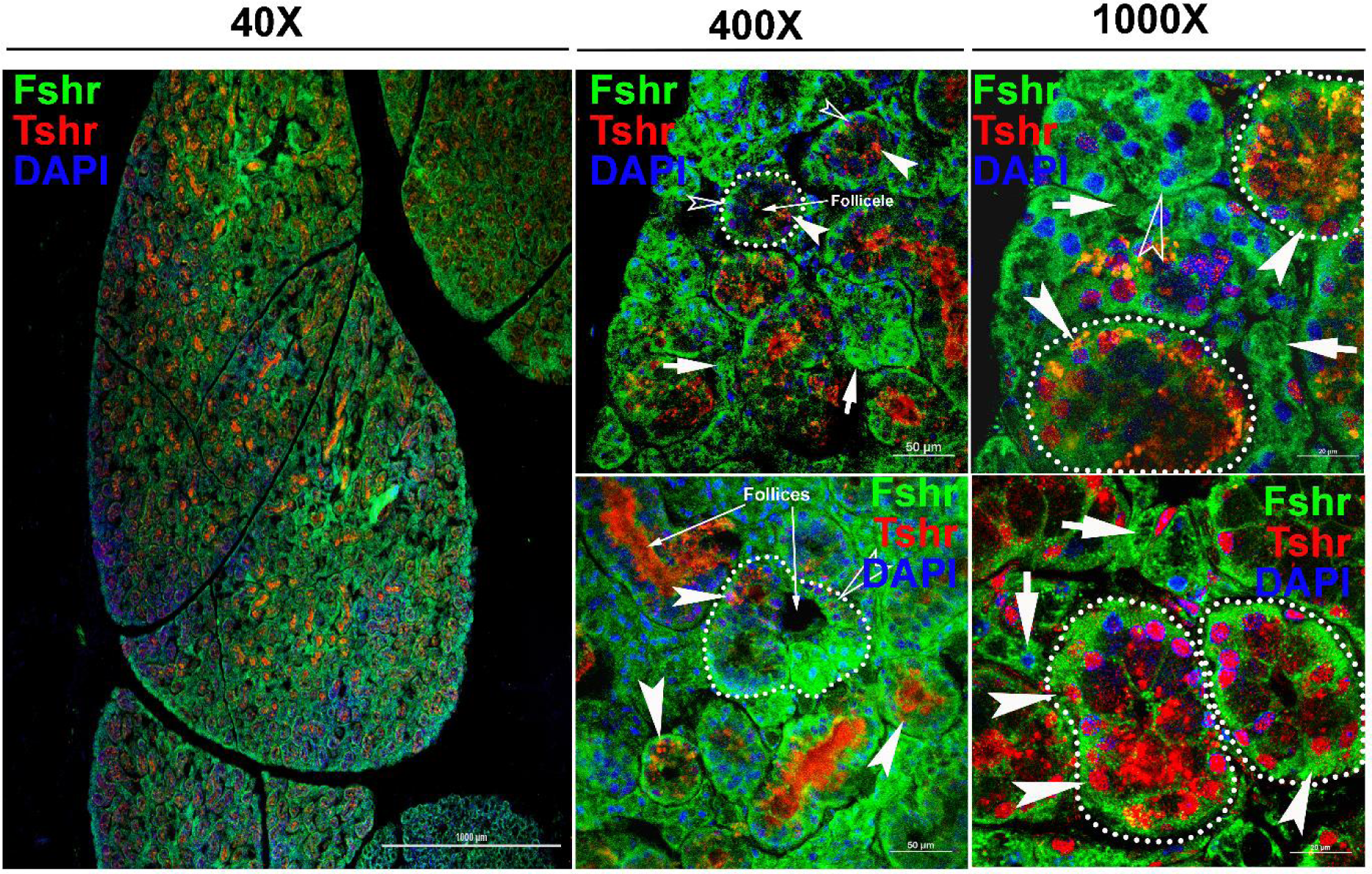
Fshr-ZsGreen expression in the thyroid. Fshr-ZsGreen expression was detected by its colocalization with immunostaining of TSH in frozen thyroid sections. The left panel is an entire image of the section, and two representative areas are in the right panels at higher magnifications, demonstrating Fshr-ZsGreen expression in follicular cells and parafollicular cells (C cells) at the edge (a and b) and the center of the thyroid (c and d). Arrows: white arrowheads indicate follicular cells, and white arrows indicate parafollicular cells. Magnification: 40X for the whole image.

Next, to examine Fshr-ZsGreen expression in the skin, we used two types of skin sections: thick skin from the tail and thin skin from the abdomen (Fig. 10-A-a and b). Although a dermis layer was not included in the image taken for a thick sample, we observed that Fshr-ZsGreen was expressed in hair follicle (HF) and sweat gland cells and fibroblasts in the dermis and hypodermis (Fig. 10-A-a). Similarly, Fshr-ZsGreen was present in HFs and fibroblasts in the dermis and keratinocytes in the epidermis of the thin skin (Fig. 10-A-b). As we detected Fshr-ZsGreen expression in hair follicles, we wondered whether stem cells in HFs express Fshr-ZsGreen. To address this question, we carried out IF with an antibody against CD34, a stem cell marker. Not surprisingly, we found that

**Figure 10.**
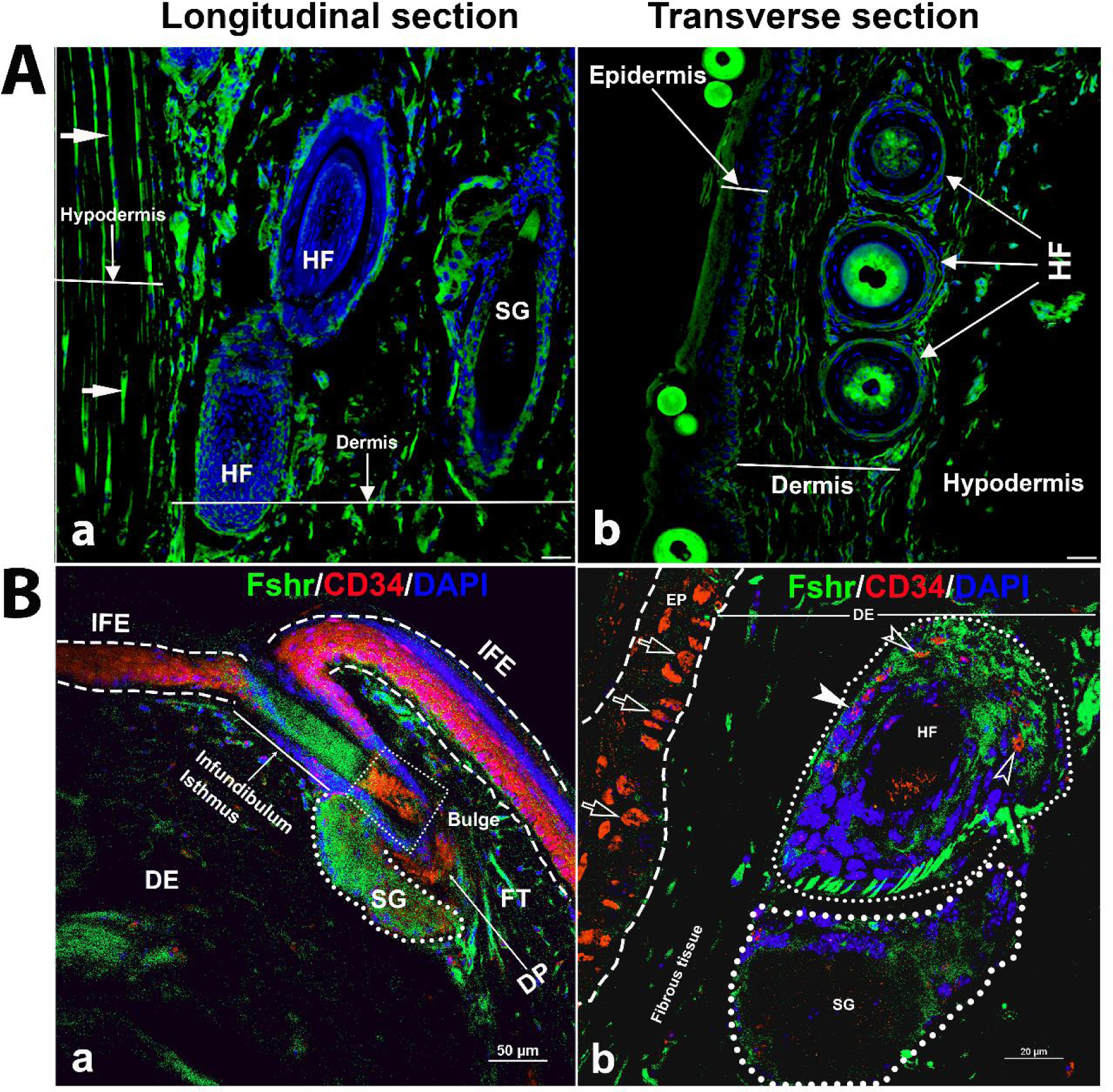
Detection of Fshr-ZsGreen expression in the skin. Fluorescence images of Fshr-ZsGreen were taken in two types of skin (A): thick skin (A-a) and thin skin (A-b). Then, IF staining was performed using an antibody against CD34 to identify stem cells with Fshr-ZsGreen expression in the hair follicles (B) at lower (B-a) and higher (B-b) magnifications (400X and 1000X, respectively). Abbrev.: HF-hair follicle; SG-sweat gland; FT-fat tissue; DP-dermal papillae. Magnifications: 400X for A and B-a, 1000X for B-b. Scale bars: 50 μm for A and B-a and 20 μm for B-b.

CD34 staining was colocalized with cells with Fshr-ZsGreen in Bulge as quiescent stem cells and in the dermal papilla (DP) and epidermis as active stem/progenitor cells, as shown in Figure S4-B-a and b. Therefore, these results indicate Fshr-ZsGreen expression in stem/progenitor cells in the skin (Fig. 10B-a and d).

We then detected Fshr expression in skeletal muscle (gastrocnemius), followed by IF staining using three antibodies against αSMA, PAX7 and TH for the identification of satellite cells and peripheral nerve fibrils. We observed Fshr-ZsGreen across the muscle sections at a lower magnification of 40X (Fig. 11a). At higher magnifications, we found that Fshr-ZsGreen was present in the muscle fibers, in which one type had higher Fshr- ZsGreen expression (indicated by white arrowheads) and another type had lower Fshr- ZsGreen expression (Fig. 11b to h). Fshr-ZsGreen was also highly expressed in satellite cells stained positively for either αSMA (Fig. 11-b to e) or PAX7 (Fig. 11-f and g) in both longitudinal and transverse sections, as indicated by white arrows. In addition, we detected Fshr-ZsGreen expression in the peripheral neural fibers identified by positive staining for TH, which were around vascular structures in the muscle tissue, as indicated by red arrowheads (Fig. 11-h and i).

**Figure 11.**
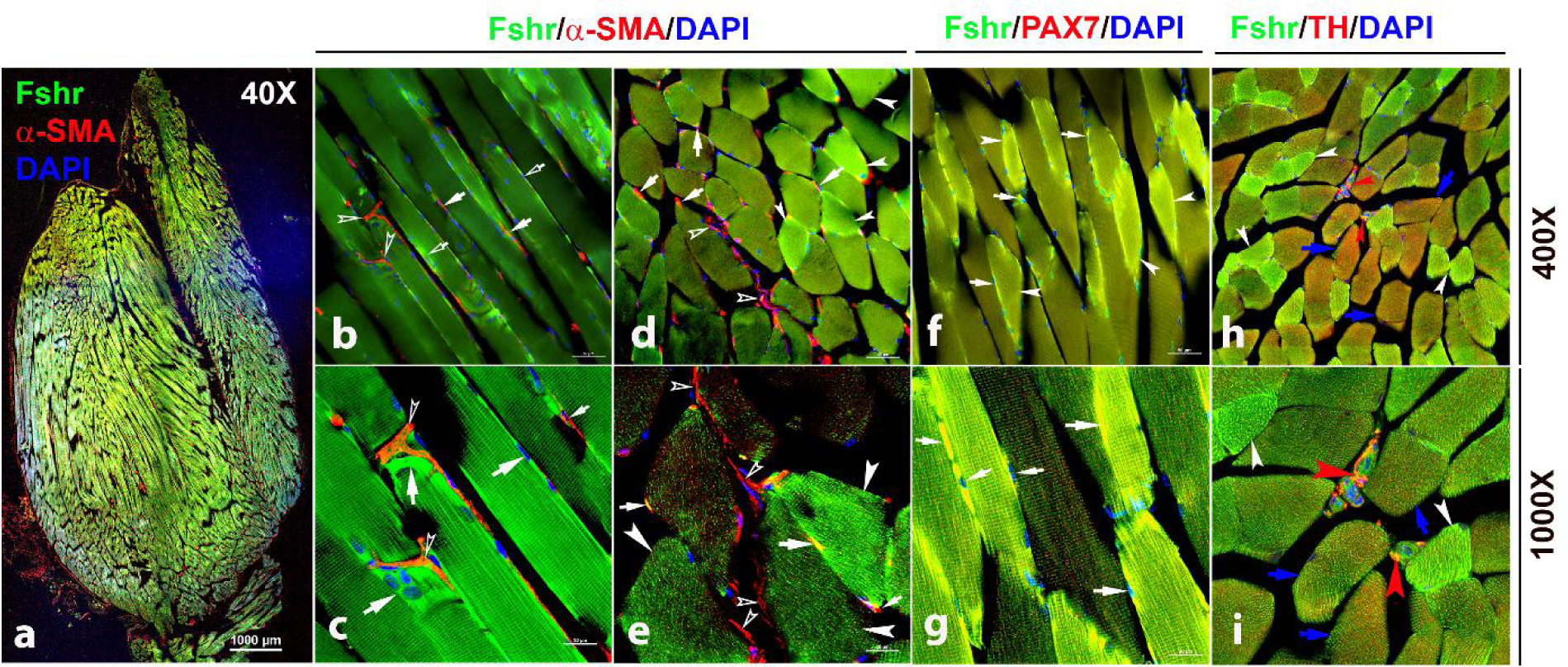
Fshr-ZsGreen expression in skeletal muscle. The whole image of frozen sectioned skeletal muscle (gastrocnemius) was taken at a lower magnification (40X) after the section was stained with anti-α-SMA antibody, as shown in the left panel (a). Then, longitudinal sections (b and c) and cross-sections (d and e) were imaged at higher magnifications (400X and 1000X), respectively. In addition, frozen sections stained with antibodies against PAX7 (f and g) and TH (h and i) were imaged at two magnifications (400X and 1000X) as indicated. Arrows: (1) In b-e, empty white arrowheads indicate both Fshr-ZsGreen- and α-SMA-positive satellite cells; empty white arrows point to only Fshr-ZsGreen-positive satellite cells without α-SMA staining; white arrowheads indicate muscle fibrils with strong Fshr-ZsGreen expression but no α- SMA staining. (2) In f and g, white arrowheads indicate muscle fibrils with strong Fshr- ZsGreen expression, blue arrows point to muscle fibrils with both Fshr-ZsGreen and TH expression, and red arrowheads indicate peripheral nerves with both Fshr-ZsGreen and TH expression. Magnifications: 40X for a, 400X for b, d, f and h, and 1000X for c, e, g and i. Scale bars: 1000 μm for a, 50 μm for b, d, f and h, and 20 μm for c, e, g and i.

To detect Fshr expression in immune cells, we examined Fshr-ZsGreen expression in the spleen and bone marrow using antibodies against CD11b and CD3, as integrin αM (CD11b) is expressed in myeloid-lineage cells such as monocytes/macrophages, neutrophils, eosinophils, and basophils and in lymphoid cells such as NK cells and B-1 cells ^42^, while CD3 marks T cells ^43^. We imaged frozen sections of the spleen under magnifications of 40X, 400X and 1000X and observed that Fshr-ZsGreen was highly expressed in trabeculae and cells in both red and white pulps (RP and WP). The Fshr- ZsGreen-positive cells were further identified by IFs with antibodies against CD11B or CD3, indicating that Fshr is expressed in immune cells, such as monocytes/macrophages, neutrophils, eosinophils, basophils, NK cells, B cells and T cells (Fig. 12-A and B). In addition, we further confirmed Fshr-ZsGreen expression in macrophages of bone marrow, which were identified with anti-CD4/80 antibody, as shown in two representative images of BM-one from an area close to cortical bone (Fig. 12-C-a) and another located in the center of BM (Fig. 12C-b) at 1000X magnification (empty white arrows indicate the colocalization of Fshr-ZsGreen and CD11B, CD3 or CD4/80 in the RPs, while white arrows point to their colocalization in the WPs).

Finally, we examined Fshr-ZsGreen expression and its colocalization with markers for either astrocytes, microglia, or neurons in the brain. As expected, we observed Fshr expression across the brain sections and the three representatives from the olfactory bulbs, pallidum, and hippocampus (CA3) are shown in Figure 12, demonstrating that Fshr is indeed expressed in astrocytes (Fig. 13-a to c), microglia (Fig. 12-d to f) and neurons (Fig. 13-g to j), as well as neuronal fibers (synapses or projections). Other cell types are needed to be further defined.

**Figure 13.**
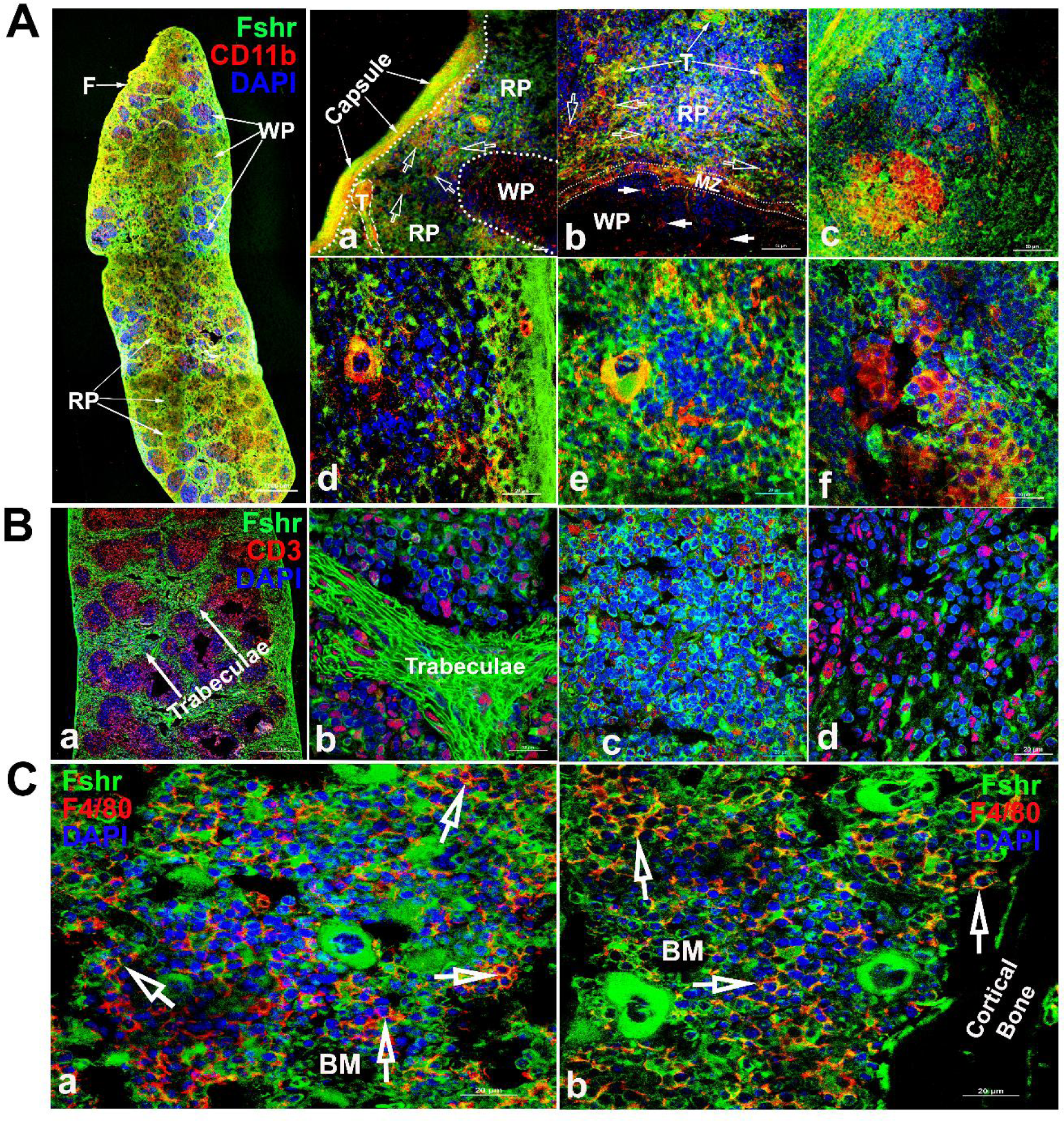
Examination of Fshr-ZsGreen expression in the spleen and bone marrow. To identify Fshr-ZsGreen expression in immune cells of the spleen, IF staining was performed in frozen sections of the spleen. Two antibodies against either CD11B or CD3 were used to identify myeloid-lineage cells or T cells. Fshr-ZsGreen expression in myeloid-lineage cells is shown in A, in which the left panel is the whole image of the spleen at a low magnification and its three representative areas are presented at higher magnifications (400X and 1000X): (1) an area located at the edge showing strong Fshr- ZsGreen and CD11B expression (a and b); (2) an area of red pulp c and d); (3) an area of white pulp (e and f). Abbrev.: RP-red pulp, WP-white pulp, BM-bone marrow, F-fibroelastic capsule and MZ- marginal zone. Scale bars are indicated in each image.

### 3. Confirmation of Fshr-ZsGreen expression with a Fshr-specific antibody and ddRT- PCR

Finally, to confirm the accuracy of the above results obtained from Fshr-ZsGreen mice, we performed fluorescence immunostaining with a specific antibody against mouse Fshr and accurate droplet digital RT‒PCR (ddRT-PCR) with mouse Fshr-specific primers to confirm the above data. An isotype-matched rabbit IgG was used as a negative control for IFs with anti-Fshr antibody using sections from Fshr-ZsGreen mice. Imaging was performed under the same conditions to record each corresponding tissue/organ stained with anti-Fshr antibody. The images of negative controls are shown in Supplementary Data 2, showing specific binding of the secondary antibody to anti-Fshr antibody without any nonspecific binding of the secondary antibody to the examined sections.

As shown in Figure 14A, in all the examined tissues, including bone, BAT, thyroid, cardiac muscle, kidney, liver, lung, aorta, ovary and testis, we observed colocalization of Fshr-ZsGreen with positive staining for Fshr, further confirming the specific expression of Fshr-ZsGreen in the examined tissues/organs.

**Figure 14.**
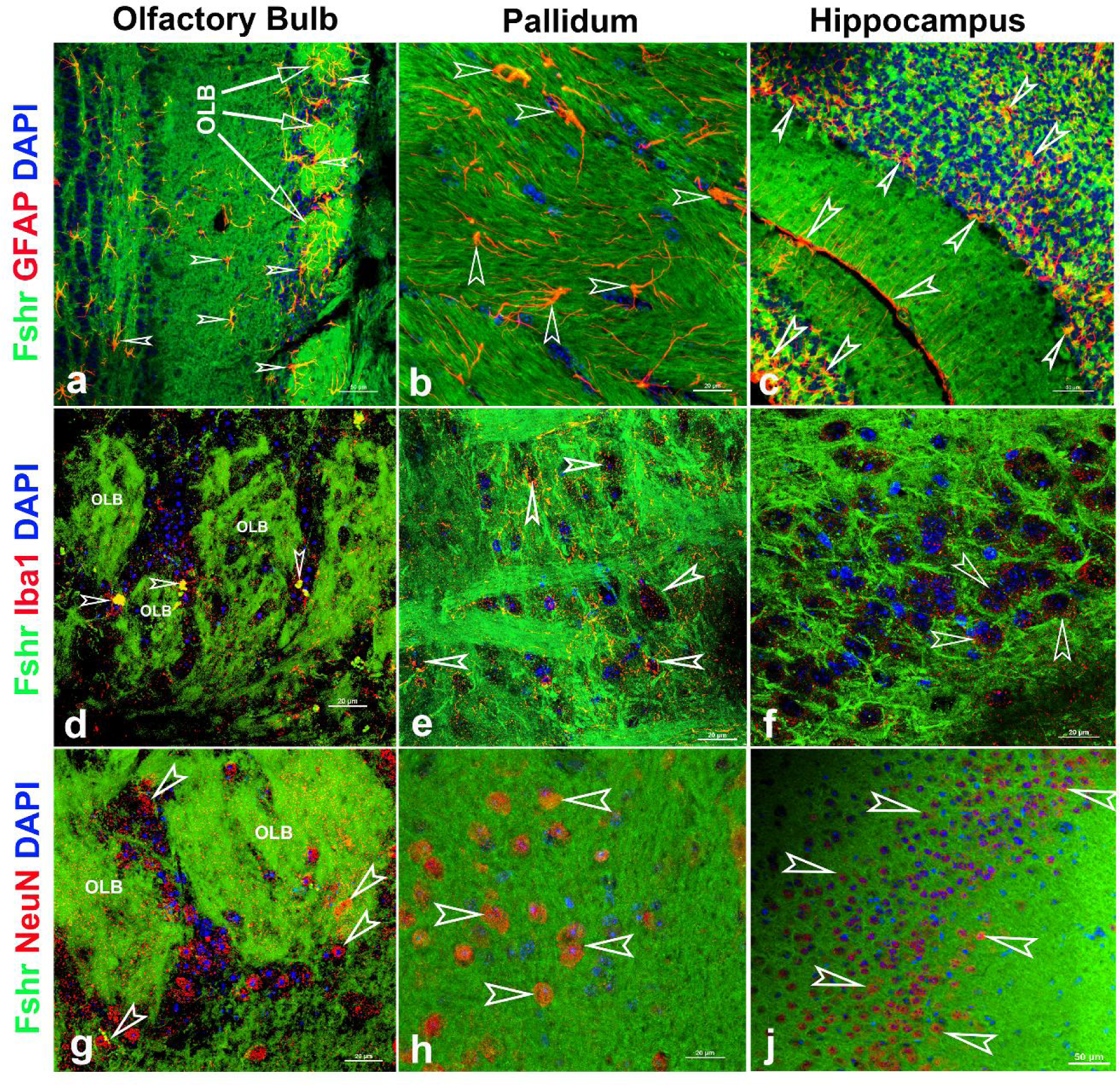
Fshr-ZsGreen expression in the representative areas of the brain. Fshr expression in three representative areas of the brain-the olfactory bulbs (a, d, and g), pallidum (b, e and h) and hippocampus (c, f and j). Each section was immunofluorescently stained with antibodies against GFAP, Iba1 or NeuN that recognized markers for astrocytes (a-c), microglia (d-f) or neuron (g-j), respectively. Their colocalizations are indicated by white arrowheads. Abbrev.: OLB-olfactory bulb. Magnifications: 400X for a, c and f, and 1000X for b, d, e, f g and h. Scale bars are indicated in each image.

**Figure 15.**
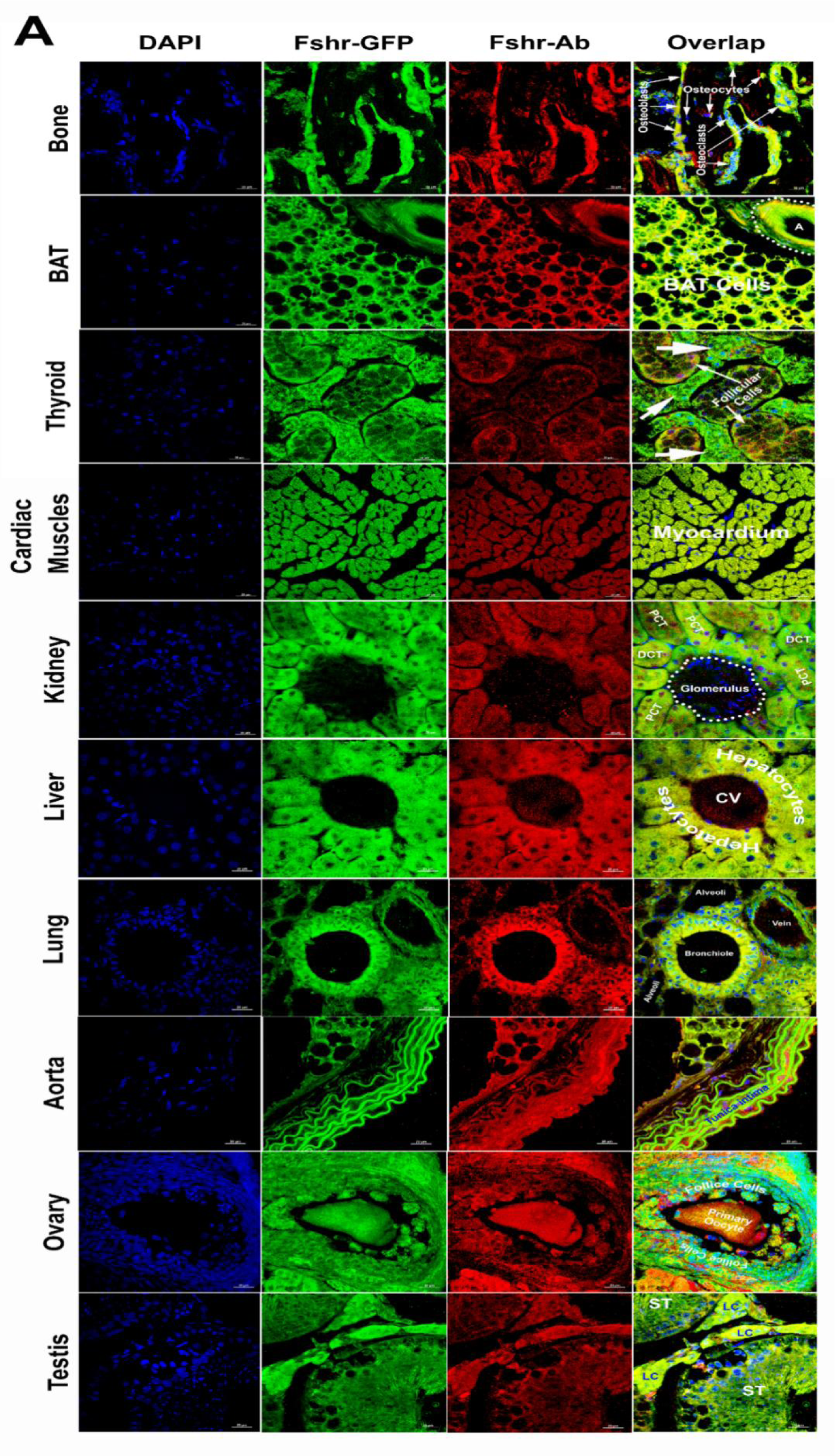
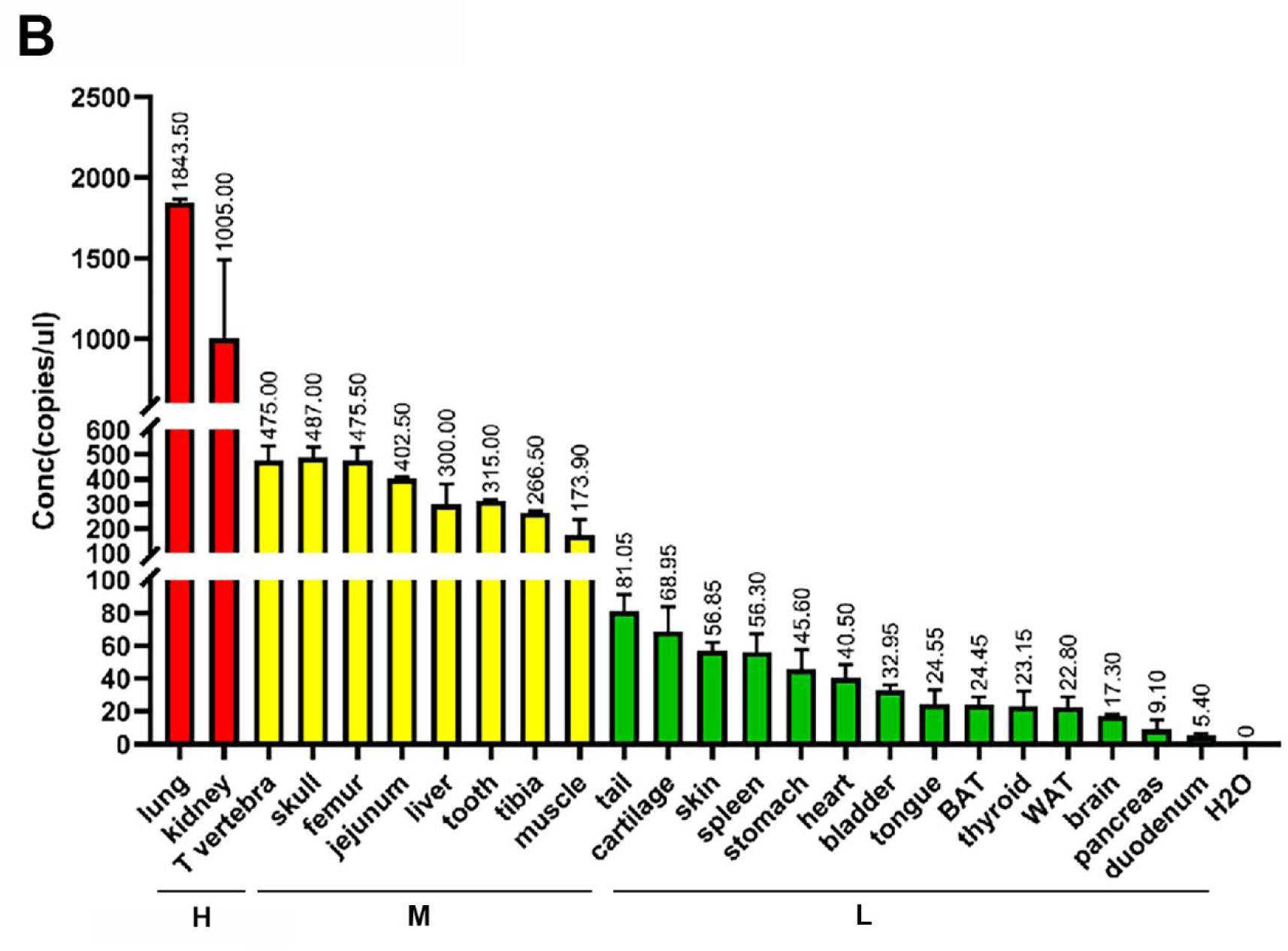
Confirmation of Fshr-ZsGreen expression by IF staining with a specific antibody against mouse Fshr and ddRT-PCR. The Fshr-ZsGreen expression described above was further confirmed by IF staining using an antibody against mouse Fshr and ddRT‒PCR. The frozen tissue sections used for this confirmation include bone, BAT, thyroid, cardiac muscles, kidney, liver, lung, aorta, ovary and testis (A), demonstrating Fshr-ZsGreen colocalization with Fshr-positive staining in these tissues/organs. Fshr expression at the mRNA level in different tissues/organs was examined by ddRT‒PCR (B) (the representative of two experiments). The results indicate that Fshr is expressed in all examined tissues/organs. Based on their expression levels, they can be categorized into three groups: (1) high (H), including lung and kidney, ranging from 1005 to 1843 copies/μL; (2) middle (M), including thoracic (T) vertebra, skull, femur, jejunum, liver, tooth, tibia and muscle, ranging from 173 to 475 copies/μL; and (3) low (L), including duodenum, pancreas, brain, WAT, thyroid, BAT, tongue, bladder, heart, stomach, spleen, skin, cartilage and tail, ranging from 5.4 to 81 copies/μL.

We also obtained total RNA from the following tissues: lung, kidney, thoracic vertebra, calvaria, femur, jejunum, liver, teeth, tibia, skeletal muscle, tails, cartilage, skin, spleen, stomach, heart, bladder, tongue, BAT, WAT, thyroid, brain, pancreas and duodenum. After reversing mRNA from total RNA, we performed ddRT-PCR with mouse-specific primers to check Fshr expression at the transcriptional level as described in the Methods section. The results demonstrated Fshr expression in these tissues, and the expression profile was categorized into three groups: 1) high expression in the lung and kidney tissues; 2) moderate expression in the thoracic (T) vertebrae, calvaria, femur, jejunum, liver, teeth, tibia, and muscle; and 3) low expression in the tail, cartilage, skin, spleen, stomach, heart, bladder, tongue, BAT, WAT, brain, pancreas, and duodenum (Fig. 14B).

Finally, we also searched Fshr expression at single cell level in 5 single cell databases including DISCO (a database of Deeply Integrated Single-Cell Omics data, (https://www.immunesinglecell.org/), BioGPS (A free extensible and customizable gene annotation portal, a complete resource for gene expression and protein function, http://biogps.org/exrna/#goto=welcome), Single Cell Portal (SCP, an interactive home for single-cell genomics data, https://singlecell.broadinstitute.org/), Genotype-Tissue Expression (GTEx portal, https://gtexportal.org), CZ CELLxGENE Discover (https://cellxgene.cziscience.com/).

The selected results from these databases are presented in Supplementary Data 3, which further support our findings of the widerspread express pattern of Fshr described as above. In particular, Fshr expression was detected in Leydig cells by scRNA-seq as shown in DISCO (immunesinglecell.org/genepage/FSHR), BioGPS (http://biogps.org/#goto=genereport&id=2492) and CZ CELLxGENE Discover (human- https://cellxgene.cziscience.com/e/535e9336-2d8d-43c3-944d-bcbebe20df8a.cxg/ and mouse- https://cellxgene.cziscience.com/e/a13bda79-9134-46c9-9ed1-a2858be9aafe.cxg/).

Taken together, these data further confirmed Fshr-ZsGreen expression patterns from the Fshr-ZsGreen reporter line, convincingly demonstrating that Fshr is not limited to previously reported cells, tissues, or organs, such as the reproductive system, osteoclasts, adipose, endothelium in tumors, and neurons in the brain, but rather has a wide expression in the cells, tissues, and organs in the body, particularly in the lung, kidney, and heart, as well as Leydig cells in the testis, which has not previously been recognized.

## Discussion

Although numerous efforts have been made to characterize Fshr expression in tissues/cells, defining the locations of Fshr expression remains an imperative challenge in Fsh-Fshr biology, primarily because of concerns about the specificity of available antibodies against Fshr ^2,15,16^. In this case, we developed CRISPR/Cas9-mediated Fshr- ZsGreen knockin reporter mice to address this issue.

To maintain the integrity of the splicing acceptor, donor, and gene promoter regions, we designed and inserted the ZsGreen (ZsG) reporter into the the C-terminus of Fshr by sequence-specific gRNA guided CRISPR/Cas9-mediated precise genome modification with a long ssDNA template to create the Fshr-P2A-ZsG reporter mice ^44^. Because of its high turnover of Fshr mRNA and a difficulty in detection of Fshr expression by rt-PCR and Northern blotting, in this reporter line, we utilized ZsG as a GFP reporter, as ZsGreen, also called ZsGreen1, is an exceptionally bright green fluorescent protein derived from a Zoanthus sp. reef coral ^45^ that has been modified for high solubility, bright emission, and rapid chromophore maturation. ZsGreen is the brightest commercially available green fluorescent protein—up to 4X brighter than EGFP with the half life of 26 hours and is ideally suited for whole-cell labelling and promoter-reporter studies to indicate the promoter activity of a gene of interest, which has been used in numerous GFP reporter mice ^46–55^. In addition, we also employed a short and conserved picornavirus-derived ‘self-cleaving’ 2A peptide to allow bicistronic expression of Fshr and ZsG, as the 19-amino acid P2A has the highest cleavage efficiency ^56,57^. The cleavage is triggered by ribosomal skipping of the peptide bond between the proline (P) and glycine (G) in C-terminal of P2A peptide, resulting in the peptide located upstream of the 2A peptide to have extra amino acids on its C-terminal end, while the peptide located downstream the 2A peptide will have an extra proline on its N-terminal end. Therefore, P2A can nearly equalize the expression of genes upstream and downstream. As a result, Fshr is normally expressed without interruptions and ZsG expression indicates Fshr promoter activity in the defined cells, demonstrated by the comparison of Fshr expression in the testes and ovaries between Fshr-ZsGreen and B6 mice (Fig. 2G).

The P2A-ZsGreen construct was precisely inserted into the site between the last exon (exon 10) and the stop codon of the Fshr locus, which was confirmed by integration detection PCR, sequencing of PCR fragments and Southern blotting with Fshr locus- specific enzyme restriction digestions. Successful insertion allows the endogenous promoter of Fshr to drive ZsGreen reporter expression. Because of the site-specific insertion of the P2A ZsG vector to produce Fshr-ZsG reporter, we only used one founder line for characterizing Fshr-ZsG expression, rather than multiple founders when random insertions of a transgene were used previously for generating transgenic mice.

This approach greatly enhanced our understanding of this critical cellular pathway in several ways. Firstly, employing the native regulatory element responsible for governing the expression of the target gene ensures that the expression pattern of the GFP reporter closely mirrors that of the gene of interest, Fshr, within its natural context. Consequently, this approach provides a more faithful representation of gene expression. Secondly, endogenous promoters often exhibit specific spatial and temporal expression patterns, driving gene expression in specific cell types or developmental stages. This capability facilitates the capture of the dynamic nature of gene regulation and enables the study of gene expression under diverse physiological conditions. Thirdly, utilizing endogenous promoters minimizes potential perturbations to the native gene regulation machinery. It avoids the need to introduce exogenous elements or artificial constructs, reducing the likelihood of altering the gene’s expression behavior or interfering with its regulatory interactions. Unlike biochemical assays or immunostaining, using a tagged protein under endogenous regulation avoids fixation artifacts and allows detection of the target’s activity in live cells. Therefore, these distinct advantages of enhanced physiological relevance, precise spatiotemporal control, preservation of regulatory elements, and minimal perturbation provide us with a powerful tool to understand Fshr expression in more accurate and context-specific ways compared to other methods, such as using antibodies, Northern blotting, RT‒PCR and in situ hybridization.

In this study, we systemically investigated Fshr expression at the single-cell level in this reporter line with confocal fluorescence microscopy and further confirmed the location of Fshr-ZsGreen expression by IF staining, in situ hybridization and ddRT‒PCR. The results from this work demonstrate that as a receptor for Fsh, Fshr is widely expressed in virtually every cell at variable levels in the examined tissues/organs of mice. As expected in the examined testis, we noticed Fshr expression in Sertoli cells in the testis and granular cells in the ovary. Surprisingly, we observed that Fshr was also more strongly expressed in Leydig cells than in Sertoli cells. This expression was also detected in spermatocytes, spermatids, spermatozoa and spermatogonia. Although this finding is different from present thoughts that Fshr is only present in Sertoli cells but not in other cell types of the testis, our finding is in line with previous works in fishes (including African catfish, zebrafish, teleosts, and Japanese eels) ^58–63^, rats ^64^, dogs ^65^ and humans ^66^ performed by IHC and in situ hybridization. In humans, Fshr was also highly expressed in Leydig cells, although it was taken as non-specificity of the anti-Fshr antibodies. However, as shown in this study, this is not the case, because Fshr is widely expressed, as demonstrated in our work.

Furthermore, our findings are supported by a report of the failure of normal Leydig cell development resulting from the deficiency of Fshr but not Fsh-beta ^67^ and single cell RNA-seq studies in Hu sheep (originated from Mongolian sheep) ^68^ and in human as shown in Supplementary Data 3 [DISCO (immunesinglecell.org/genepage/FSHR), BioGPS (http://biogps.org/#goto=genereport&id=2492) and CZ CELLxGENE Discover (https://cellxgene.cziscience.com/e/535e9336-2d8d-43c3-944d-bcbebe20df8a.cxg/)].

Fshr expression in Leydig cells strongly indicates that Fsh-Fshr plays a role in the production of steroids in males. Unexpectedly, Leydig cell line TM3 expresses much lower Fshr, compared to the Leydig cells *in vivo* (Fig. 2B-G). It may explain that these cells do not respond well to FSH treatments ^30^ and indicates that this cell line may be not a typical Leydig cell population and new Leydig cell lines should be established in the future.

In the ovary, Fshr is highly expressed in follicles at different stages, from primordial cells, primary follicles, and secondary follicles to mature/Graafian follicles and the corpus luteum. In addition to granulosa cells, Fshr expression was observed in oocytes of follicles. Taken together, these data indicate that Fshr plays an intragonadal role in the ovary and testis beyond the granulosa and Sertoli cells.

In the skeletal system, we observed Fshr expression in vivo not only in osteoclasts, as reported previously ^11,69,70^, but also, interestingly, in osteoblast lineage cells, such as osteoblasts, bone lining cells, osteocytes and progenitor cells of the periosteum, as well as in chondrocytes. In previous reports, Fshr expression in osteoclasts was detected only by RT‒PCR, Western blot and immunostaining in cultures of primary murine precursors ^11,69,70^. Using the Fshr-ZsGreen reporter line, we visualized Fshr expression in multinucleated osteoclasts in frozen sections, clearly demonstrating Fshr expression in osteoclasts. Intriguingly, this powerful tool enabled us to examine Fshr expression in other cell types in bone. Surprisingly, we observed Fshr expression in cells of the osteoblast lineage, from osteoprogenitor cells and osteoblasts to osteocytes and bone lining cells. This finding indicates that Fsh may regulate not only osteoclast-mediated bone resorption but also osteoblasts for bone formation. To functionally prove the presence of Fshr in osteoblasts/osteocytes, we also deleted Fshr in osteocytes in an inducible model. The Fshr cKO induced in osteocytes significantly reduce Fshr expression and triggered an increase in the cortical thickness (Fig. 3D) and a much more profound increase in bone mass and a decrease in fat mass than blockade by Fsh antibodies (unpublished data), illuminating Fshr expression in the osteoblast lineage.

In addition to its expression in the reproductive system and skeletal system, we also strikingly identified other cell types that highly express Fshr: endothelial cells in blood vessels and epithelial cells in the lung and kidney. In every examined tissue/organ, we found that endothelial cells stained positively for CD34 lining on the arterioles had a higher expression of Fshr-ZsGreen than other cell types. This bright Fshr-ZsGreen is more obviously seen in large arteries, such as the ascending aorta and others located in the heart. Similarly, Fshr was detected in vessels in solid malignant tumors by immunohistochemistry or RT‒PCR ^71^. However, it was not seen in normal tissues or organs by these methods, possibly due to its rapid turnover, fast degradation, or selected antibodies. Strikingly, we detected the highest Fshr-ZsGreen expression in bronchial and bronchiole ciliated epithelial cells by both fluorescence microscopy and ddRT‒PCR. It was also more highly expressed in other cell types in the lung, such as type I pneumocytes (alveolar lining cells), type II pneumocytes (great alveolar or septal cells), and gland cells. However, the role of this unexpectedly high Fshr expression in the lung remains unknown. Similarly, we also found the second highest expression of Fshr- ZsGreen in renal epithelial cells in proximal and distal convoluted tubules but weak expression in renal corpuscles. In addition, our finding of Fshr expression in adipose tissues is consistent with the observations from previous works ^12,72^. Recently, Fshr expression was reported to be present in β-cells of the pancreas to regulate glucose- stimulated insulin secretion, further supporting our findings of the Fshr expression pattern ^73^.

In summary, we established and validated an Fshr-ZsGreen protein reporter in vivo that faithfully recapitulates endogenous Fshr expression at single-cell resolution. Our compelling findings reveal that in addition to gonadal tissues, Fshr is also highly expressed in extragonadal systems, such as the lung, kidney, heart, and pancreas. This will provide insight to better understand the biology of the Fsh-Fshr axis and its roles in the physiology and pathology of these tissues/organs. In addition to the above described, we detected Fshr expression in cells of the teeth and brain; those findings are not presented here because of space limitations and will be published elsewhere, except that three representative areas of the brain are shown in Figure 14.

Although the Fshr-ZsGreen reporter line is a powerful tool for detecting the location of Fshr expression, it is limited in the definition of individual isoforms of Fshr transcripts and the detection of their turnover, which should be addressed by specific antibodies and determined by the rates of transcription and RNA degradation, including Xrn1 for 5’-to-3’ degradation, exosomes for 3’-to-5’ degradation and nonsense-mediated decay ^74 53^.

## Acknowledgments

This study was supported by a fund from Shanxi Medical University to PL.

## Author contributions

Hong-Qian Chen, Conceptualization, Data curation, Investigation, Methodology, Formal analysis, Project administration; Hui-Qing Fang, Data curation, Investigation, Methodology, Formal analysis; Jin-Tao Liu, Data curation, Investigation, Methodology; Shi-Yu Chang, Data curation, Investigation, Methodology; Li-Ben Cheng, Data curation, Investigation, Methodology; Ming-Xin Sun, Data curation, Investigation, Methodology; Jian-Rui Feng, Data curation and Analysis; Ze-Min Liu, Resource and Analysis; Xiao-Li Li, Data curation and Analysis; Yong-Hong Zhang, Resource, Writing-Original draft; Clifford Rosen, Writing – original review and editing; Peng Liu, Conceptualization, Formal analysis, Funding acquisition, Investigation, Project administration, Resources, Supervision, Writing – original draft, Writing – review and editing.

## Supplementary Materials 1

1. The first set of Integration detection PCR primer design and sequences:

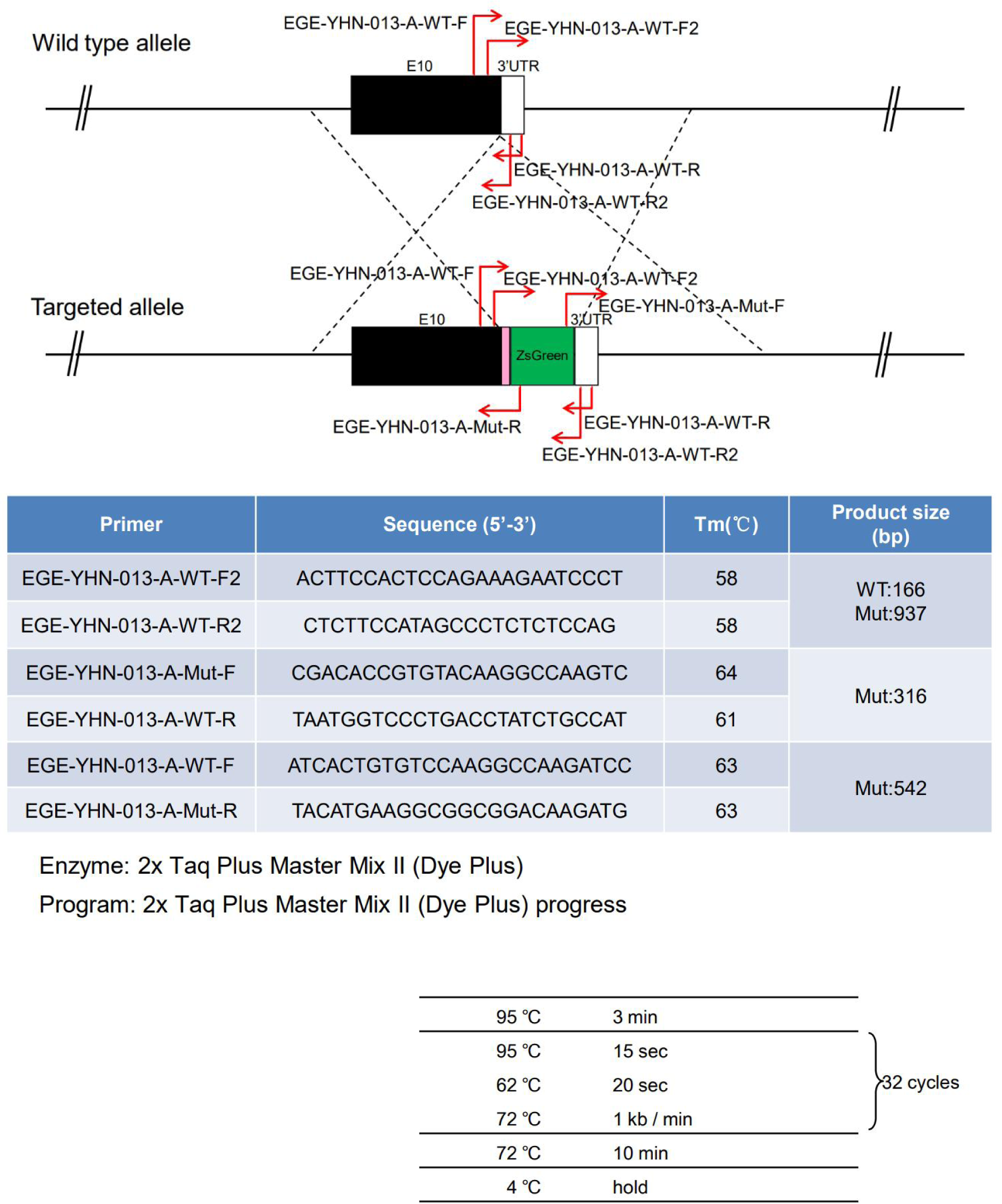

2. The 2^nd^ set of junction PCR primer design and sequences for genotyping F0 and F1:

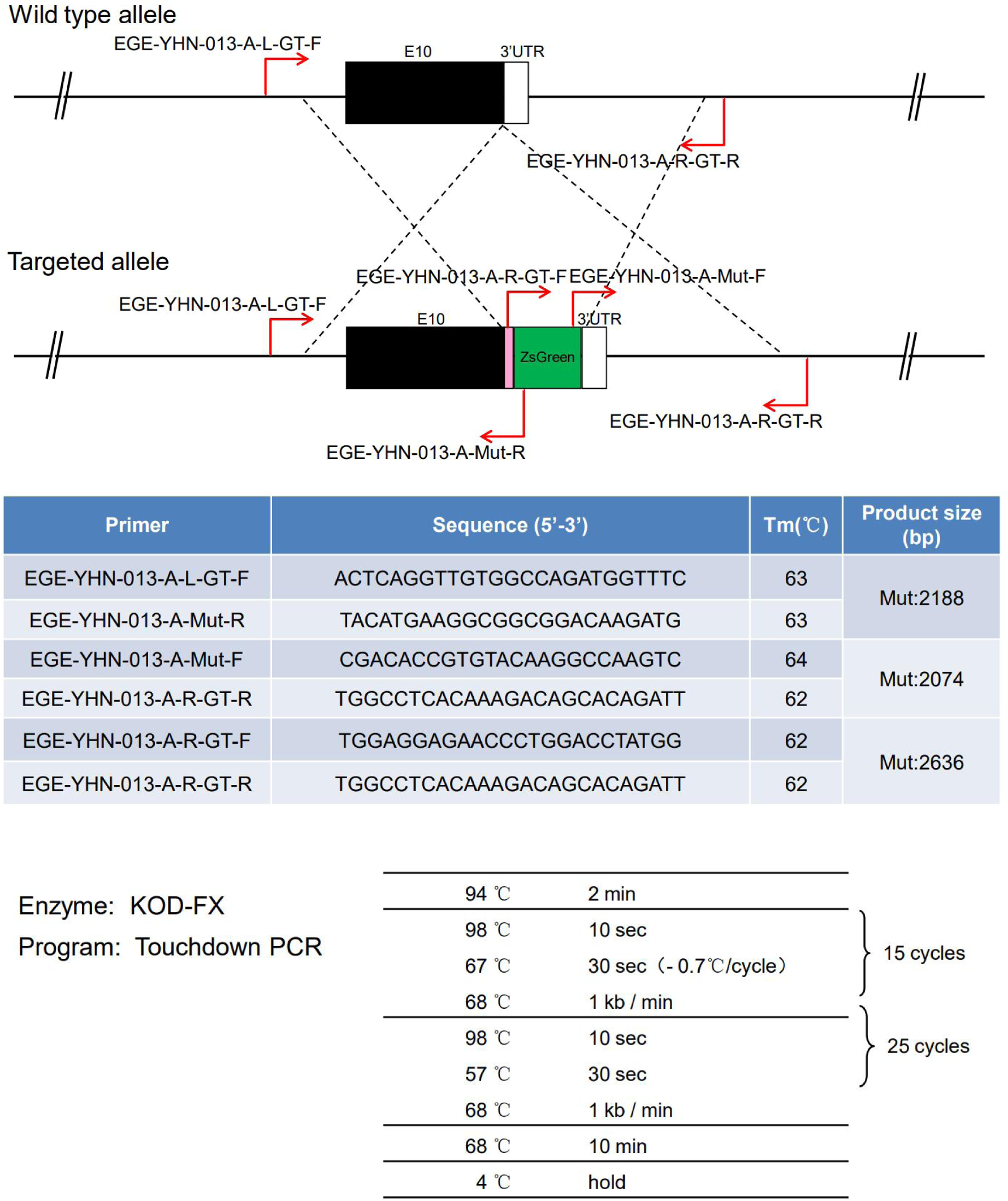

3. PCR primer design and sequences for genotyping heterozygotes and homozygotes:

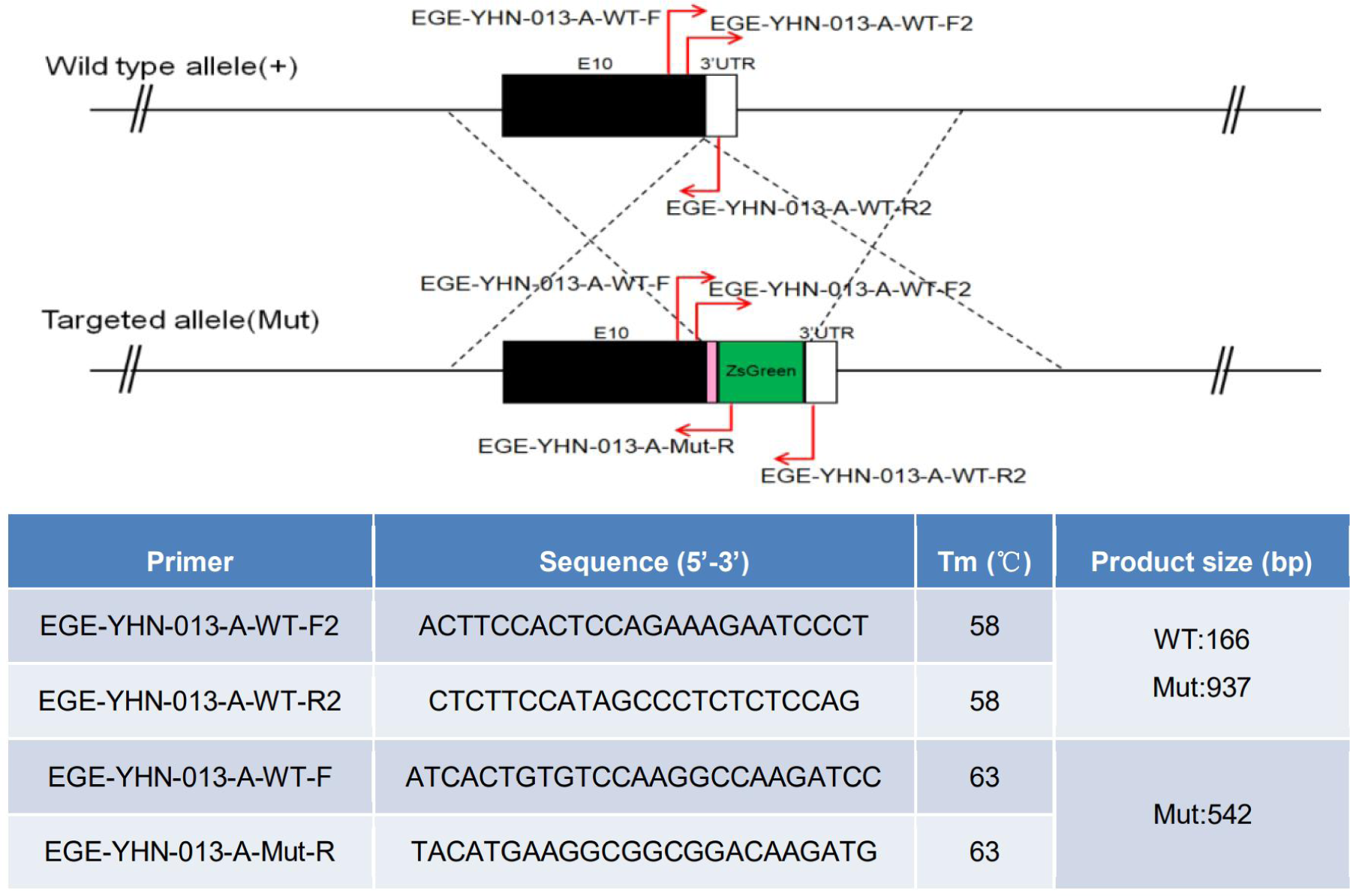

## Supplementary materials 2

### 1. The primary antibodies used for IFs

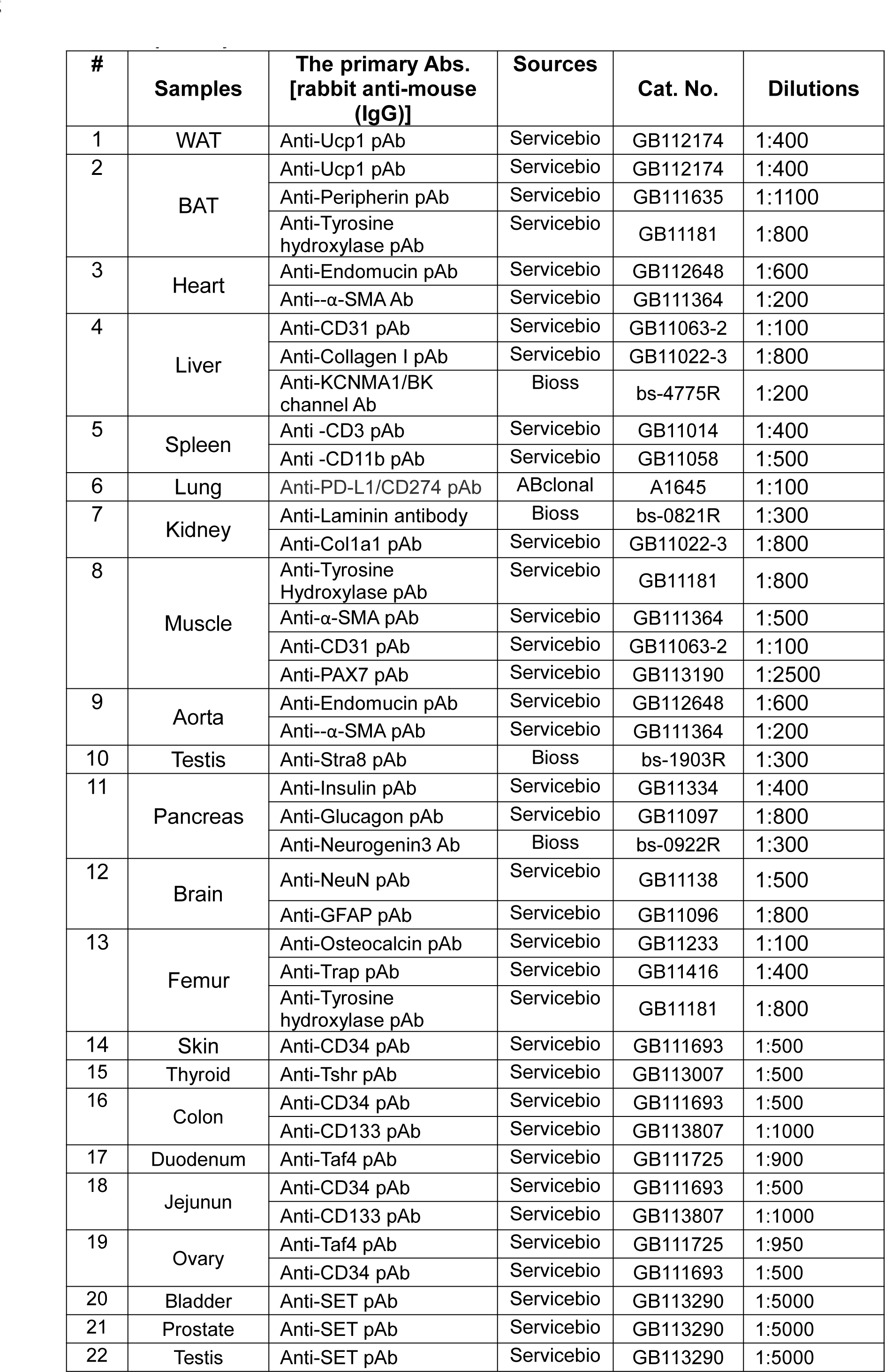

### 2. The sequences of sense and antisense probes for FISH in testis

#### 1) For the sense probe

5’-ATCACTGGCTGTGTCATTGCTCTAA-3’

#### 2) For antisense probes: a mixture of the following 5 probes used for FISH

5’-TTAGAGCAATGACACAGCCAGTGAT-3’

5’-CACCTTGCTATCTTGGCAGAGGAAG-3’

5’-AGCACAAATCTCAGTTCAATGGCGT-3’

5’-CAAGTGTTTAATGCCTGTGTTGGAT-3’

5’-GCAGGGAATAGACCTTTGTCCTTGA-3’

## Supplementary Data 1

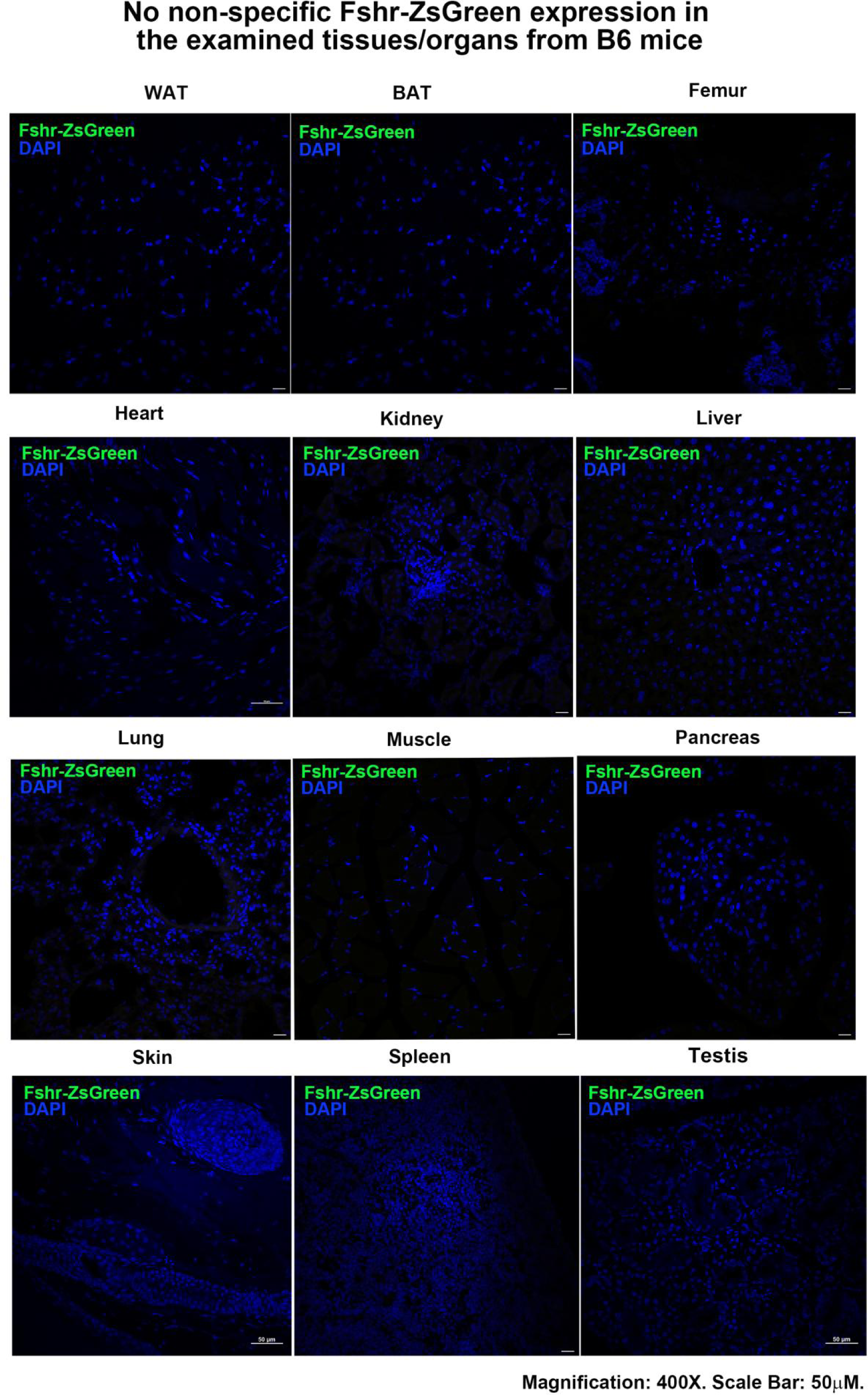

## Supplementary Data 2

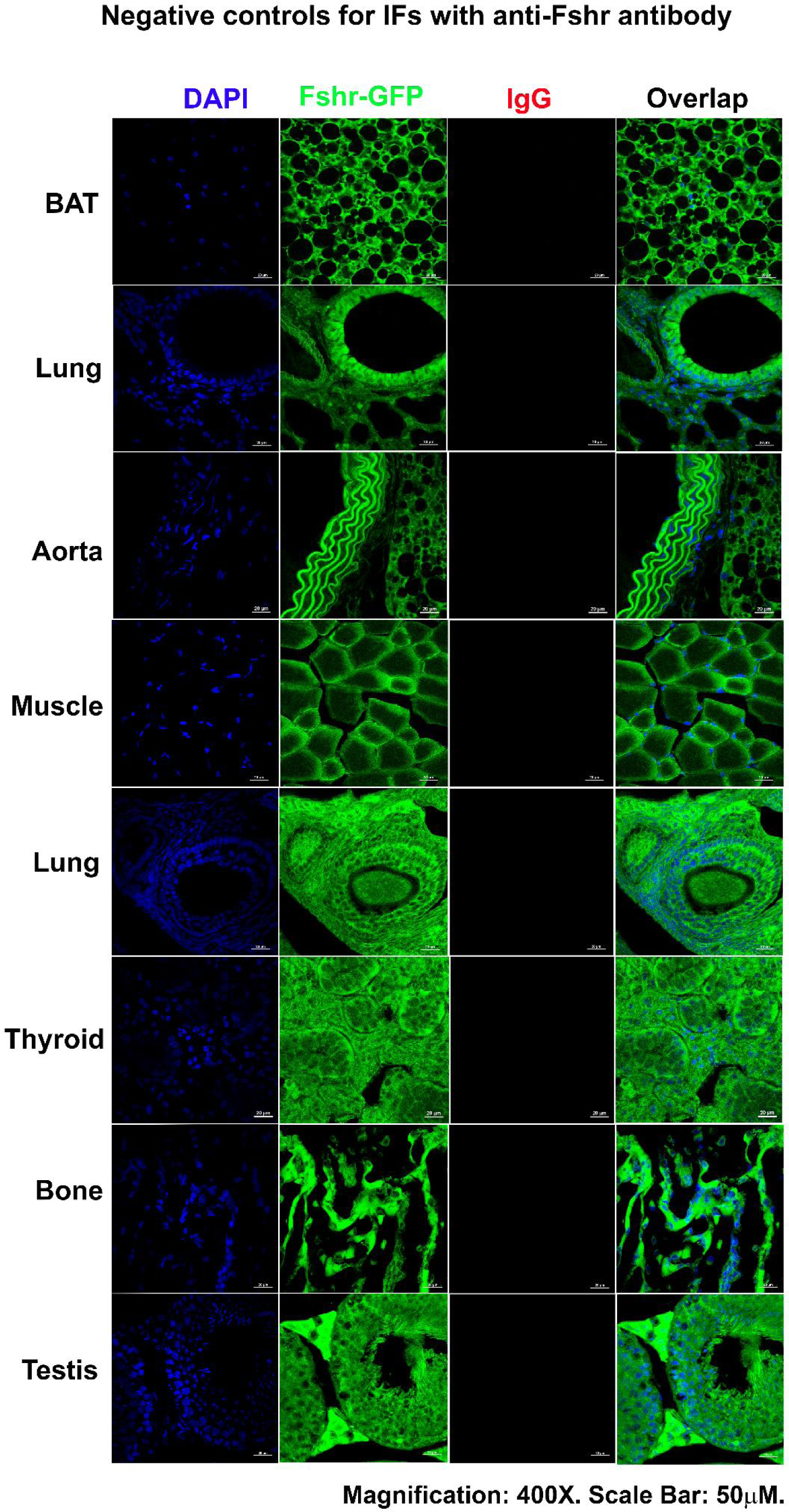

## Supplementary Data 3

### FSHR expression detected by scRNA-seq from 4 public single cell portals

#### 1. DISCO (immunesinglecell.org/genepage/FSHR)

##### (1) Testis and ovary

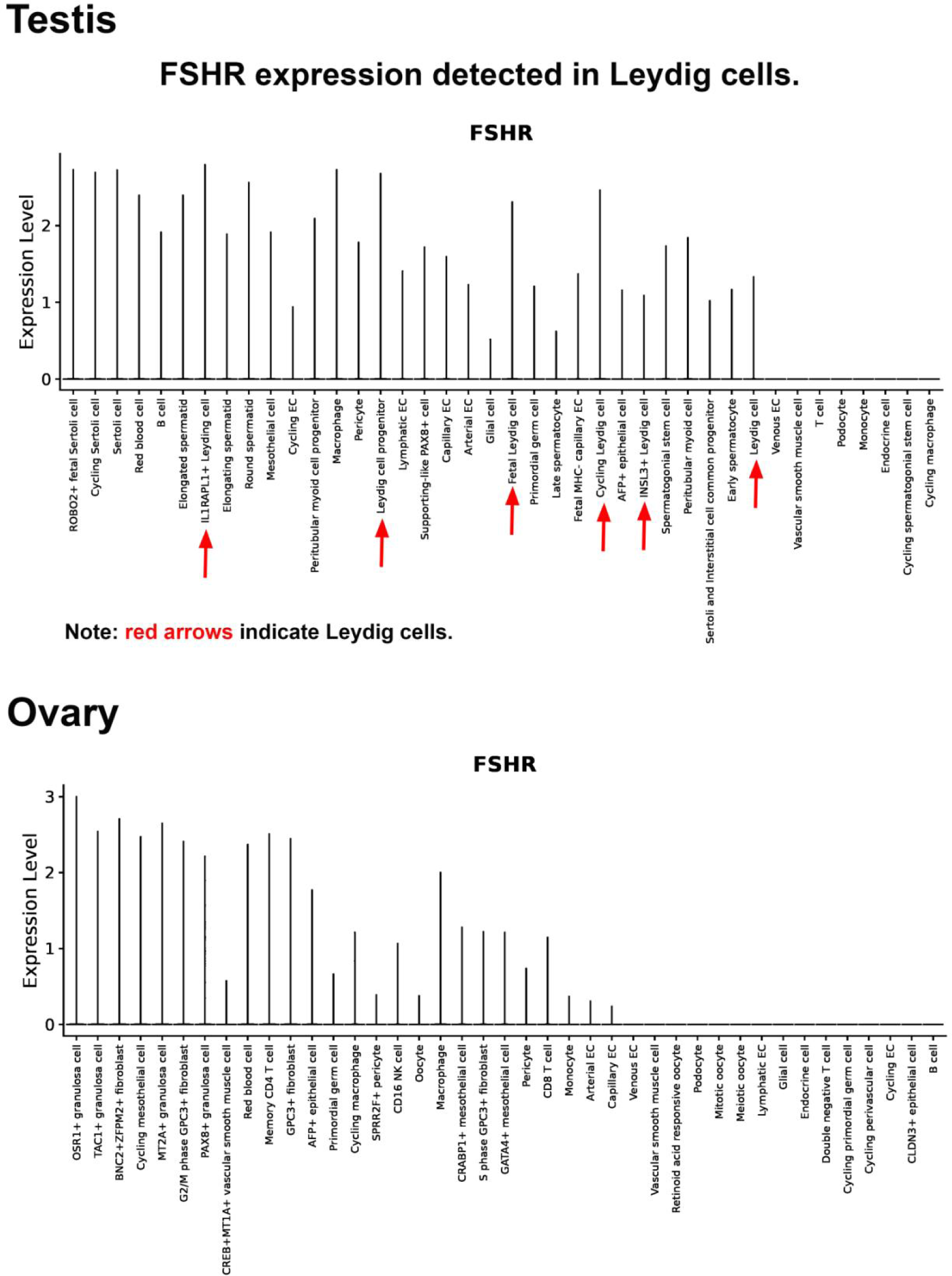

##### (2) Bone marrow and brain

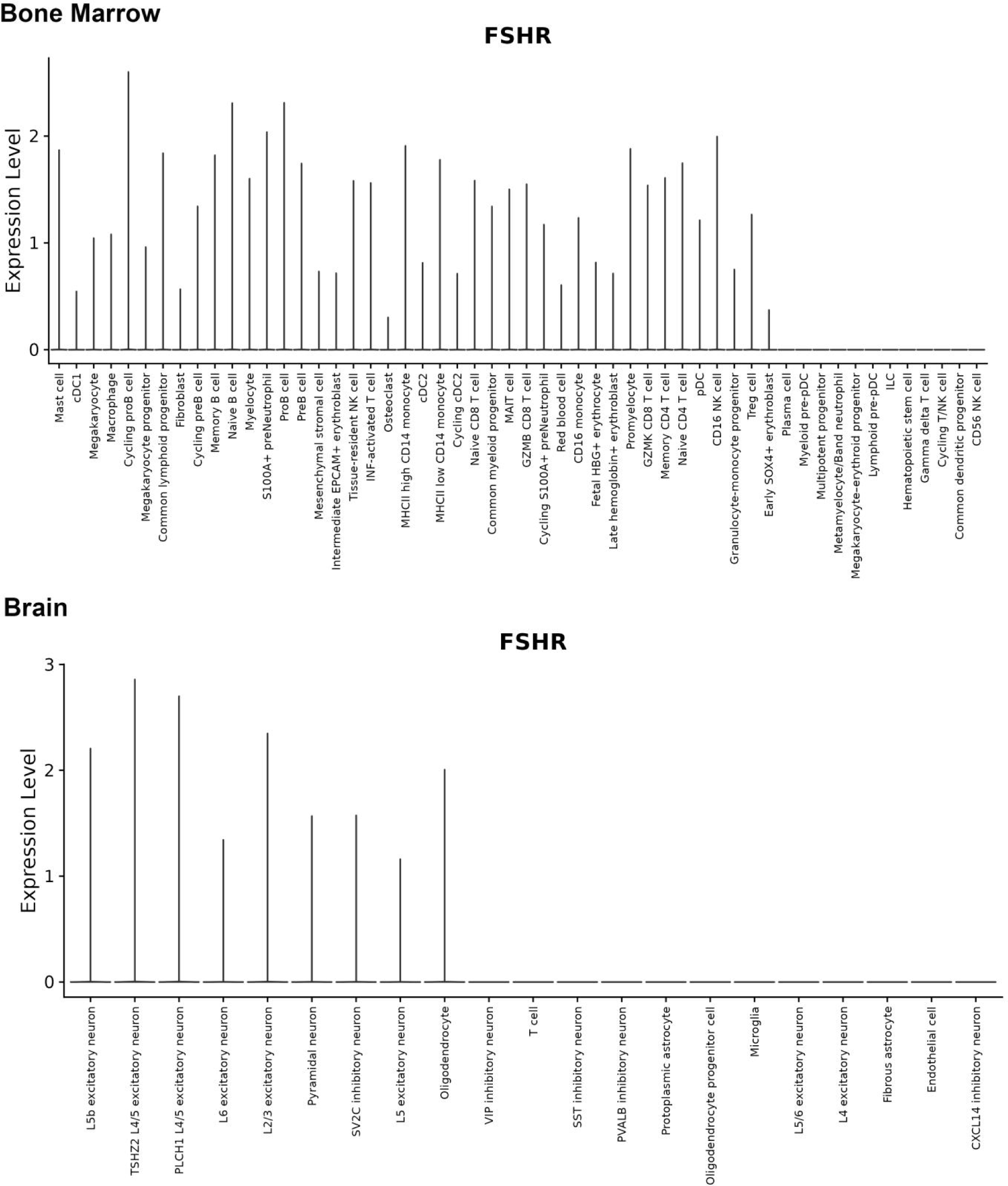

##### (3) Breast and Heart

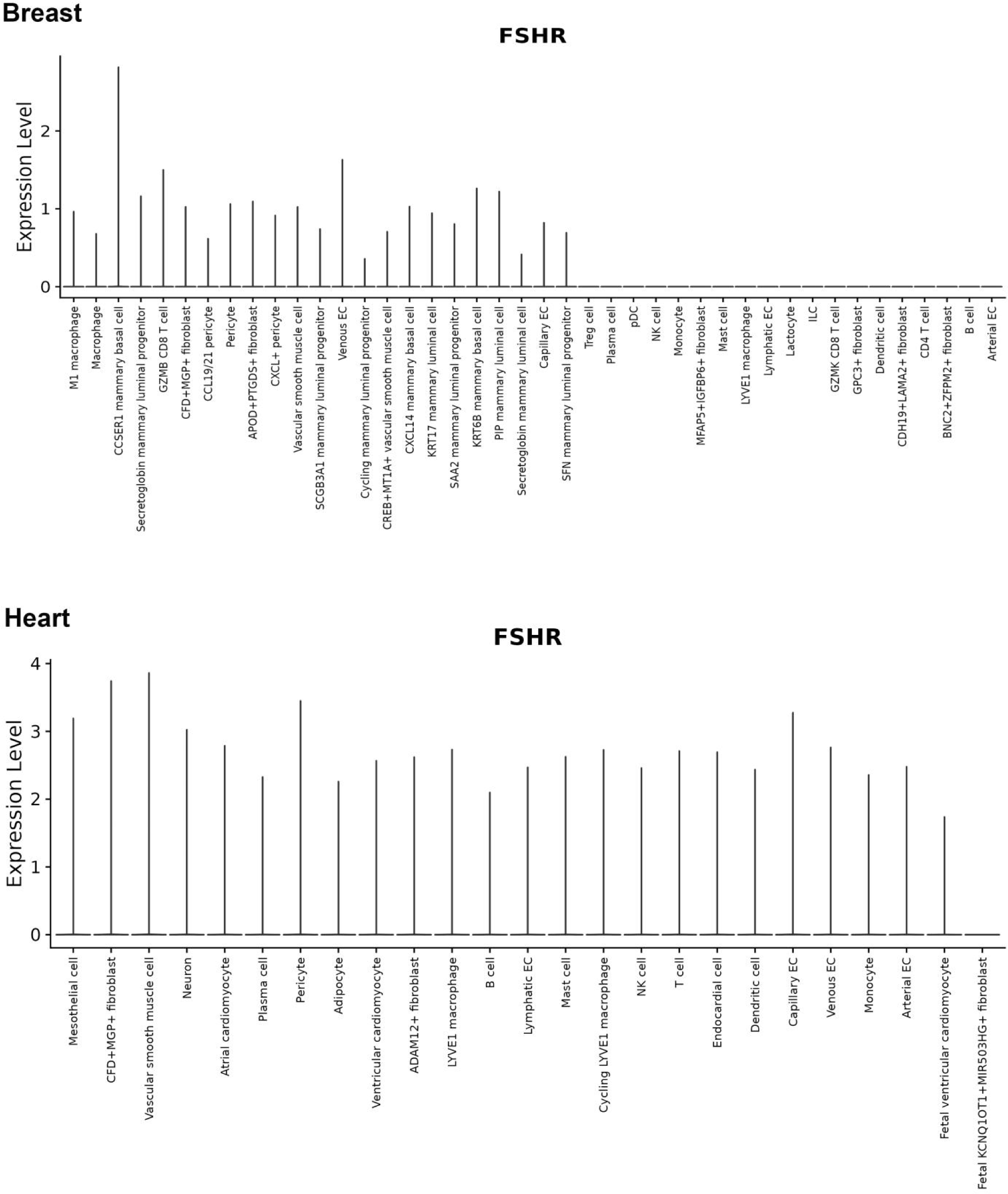

##### (4) Liver and Lung

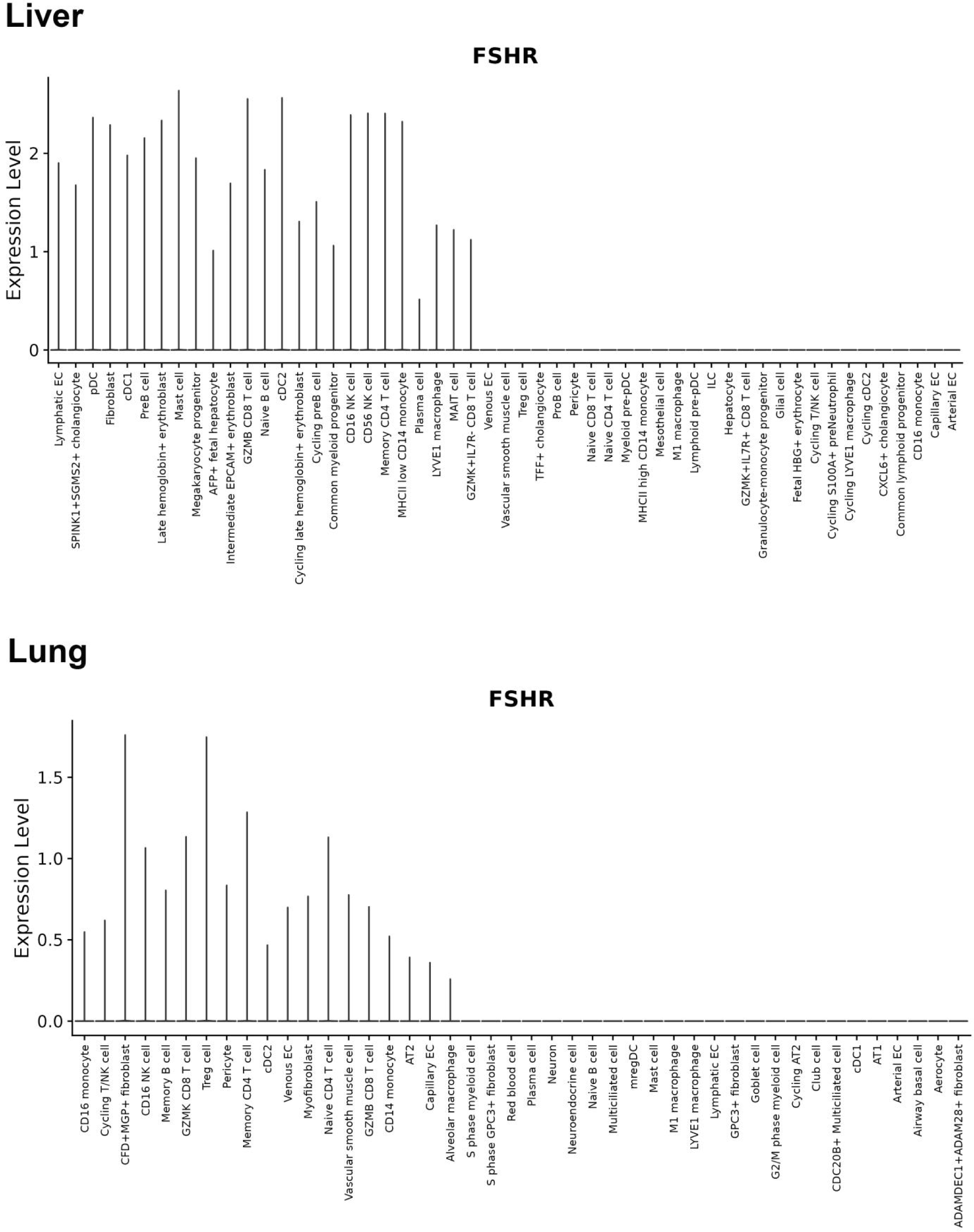

##### (5) Skeletal muscle and thymus

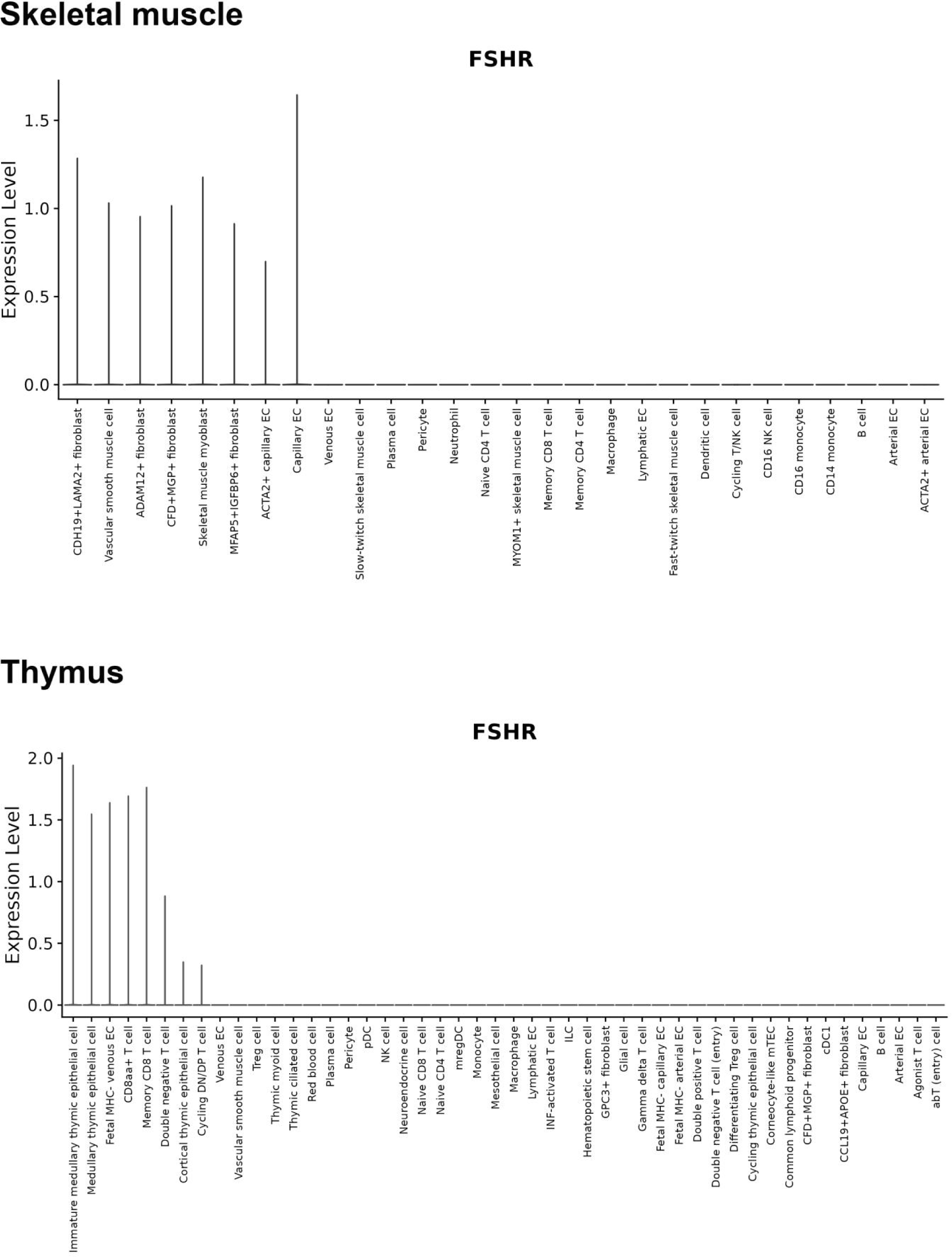

#### 2. GTEx: https://www.gtexportal.org/home/gene/FSHR#singleCell

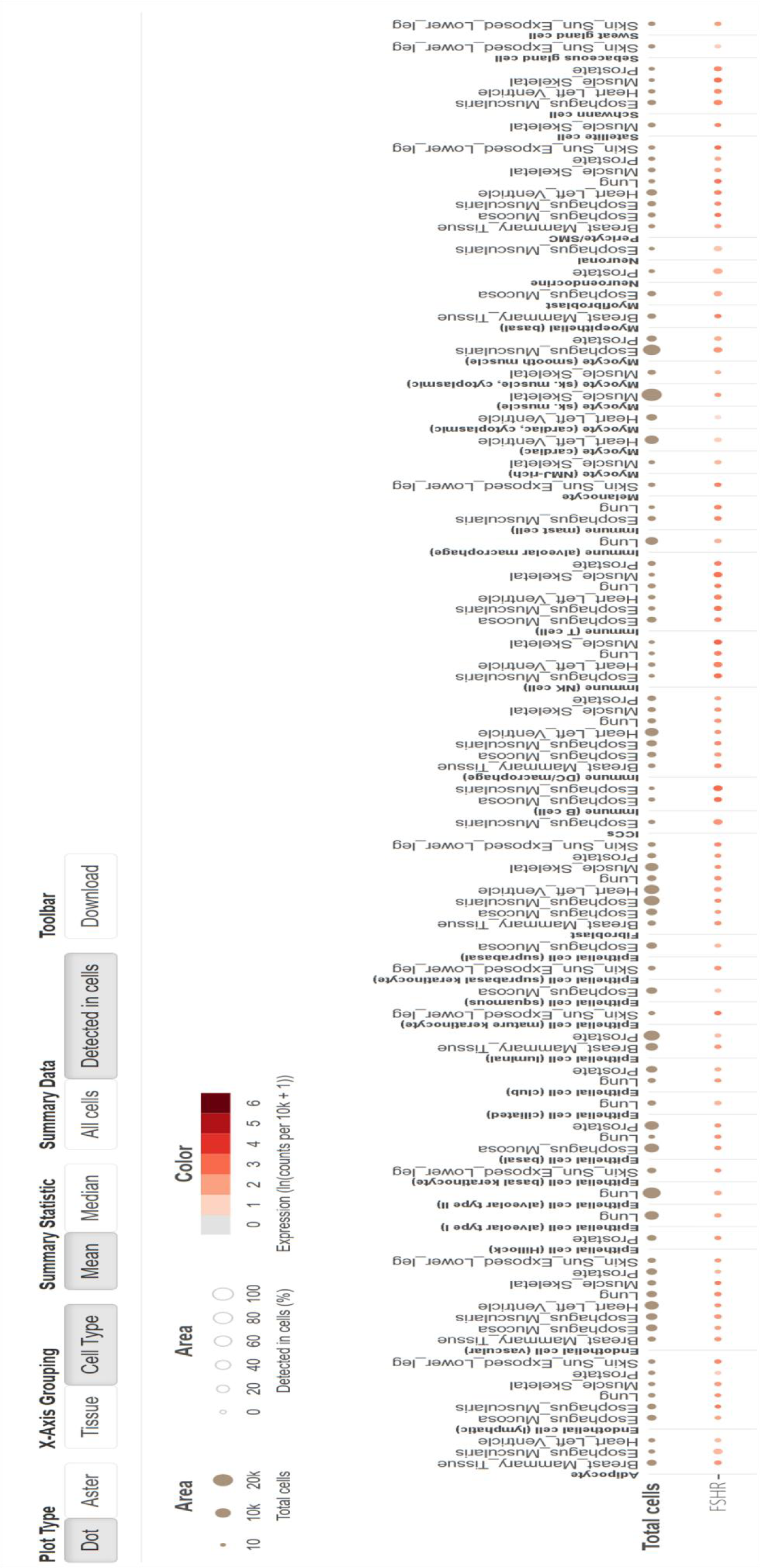

#### 3. BioGPS: (http://biogps.org/#goto=searchresult)

##### (1) Human: http://biogps.org/#goto=genereport&id=2492

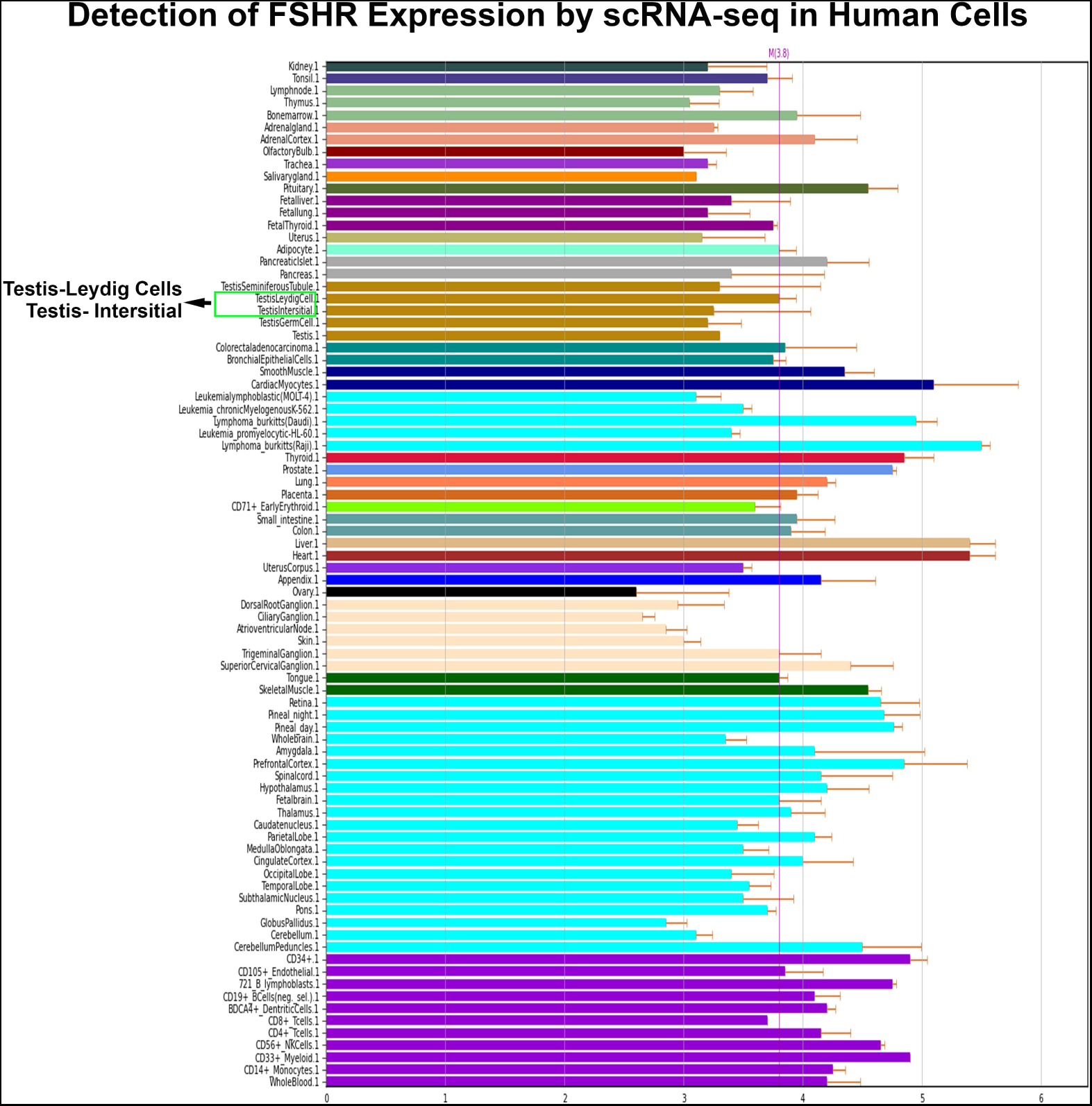

##### (2) Rat: http://ds.biogps.org/?dataset=GSE952&gene=25449

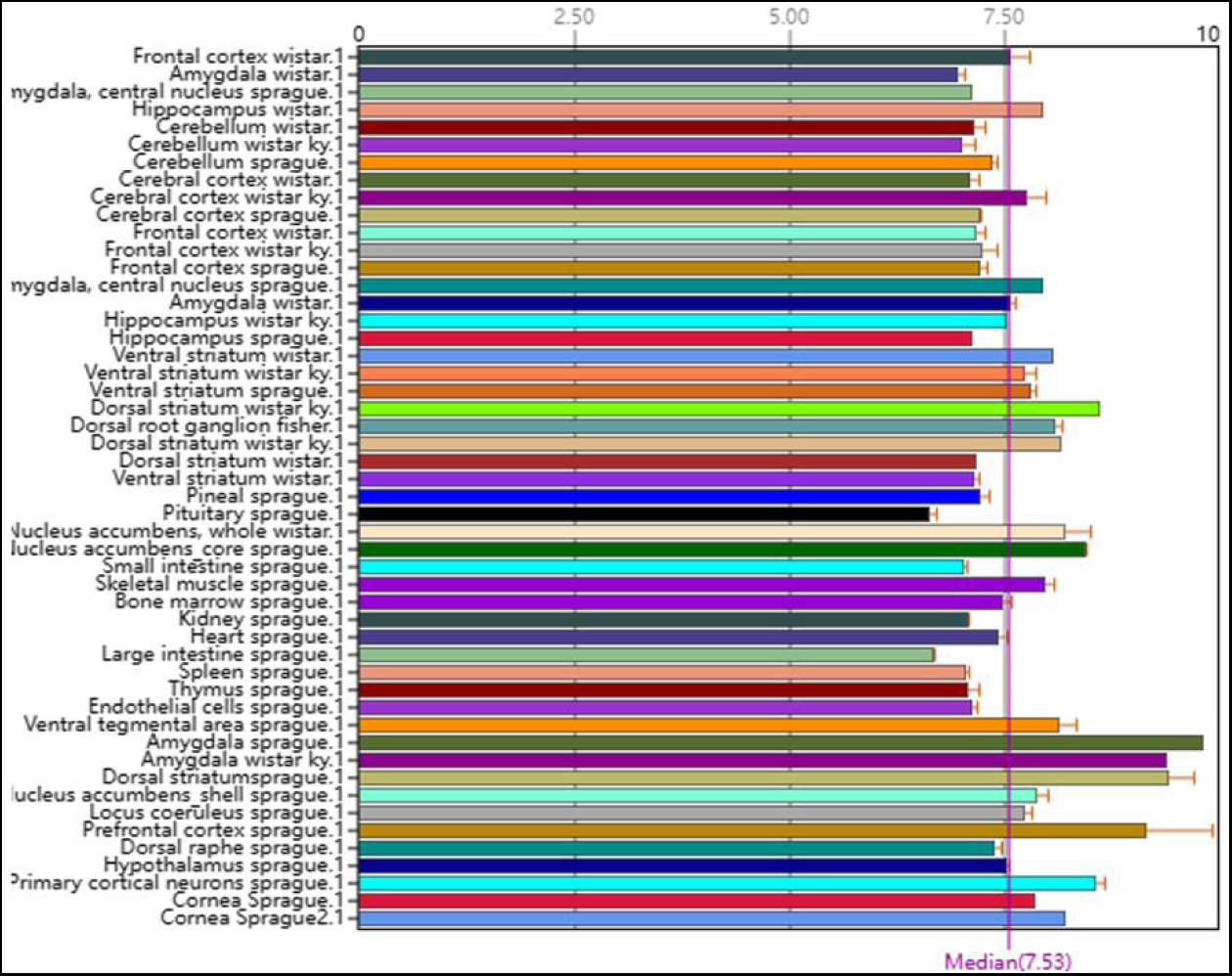

#### 4. Single cell portal: https://singlecell.broadinstitute.org/single_cell

##### (1) Adipose

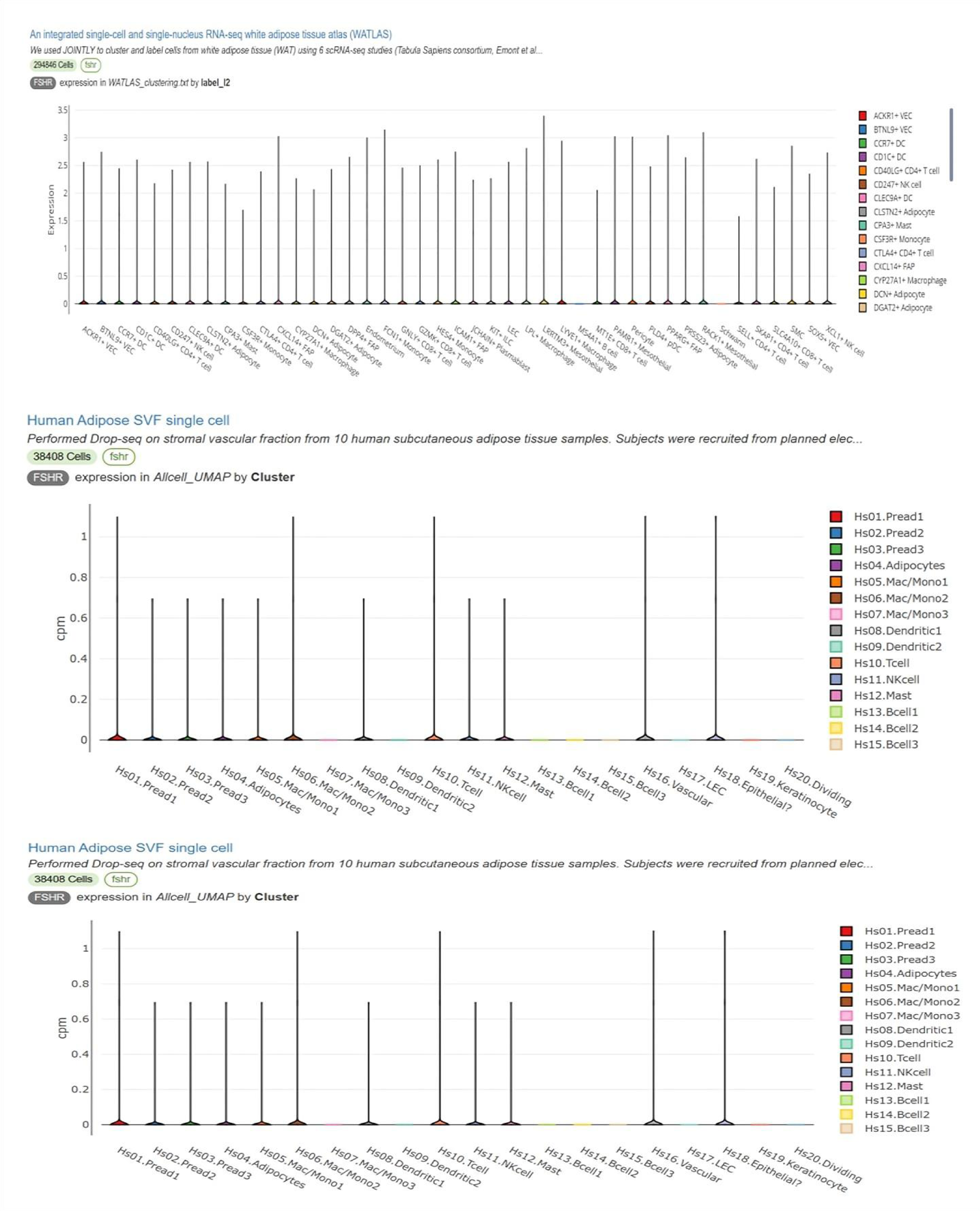

##### (2) Brain

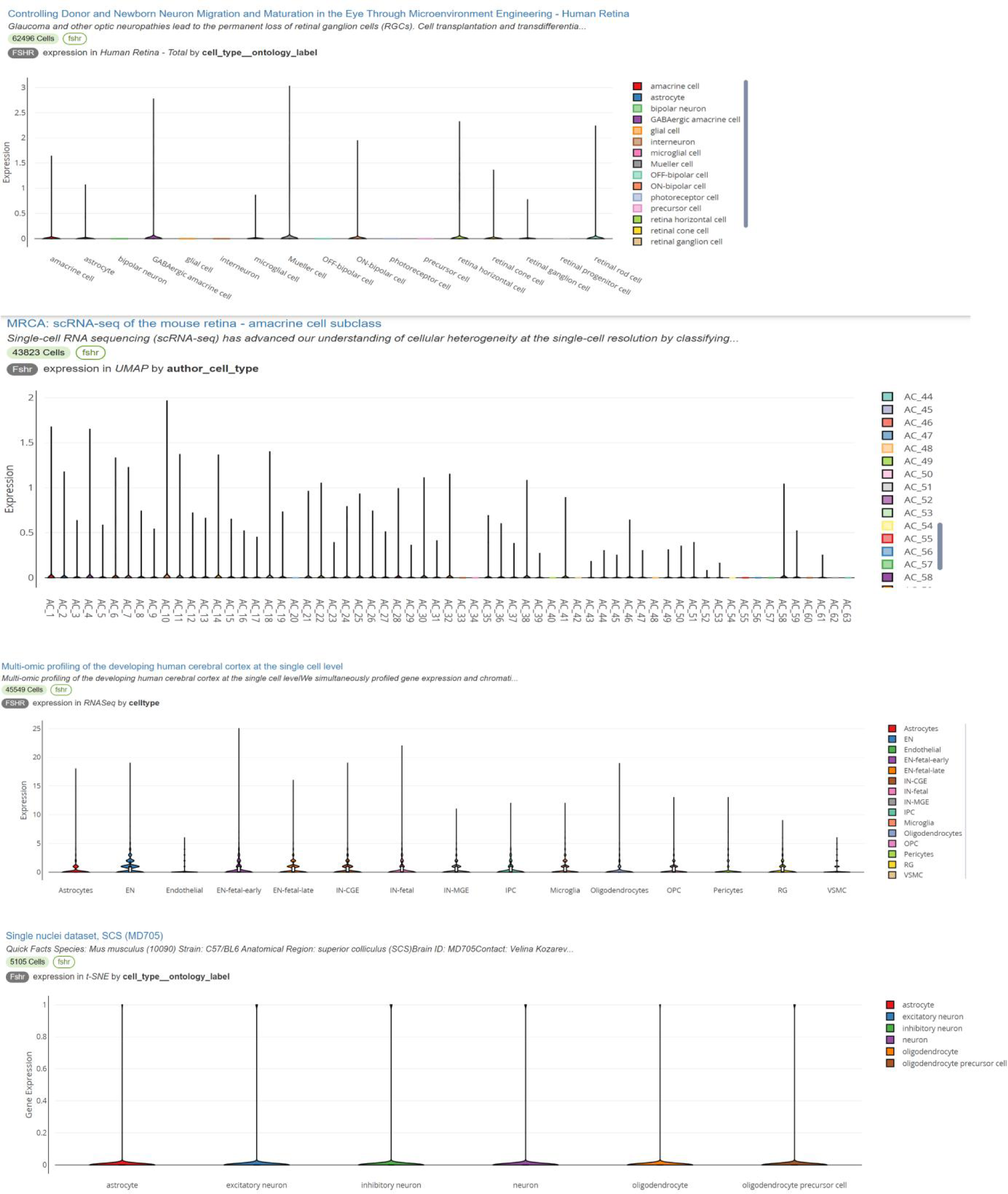

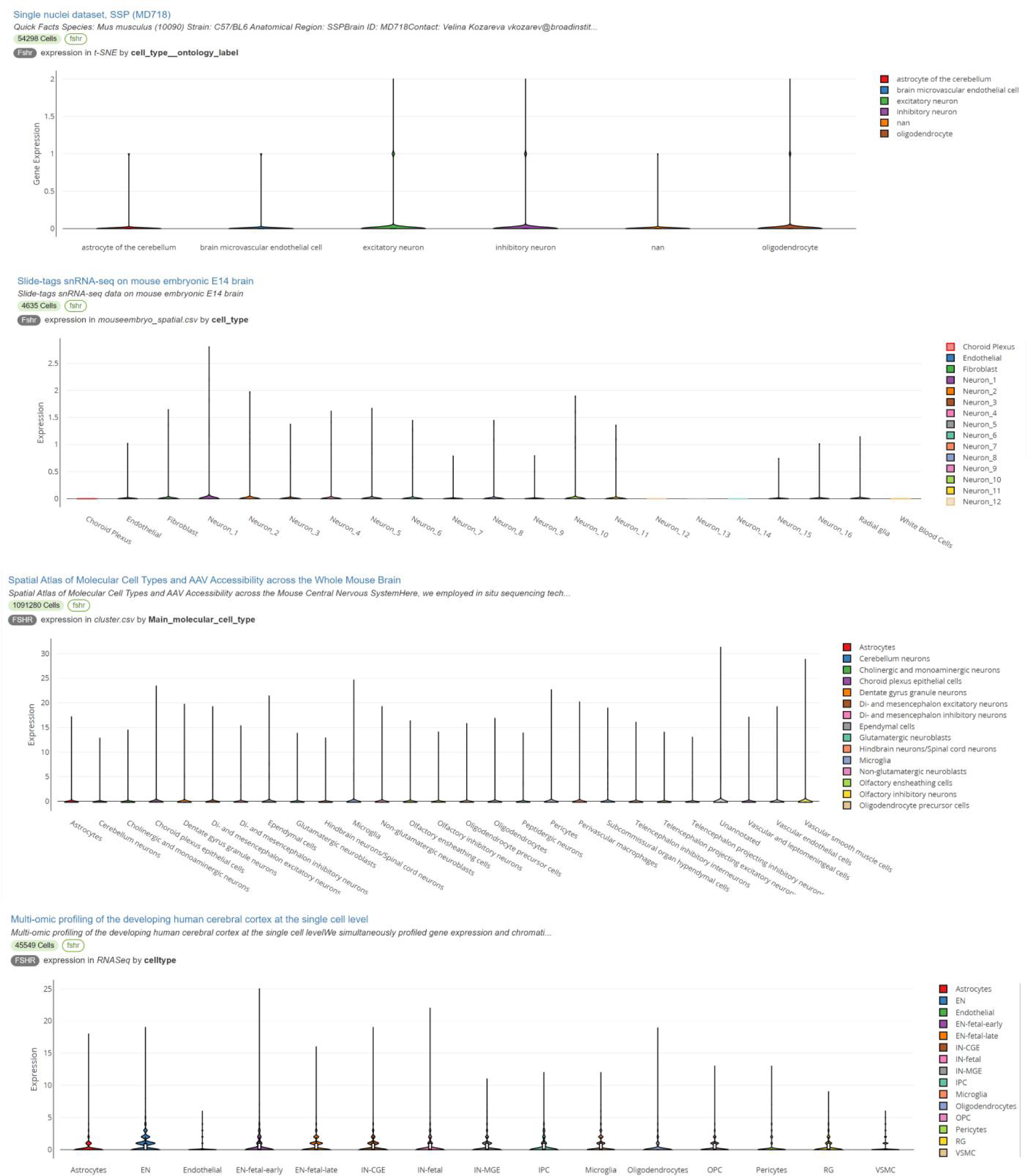

##### (3) Kidney

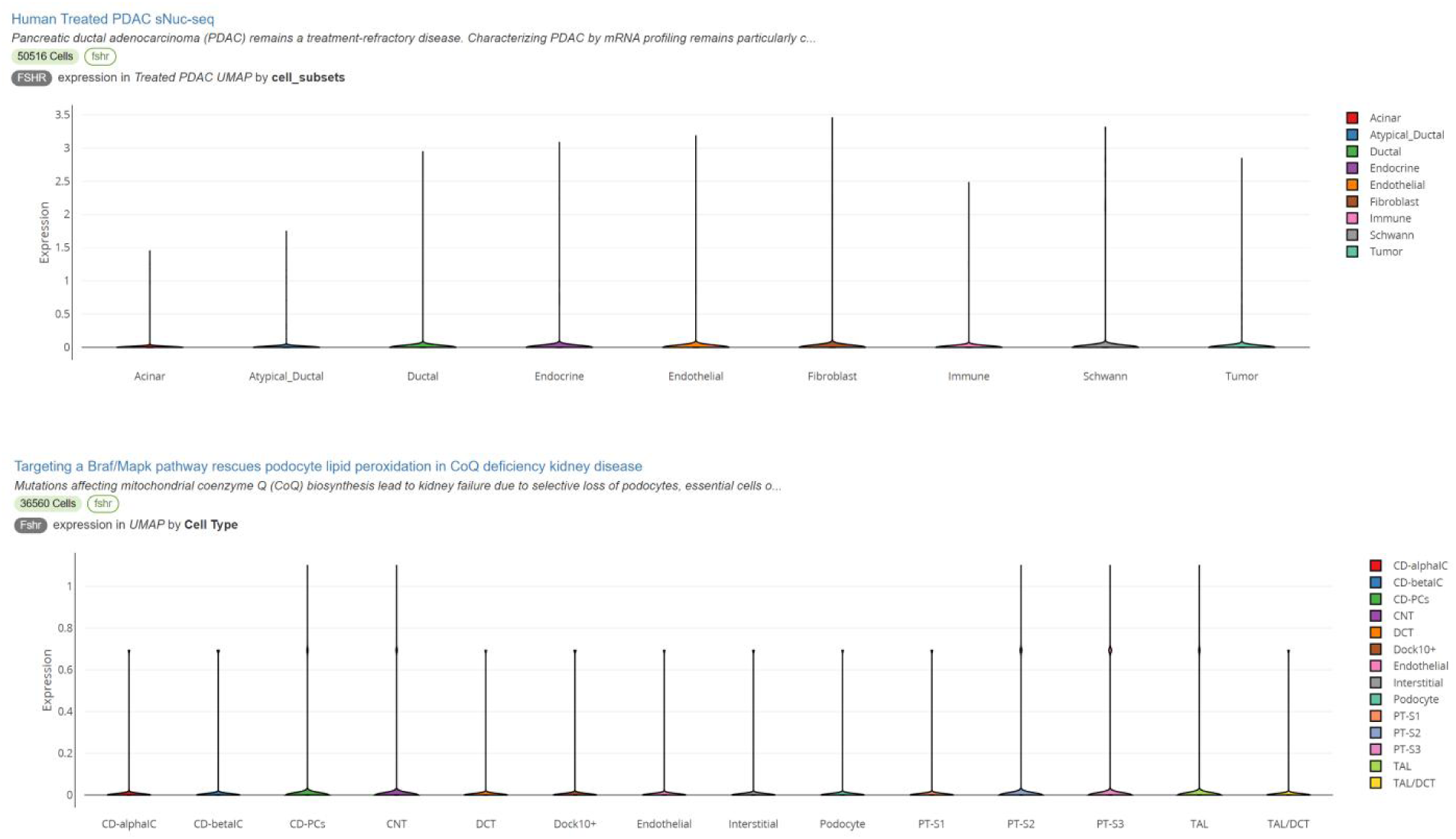

##### (4) Liver

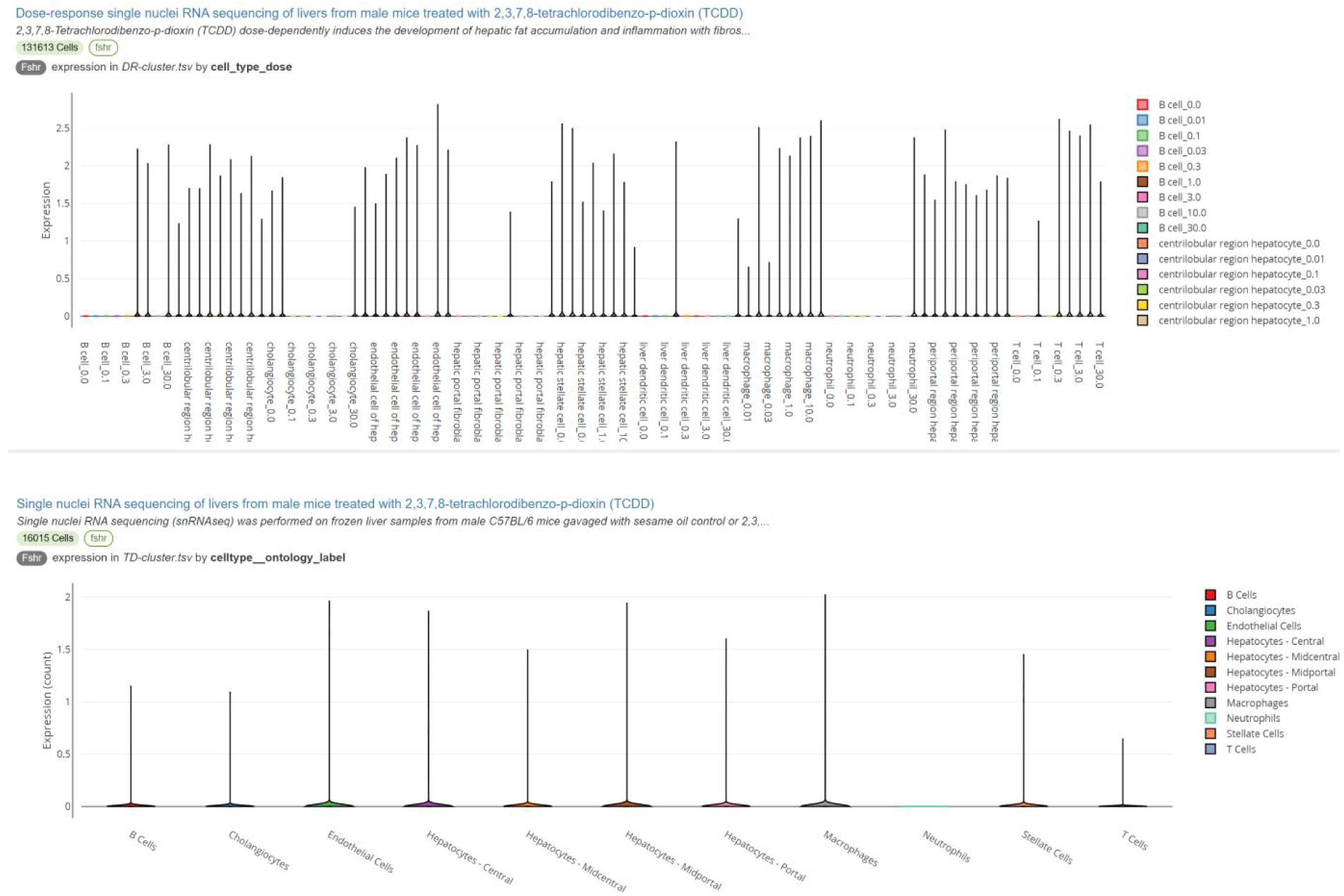

##### (5) Lung

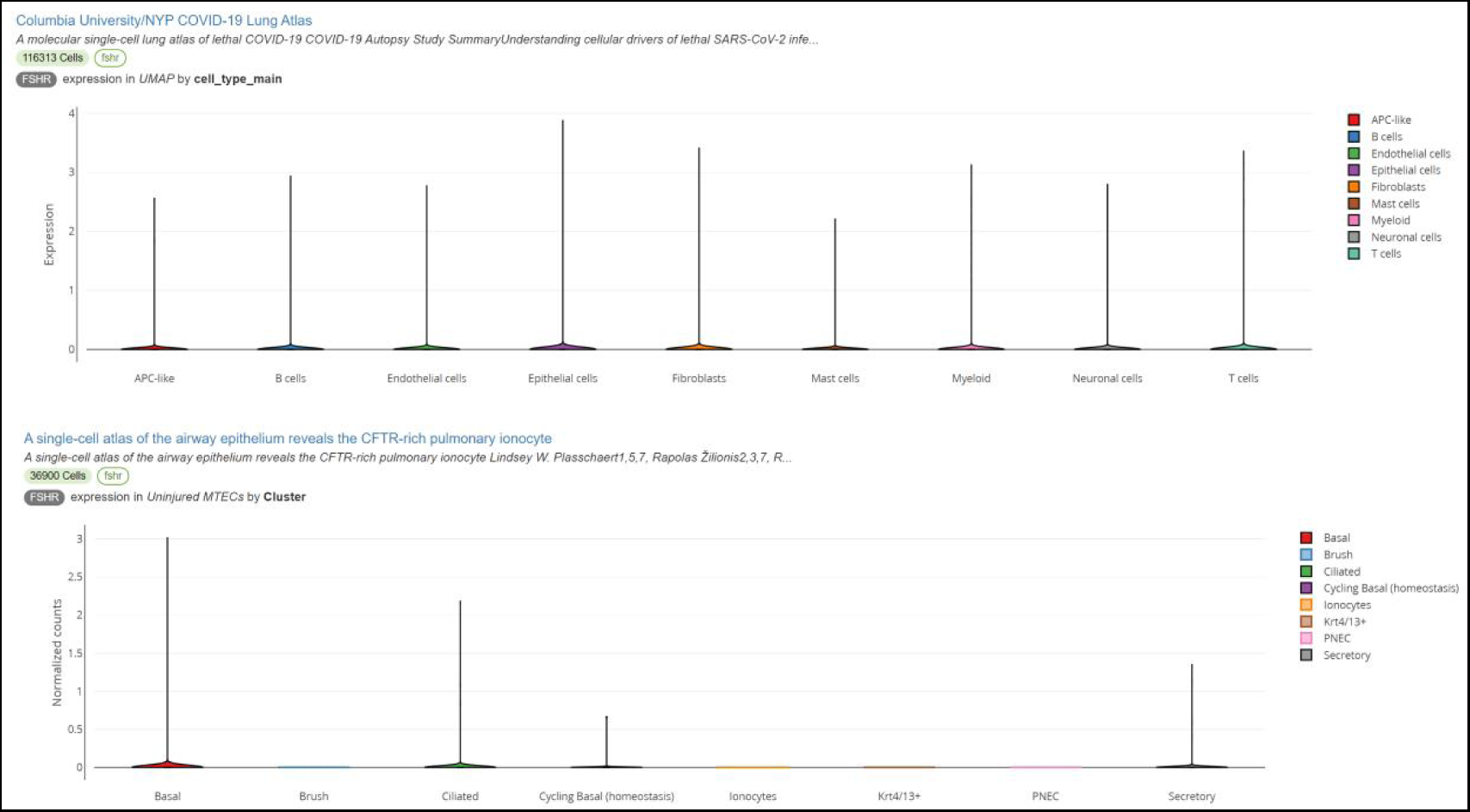

##### (6) Ovary and Pancreas

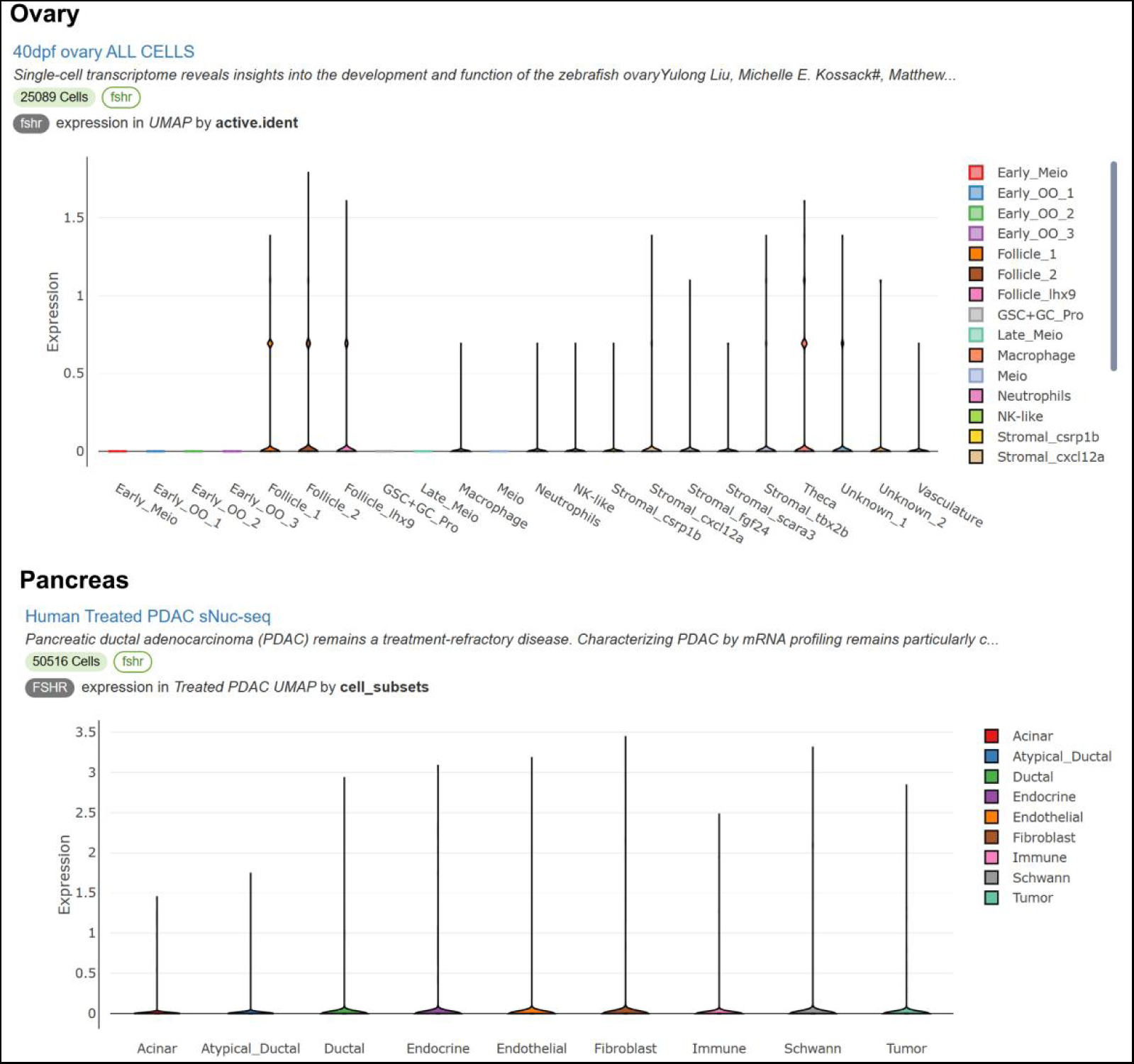

##### (7) Heart and vascular system

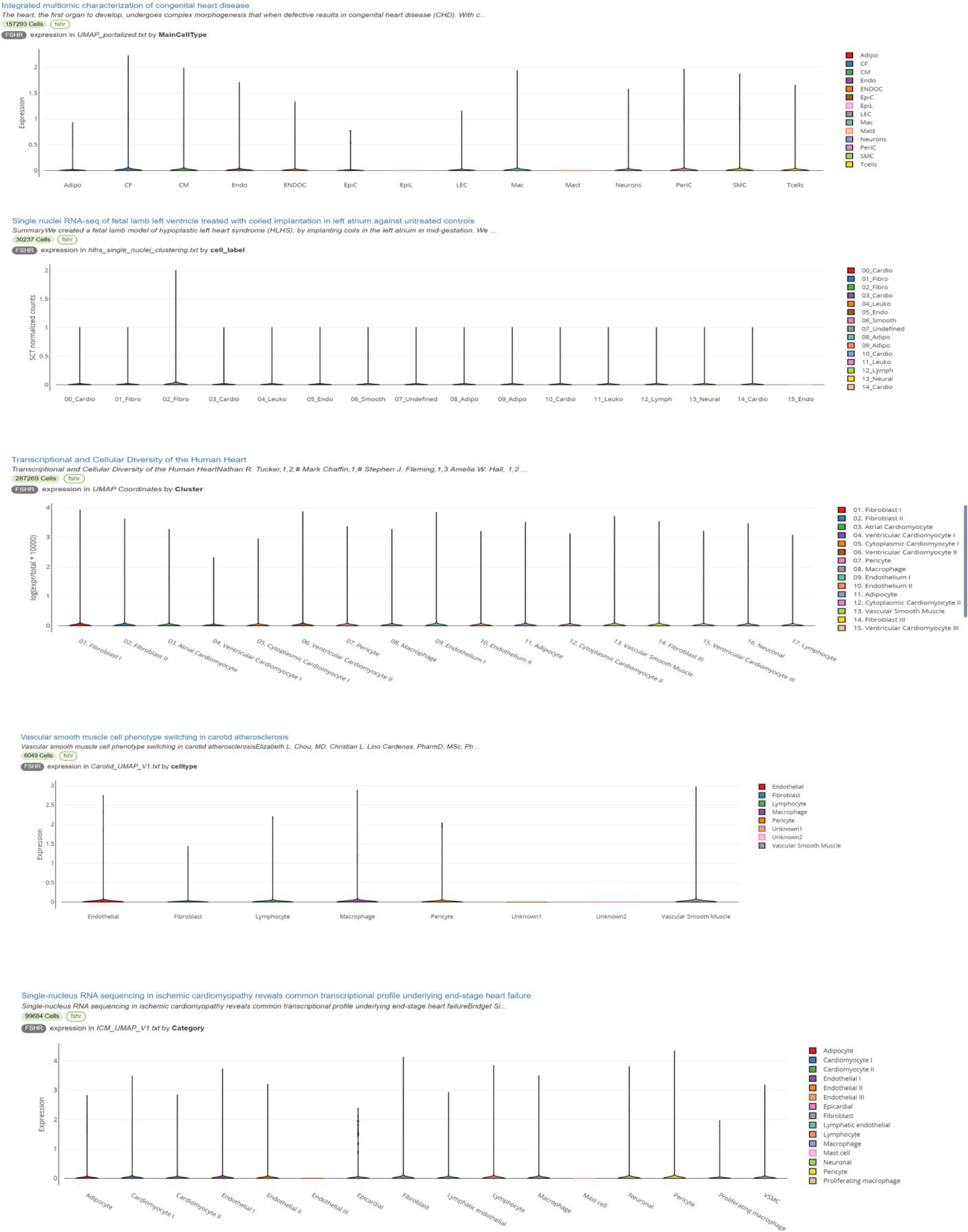

#### 5. FSHR/Fshr expression in Leydig cells of human and mouse males from Chan Zuckerberg CELLxGENE Discover

(1) Human male: https://cellxgene.cziscience.com/e/535e9336-2d8d-43c3-944d-bcbebe20df8a.cxg/.

(2) Mouse male: https://cellxgene.cziscience.com/e/a13bda79-9134-46c9-9ed1-a2858be9aafe.cxg/.

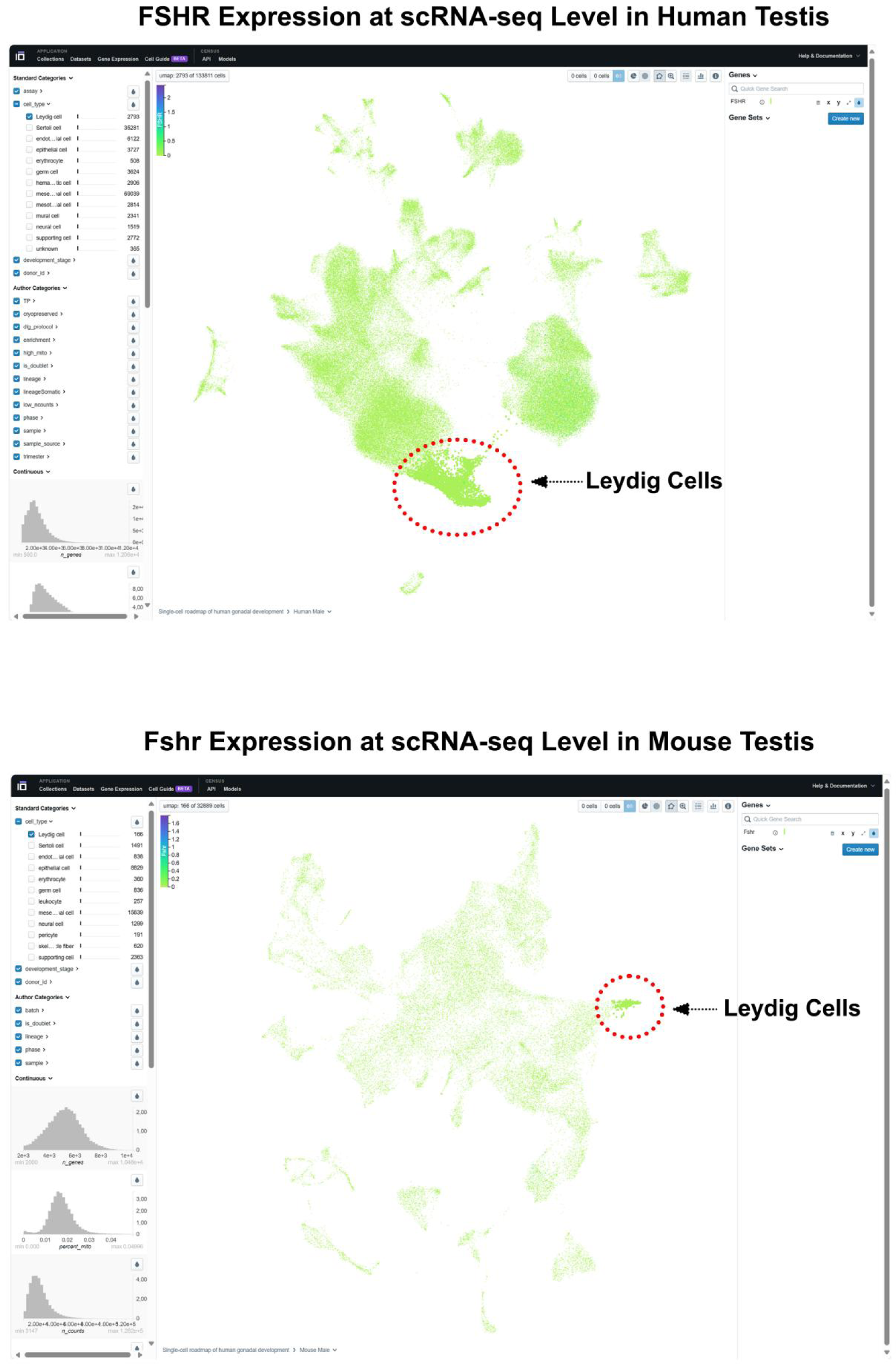

(3) FSHR in human organs: (https://cellxgene.cziscience.com/gene-expression)

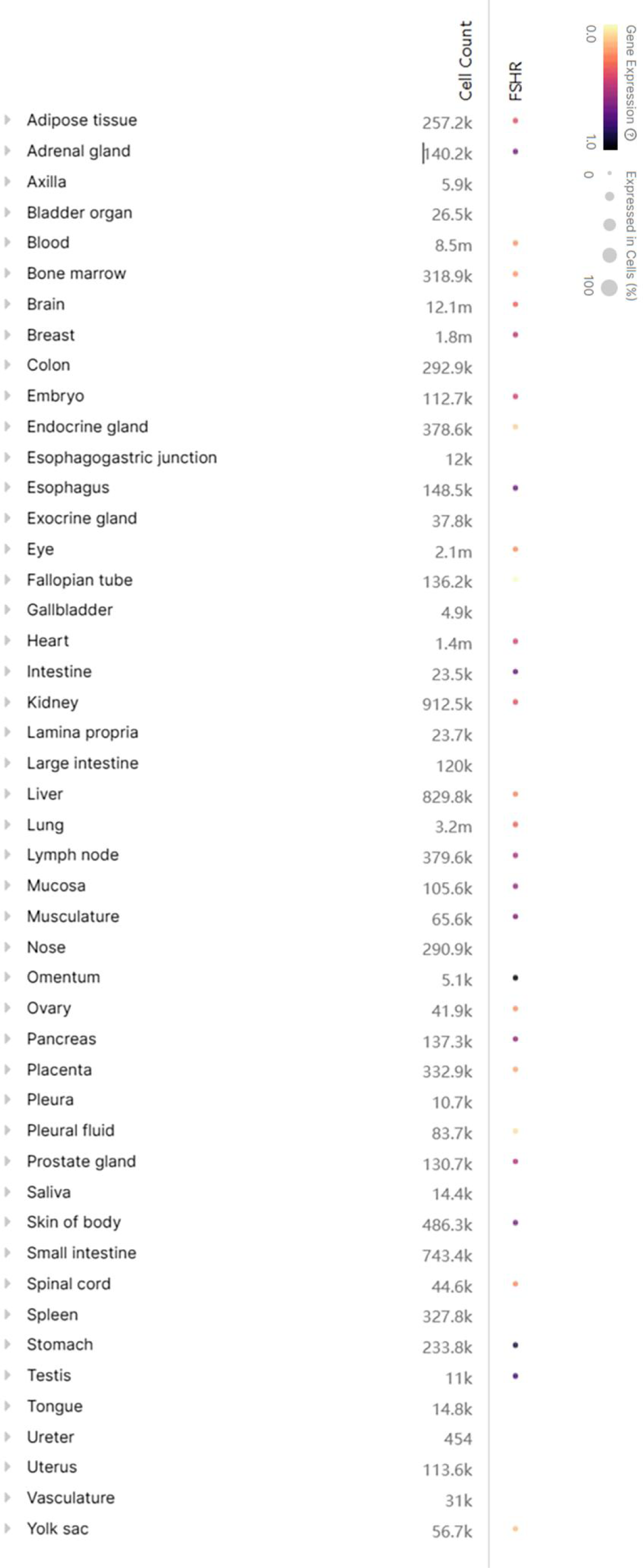

